# Identifying and classifying shared selective sweeps from multilocus data

**DOI:** 10.1101/446005

**Authors:** Alexandre M. Harris, Michael DeGiorgio

## Abstract

Positive selection causes beneficial alleles to rise to high frequency, resulting in a selective sweep of the diversity surrounding the selected sites. Accordingly, the signature of a selective sweep in an ancestral population may still remain in its descendants. Identifying signatures of selection in the ancestor that are shared among its descendants is important to contextualize the timing of a sweep, but few methods exist for this purpose. We introduce the statistic SS-H12, which can identify genomic regions under shared positive selection across populations and is based on the theory of the expected haplotype homozygosity statistic H12, which detects recent hard and soft sweeps from the presence of high-frequency haplotypes. SS-H12, is distinct from other statistics that detect shared sweeps because it requires a minimum of only two populations, and properly identifies and differentiates between independent convergent sweeps and true ancestral sweeps, with high power and robustness to a variety of demographic models. Furthermore, we can apply SS-H12 in conjunction with the ratio of a different set of expected haplotype homozygosity statistics to further classify identified shared sweeps as hard or soft. Finally, we identified both previously-reported and novel shared sweep candidates from whole-genome sequences of global human populations. Previously-reported candidates include the well-characterized ancestral sweeps at *LCT* and *SLC24A5* in Indo-European populations, as well as *GPHN* worldwide. Novel candidates include an ancestral sweep at *RGS18* in sub-Saharan African populations involved in regulating the platelet response and implicated in sudden cardiac death, and a convergent sweep at *C2CD5* between European and East Asian populations that may explain their different insulin responses.Introduction

## Introduction

Alleles under positive selection increase in frequency in a population toward fixation, causing nearby linked neutral variants to also rise to high frequency. This process results in selective sweeps of the diversity surrounding the sites of selection, which can be hard or soft [Hermisson and Pennings, 2005, Pennings and Hermisson, 2006a,b, Hermisson and Pennings, 2017]. Under hard sweeps, beneficial alleles exist on a single haplotypic background at the time of selection, and a single haplotype rises to high frequency with the selected variants. In contrast, soft sweeps occur when beneficial alleles are present on multiple haplotypic backgrounds, each of which increases in frequency with the selected variants. Thus, individuals carrying the selected alleles do not all share a common haplotypic background. The signature of a selective sweep, hard or soft, is characterized by elevated levels of linkage disequilibrium (LD) on either side of the beneficial mutation, and elevated expected haplotype homozygosity [Maynard Smith and Haigh, 1974, Sabeti et al., 2002, Schweinsberg and Durrett, 2005]. Thus, the signature of a selective sweep decays with distance from the selected site as mutation and recombination erode tracts of sequence identity produced by the sweep, returning expected haplotype homozygosity and LD to their neutral levels [Messer and Petrov, 2013].

Various approaches exist to detect signatures of selective sweeps in single populations, but few methods can identify sweep regions shared across populations, and these methods primarily rely on allele frequency data as input. Existing methods to identify shared sweeps [Bonhomme et al., 2010, Fariello et al., 2013, Racimo, 2016, Librado et al., 2017, Peyrégne et al., 2017, Cheng et al., 2017, Johnson and Voight, 2018] leverage the observation that study populations sharing similar patterns of genetic diversity at a putative site under selection descend from a common ancestor in which the sweep occurred. Such approaches therefore infer a sweep ancestral to the study populations from what may be coincidental (*i.e*., independent) signals. Moreover, many of these methods require data from at least one reference population in addition to the study populations, and of these, most may be misled by sweeps in their set of reference populations. These constraints may therefore impede the application of these methods to study systems that do not fit their model assumptions or data requirements.

Identifying sweeps common to multiple populations provides an important layer of context that specifies the branch of a genealogy on which a sweep is likely to have occurred. In this way, the timing and types of pressures that contributed to particular signals among sampled populations can become clearer. For example, identifying sweeps that are shared ancestrally among all populations within a species highlights the selective events that contributed to their most important modern phenotypes. On a smaller scale, methods to identify shared sweeps can be leveraged to distinguish signatures of local adaptation in particular populations [Librado and Orlando, 2018]. In contrast, single-population tests would provide little information about the timing and therefore relative importance of detected sweeps. More generally, tests tailored to the detection of sweeps within samples drawn from multiple populations are likely to have higher power to detect such events than are tests that do not account for sample complexity [Bonhomme et al., 2010, Fariello et al., 2013], underscoring the usefulness of multi-population approaches.

Here, we develop an expected haplotype homozygosity-based statistic, denoted SS-H12, that addresses the constraints of other methods. SS-H12 detects shared selective sweeps from a minimum of two sampled populations using haplotype data (Figure 1), and classifies sweep candidates as either ancestral (shared through common ancestry) or convergent (occurring independently). We base our approach on the theory of H12 [Garud et al., 2015, Garud and Rosenberg, 2015], a summary statistic that measures expected homozygosity in haplotype data from a single population. This single-population statistic has high power to detect recent selective sweeps, and identifies hard and soft sweeps with similar power due to its unique formulation. For a genomic window in which there are *I* distinct haplotypes, H12 [Garud et al., 2015] is defined as

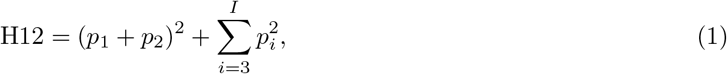

where *p_i_* is the frequency of the *i*th most frequent haplotype, with *p*_1_ ≥ *p*_2_ ≥ ⋯ ≥ *p_I_*. The two most common haplotype frequencies are pooled into a single value to reflect the presence of at least two high-frequency haplotypes in the population under a soft sweep. Meanwhile, the squares of the remaining haplotype frequencies are summed as in standard computations of expected homozygosity, reflecting the probability of drawing two copies of the third through *I*th most frequent haplotypes at random from the population. Thus, H12 yields similar values for hard and soft sweeps. In addition, the framework of the single-population statistic distinguishes hard and soft sweeps using the H2/H1 ratio [Garud et al., 2015, Garud and Rosenberg, 2015], where 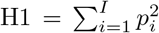 is the expected haplotype homozygosity, and where 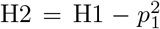 is the expected haplotype homozygosity omitting the most frequent haplotype. The value of H2/H1 is small under hard sweeps, because the second through *I*th frequencies are small, as the beneficial alleles exist only on a single haplotypic background. Accordingly, H2/H1 is larger for soft sweeps [Garud et al., 2015], and can therefore be used to classify sweeps as hard or soft, conditioning on an elevated value of H12.

**Figure 1:**
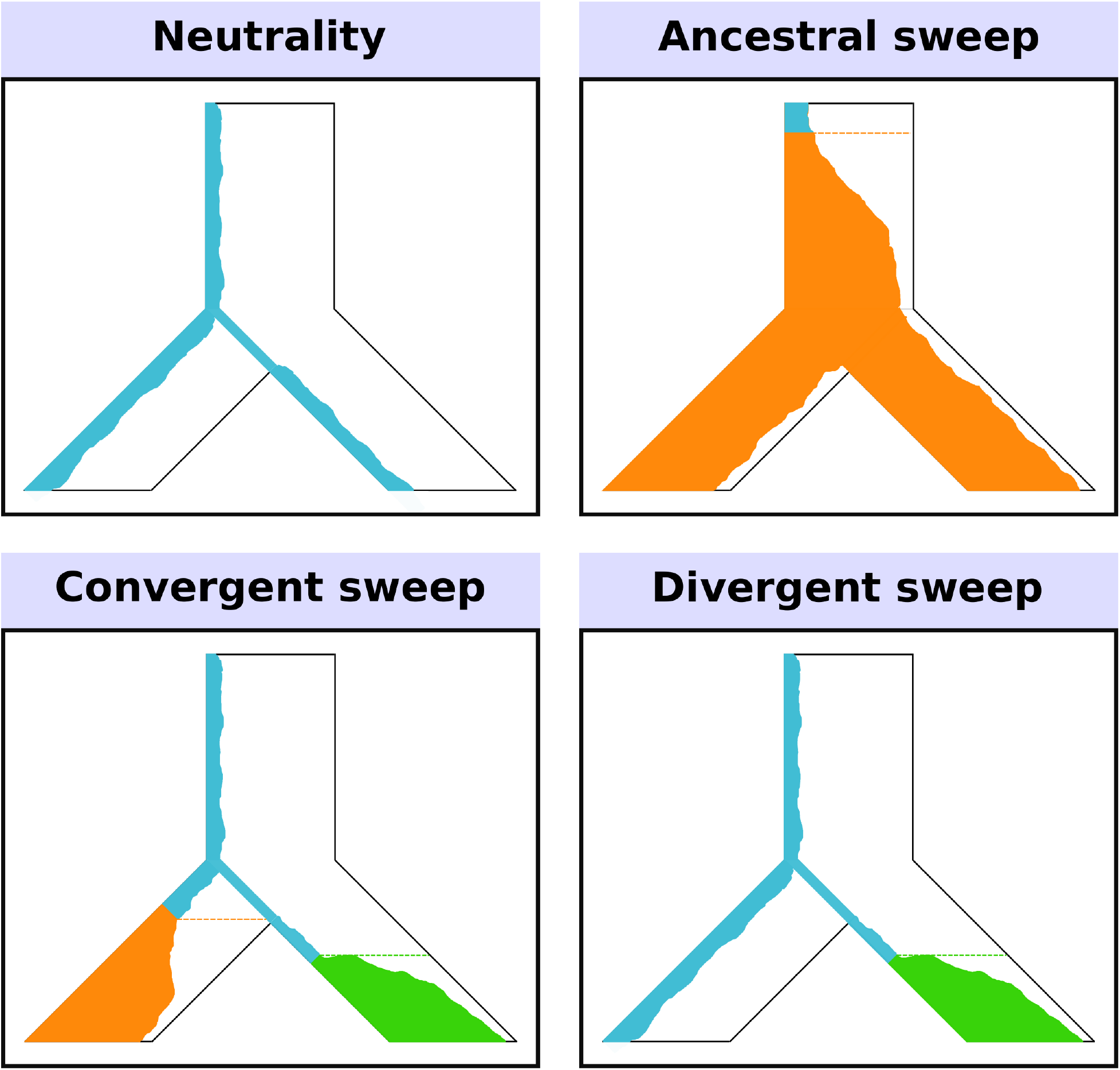
Model of a two-population phylogeny for which SS-H12 detects recent shared sweeps. Here, an ancestral population splits in the past into two modern lineages, which are sampled. Each panel displays the frequency trajectory of a haplotype in the population. Under neutrality, there is high haplotypic diversity such that many haplotypes, including the reference haplotype (blue), exist at low frequency. In the ancestral sweep, the reference haplotype becomes selectively advantageous (turning orange) and rises to high frequency prior to the split, such that both modern lineages carry the same selected haplotype at high frequency. The convergent sweep scenario involves different selected haplotypes independently rising to high frequency in each lineage after their split. Under a divergent sweep, only one sampled lineage experiences selection.

We now define SS-H12, which provides information about the location of a shared sweep on the phylogenetic tree relating the sampled populations, and describe its application to a sample consisting of individuals from two populations. SS-H12 is computed from multiple statistics that quantify the diversity of haplotypes within each population, as well as within the pool of the populations, and is a modification of the single-population expected homozygosity-based approach. Consider a pooled sample consisting of haplotypes from two populations, in which a fraction *γ* of the haplotypes derives from population 1 and a fraction 1 − *γ* derives from population 2. For a pooled sample consisting of individuals from two populations, we define the total-sample expected haplotype homozygosity statistic H12_Tot_ within a genomic window containing *I* distinct haplotypes as

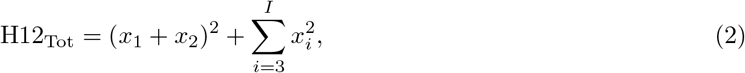

where *x_i_* = *γp*_1*i*_ + (1 − *γ*)*p*_2*i*_, *x*_1_ ≥ *x*_2_ ≥ ⋯ ≥ *x_I_*, is the frequency of the *ith* most frequent haplotype in the pooled population, and where *p*_1*i*_ and *p*_2*i*_ are the frequencies of this haplotype in populations 1 and 2, respectively. H12_Tot_ therefore has elevated values at the sites of shared sweeps because the pooled genetic diversity at a site under selection in each sampled population remains small.

Next, we seek to define a statistic that classifies the putative shared sweep as ancestral or convergent between the pair of populations. To do this, we define a statistic 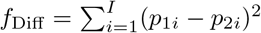, which measures the sum of the squared difference in the frequency of each haplotype between both populations. *f*_Diff_ takes on values between 0, for population pairs with identical haplotype frequencies, and 2, for populations that are each fixed for a different haplotype. The former case is consistent with an ancestral sweep scenario, whereas the latter is consistent with a convergent sweep—though we caution that genetic drift can also produce extreme values of *f*_Diff_, which is unlikely to be problematic provided test populations are closely enough related.

From the summary statistics H12_Tot_ and *f*_Diff_, we now formulate SS-H12, which measures the extent to which an elevated H12_Tot_ is due to shared ancestry. First, we specify a statistic that quantifies the shared sweep, H12_Anc_ = H12_Tot_ − *f*_Diff_. The value of H12_Anc_ lies between −1 for convergent sweeps, and 1 for ancestral sweeps, with a negative value near 0 in the absence of a sweep. H12_Anc_ is therefore easy to interpret because convergent sweeps on non-identical haplotypes cannot generate positive values, and ancestral sweep signals that have not eroded due to the effects of recombination and mutation cannot generate negative values. Because a sufficiently strong and complete sweep in one population (divergent sweep; Figure 1) may also yield negative values of H12_Anc_ with elevated magnitudes distinct from neutrality, we introduce a correction factor that yields SS-H12 by dividing the minimum value of H12 between a pair of populations by the maximum value. This modification allows SS-H12 to overlook spurious signals driven by strong selection in a single population by reducing their prominence relative to true shared sweep signals. Applying this correction factor yields SS-H12, which is computed as

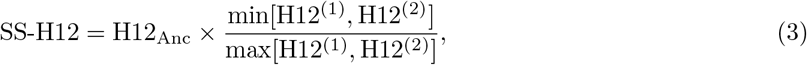

where H12^(1)^ and H12^(2)^ are the H12 values for populations 1 and 2, respectively. The correction factor has a value close to 1 for shared sweeps of either type, but a small value for divergent sweeps. Thus, the corrected SS-H12 is sensitive only to shared sweeps.

Using simulated genetic data, we show that SS-H12 has high power to detect recent shared selective sweeps in pairs of populations, displaying a similar range of detection to the single-population H12 statistic on which it is based. Additionally, we demonstrate that, in accordance with our expectations, SS-H12 correctly identifies recent ancestral sweeps from elevated positive values, and convergent sweeps from negative values of large magnitude, generally without confusing the two scenarios. Furthermore, we extended the application of SS-H12 to an arbitrary number of populations *K* (see *Materials and Methods*), finding once again that our approach classifies sweeps correctly and with high power. Moreover, the SS-H12 approach retains the ability to distinguish between ancestral and convergent hard and soft sweeps from the inferred number of distinct sweeping haplotypes, with each sweep type occupying a distinct subset of paired (|SS-H12|, H2_Tot_/H1_Tot_) values, where 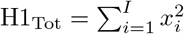 and 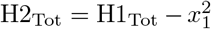, with *x_i_* defined as in Equation 2. As with H2/H1 for the single-population approach, H2_Tot_/H1_Tot_ is larger for soft sweeps and smaller for hard sweeps. Finally, our analysis of whole-genome sequences from global human populations recovered previously-identified sweep candidates at the *LCT* and *SLC24A5* genes in Indo-European populations, corroborated recently-characterized sweeps that emerged from genomic scans with the single-population approach [Harris et al., 2018], such as *RGS18* in African and *P4HA1* in Indo-European populations, and uncovered novel shared sweep candidates, such as the convergent sweeps *C2CD5* between Eurasian populations and *PAWR* between European and sub-Saharan African populations.

## Results

We evaluated the ability of SS-H12 to differentiate among the simulated scenarios of shared selective sweeps, sweeps unique to only one sampled population, and neutrality, using the signature of expected haplotype homozygosity in samples consisting of individuals from two or more populations. Although our formulation of SS-H12 does not explicitly constrain the definition of a population, we define a population as a discrete group of individuals that mate more often with each other than they do with individuals from other discrete groups, and the models we considered here represent extreme examples in which there is no gene flow between populations. We performed simulations using SLiM 2 [Haller and Messer, 2017] under human-inspired parameters [Takahata et al., 1995, Nachman and Crowell, 2000, Payseur and Nachman, 2000, Terhorst et al., 2017, Narasimhan et al., 2017] for diploid populations of fluctuating size (*N*) under non-equilibrium models, as well as constant-size models, subject to changing selection start times (*t*) and strengths (*s*), across differing split times (*τ*) between sampled populations. Additionally, we evaluated the robustness of SS-H12 to confounding scenarios of population admixture and background selection. We then used an approximate Bayesian computation (ABC) approach in the same manner as Harris et al. [2018] to demonstrate our ability to distinguish between shared hard and soft sweeps in samples from multiple populations. Finally, we show that SS-H12 recovers previously-hypothesized signatures of shared sweeps in human whole-genome sequences [Auton et al., 2015], while also uncovering novel candidates. See *Materials and Methods* for further explanation of experiments.

### Detection of ancestral and convergent sweeps with SS-H12

We conducted experiments to examine the ability of SS-H12 to not only identify shared sweep events among two or more sampled populations (*K*), but categorize them as shared due to common ancestry, or due to convergent evolution. Across all scenarios, we scanned 100 kb simulated chromosomes using a 20 or 40 kb sliding window with a step size of one kb. We selected windows of this size because it is over this interval that neutral pairwise LD, measured with *r*^2^, decays to 1/3 of the value for loci one kb apart (Figure S1). This means that we do not expect elevated values of SS-H12 to arise from patterns of background LD. For each sweep scenario, we studied the power at 1% and 5% false positive rates (FPRs) for detecting shared selective sweeps (Figures 2, 3, and S2-S9) as a function of time at which beneficial alleles arose, under scenarios of ancestral, convergent, and divergent sweeps.

**Figure 2:**
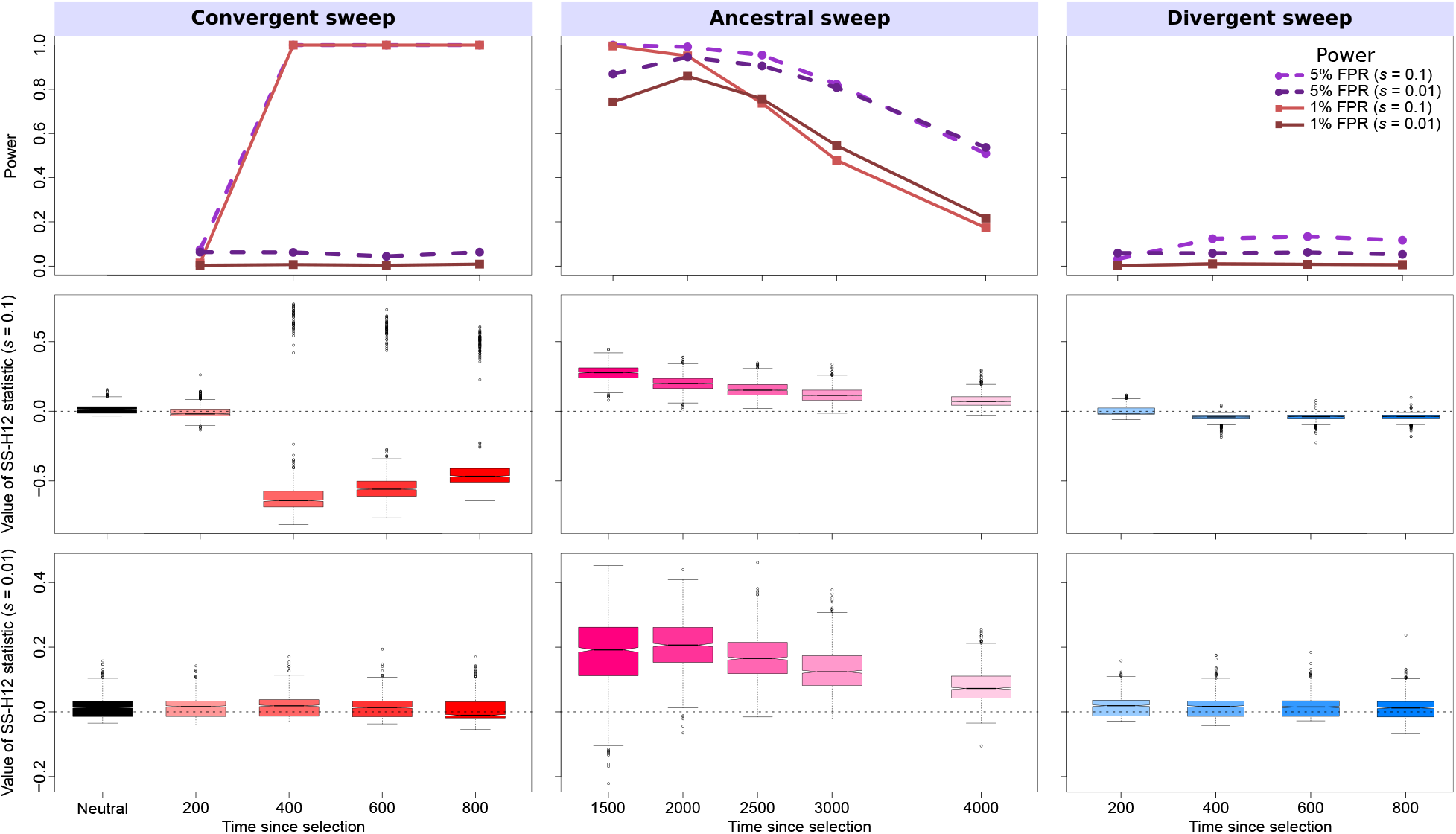
Properties of SS-H12 for simulated strong (*s* = 0.1) and moderate (*s* = 0.01) hard sweep scenarios under the CEU-GIH model (*τ* = 1100 generations before sampling). (Top row) Power at 1% (red lines) and 5% (purple lines) false positive rates (FPRs) to detect recent ancestral, convergent, and divergent hard sweeps (see Figure 1) as a function of time at which positive selection of the favored allele initiated (*t*), with FPR based on the distribution of maximum |SS-H12| across simulated neutral replicates. (Middle row) Box plots summarizing the distribution of SS-H12 values from windows of maximum |SS-H12| across strong sweep replicates, corresponding to each time point in the power curves, with dashed lines in each panel representing SS-H12 = 0. (Bottom row) Box plots summarizing the distribution of SS-H12 values across moderate sweep replicates. For convergent and divergent sweeps, *t* < *τ*, while for ancestral sweeps, *t* > *τ*. All replicate samples for the CEU-GIH model contain 99 simulated CEU individuals and 103 simulated GIH individuals, as in the 1000 Genomes Project dataset [Auton et al., 2015], and we performed 1000 replicates for each scenario.

**Figure 3:**
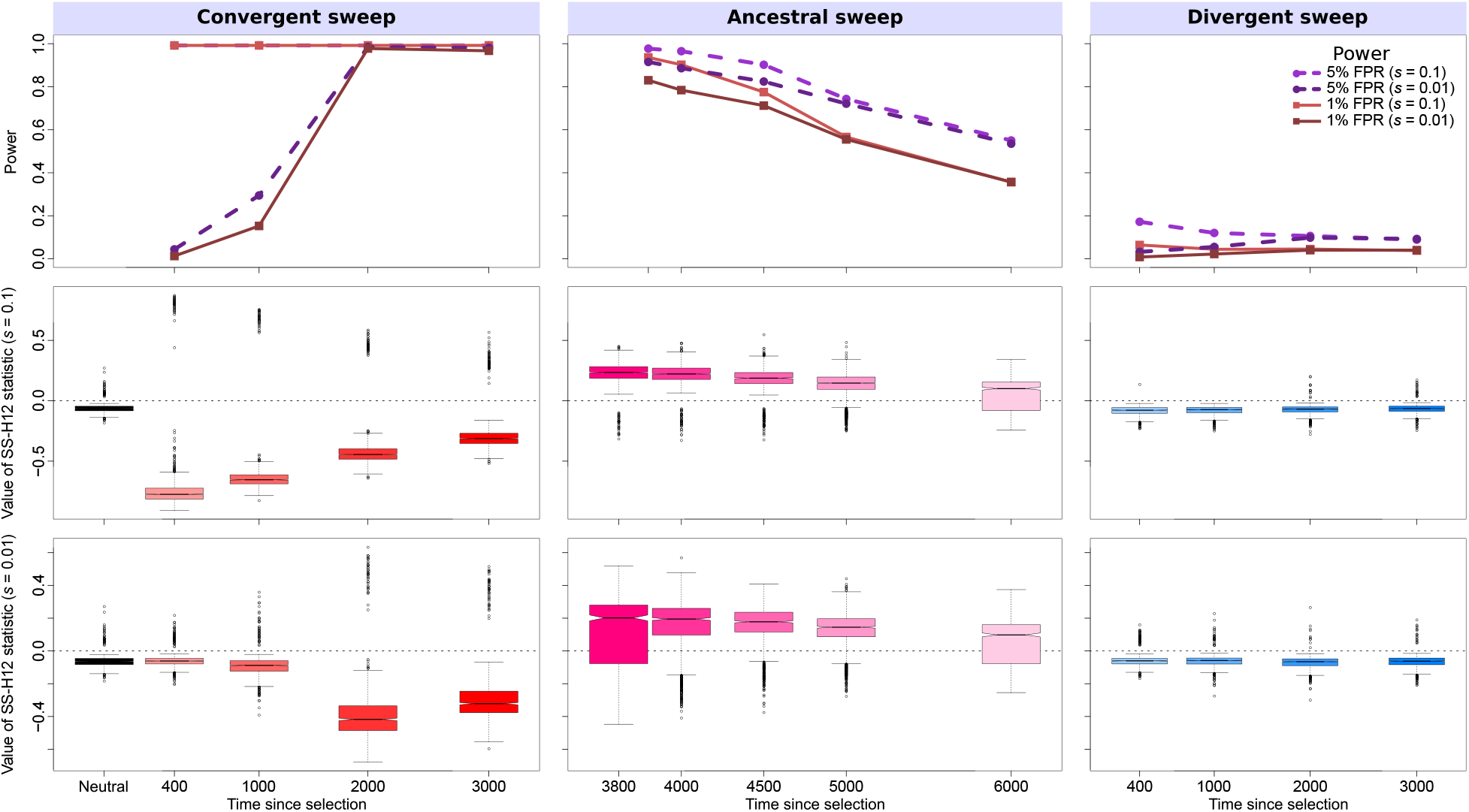
Properties of SS-H12 for simulated strong (*s* = 0.1) and moderate (*s* = 0.01) hard sweep scenarios under the CEU-YRI model (*τ* = 3740 generations before sampling). (Top row) Power at 1% (red lines) and 5% (purple lines) false positive rates (FPRs) to detect recent ancestral, convergent, and divergent hard sweeps (see Figure 1) as a function of time at which positive selection of the favored allele initiated (*t*), with FPR based on the distribution of maximum |SS-H12| across simulated neutral replicates. (Middle row) Box plots summarizing the distribution of SS-H12 values from windows of maximum |SS-H12| across strong sweep replicates, corresponding to each time point in the power curves, with dashed lines in each panel representing SS-H12 = 0. (Bottom row) Box plots summarizing the distribution of SS-H12 values across moderate sweep replicates. For convergent and divergent sweeps, *t* < *τ*, while for ancestral sweeps, *t* > *τ*. All replicate samples for the CEU-YRI model contain 99 simulated CEU individuals and 108 simulated YRI individuals, as in the 1000 Genomes Project dataset [Auton et al., 2015], and we performed 1000 replicates for each scenario.

First, we simulated scenarios in which an ancestral population split into *K* = 2 descendant populations using a realistic non-equilibrium model based on the history of the human CEU (European descent) and GIH (South Asian descent) populations, which we inferred from variant calls [Auton et al., 2015] with `smc++` [Terhorst et al., 2017]. We first examined scenarios of strong (*s* = 0.1), hard (*ν* = 1 sweeping haplotype) sweeps starting between 200 and 4000 generations prior to sampling and applied an analysis window of size 40 kb (Figure 2). Our CEU-GIH model features a split time of *τ* = 1100 generations prior to sampling, which matches prior estimates of the split time between Eurasian human populations [Gravel et al., 2011, Gronau et al., 2011, Schiffels and Durbin, 2014]. This series of experiments illustrates the range of sweep start times over which SS-H12 can detect prominent selective sweeps. SS-H12 has high power for recent strong shared sweeps starting between 400 and 2500 generations prior to sampling, with power dropping rapidly for shared sweeps older than 2500 generations (Figure 2). As expected, the distributions of SS-H12 values for detectable convergent sweeps center on negative values (Figure 2), whereas the SS-H12 distributions of ancestral sweeps center on positive values (Figure 2). The power of SS-H12 to detect convergent sweeps for *t* ∈ [400, 1000] is greater because these sweeps are more recent, with the selected haplotype having established in the population, and not yet eroding due to the effect of mutation and recombination, as with older ancestral sweeps. Additionally, because we compute power from the distribution of maximum |SS-H12| values for each sweep scenario, this means that the magnitude of SS-H12 for replicates of shared sweeps must exceed the magnitude under neutrality for the sweep to be detected, which for any combination of *t* and *s* is more likely for convergent than ancestral sweeps.

To further characterize the performance of SS-H12 for hard sweeps, we repeated experiments on simulated samples from *K* = 2 populations for more anciently-diverged populations (larger *τ*), and weaker sweeps (smaller *s*). SS-H12 maintains excellent power to distinguish strong shared sweeps from neutrality for a model based on the split between CEU and the sub-Saharan African YRI population (Figure 3; 20 kb window). We inferred *τ* = 3740 for this model using `smc++` [Terhorst et al., 2017] once again, and this estimate fits existing estimates of split times between African and non-African human populations [Gravel et al., 2011, Gronau et al., 2011, Schiffels and Durbin, 2014]. Notably, the signal of ancestral sweeps remains elevated across many of the tested sweep scenarios, staying above 0.6 for sweeps starting *t* = 4500 generations before sampling, extending our range of sweep sensitivity at this threshold by approximately 1500 generations relative to the CEU-GIH model. This is because it is easier to detect selective sweeps in more diverse genomic backgrounds [Harris et al., 2018], such as that of the YRI population, despite the advanced age of simulated ancestral sweeps. Despite this, we observed a greater proportion of ancestral sweeps with spuriously negative values of SS-H12 in the CEU-YRI model than in the CEU-GIH model because over 3740 generations, the two simulated populations had sufficient time to accumulate unique mutations and recombination events that differentiated their common high-frequency haplotypes. Reducing the selection coefficient to *s* = 0.01 for both CEU-GIH and CEU-YRI scenarios had the effect of shifting the range of *t* over which SS-H12 had high power to detect shared sweeps. Because weakly-selected haplotypes rise to high frequency more slowly than strongly-selected haplotypes, there is a greater delay between the selection start time and the time at which a shared sweep can be detected for smaller values of *s*. Thus, SS-H12 reaches a maximum power to detect moderate shared sweeps (*s* = 0.01) for older values of t, additionally maintaining this power for less time than for strong sweeps under both models (Figures 2 and 3).

Because the single-population statistic H12 has power to detect both hard and soft sweeps, we next performed analogous experiments for simulated soft sweep scenarios. Maintaining values of *t, τ*, and *s* identical to those for hard sweep experiments, we simulated soft sweeps as selection on standing genetic variation for *ν* = 4 and *ν* = 8 distinct sweeping haplotypes (Figures S2-S5). We found that trends in the power of SS-H12 to detect shared soft sweeps remained consistent with those for hard sweeps. However, the power of SS-H12 for detecting soft sweeps was attenuated overall relative to hard sweeps, proportionally to the number of sweeping haplotypes, with a larger drop in power for older sweeps and little to no effect on power for more recent sweeps. Our observations therefore align with results for the single-population H12 statistic [Garud et al., 2015, Harris et al., 2018]. Thus, the ability to detect a sweep derives from the combination of *s, t*, and *ν*, with stronger recent sweeps on fewer haplotypes being easiest to detect, and detectable over larger timespans.

We contrast our results for shared sweeps across population pairs with those for divergent sweeps, which we present in parallel (Figures 2, 3, and S2-S5). Across identical values of *t* as for each convergent sweep experiment, we found that divergent sweeps, in which only one of the two simulated sampled populations experiences a sweep (*t* < *τ*), are not visible to SS-H12 for any combination of simulation parameters. To understand the properties of divergent sweeps relative to shared sweeps, we compared the distributions of their SS-H12 values at peaks identified from the maximum values of |SS-H12| for each replicate. We observed that the distributions of the divergent sweeps remain broadly unchanged from one another under all parameter combinations, and closely resemble the distribution generated under neutrality, as all are centered on negative values with small magnitude, and have small variance. Thus, the use of a correction factor that incorporates the values of H12 from each component population in the sample (see Equation 3) provides an appropriate approach for preventing sweeps that are not shared from appearing as outlying signals.

Next, we evaluated the power of SS-H12 across samples simulated under a simplified demographic history with a constant diploid population size of *N* = 10^4^ and higher mutation and recombination rates (intended as a more general eukaryotic model rather than specifically human; see *Materials and Methods*) in order to evaluate the effect of changing the number of sampled populations *K*. Our samples consisted of *K* = 2 (*τ* = 1000), 3 (*τ*_1_ = 1000 and *τ*_2_ = 750), 4 (*τ*_1_ = 1000, *τ*_2_ = 750, and *τ*_3_ = 500), and 5 (*τ*_1_ = 1000, *τ*_2_ = 750, *τ*_3_ = 500, and *τ*_4_ = 250) sampled populations experiencing strong sweeps (*s* = 0.1). We found that SS-H12 maintains power to identify shared sweeps, but that the properties of the method change somewhat as more populations are sampled (Figures S6-S9). To assign values of SS-H12 to samples of *K* ≥ 3 populations, we employed two types of approaches, and found that these generally yielded comparable power to detect simulated shared sweeps. First, we measured SS-H12 for each possible population pair within the sample, and conservatively retained the value of smallest magnitude as the overall SS-H12. Second, we computed SS-H12 across the two branches directly subtending the root of the *K*-population phylogeny underlying the sample, grouping together all populations descending from the same internal branch (see *Materials and Methods*).

For both the conservative and grouped approaches, the power of SS-H12 to detect strong shared sweeps is high for sweeps more recent than 1500 generations ago, and rapidly attenuates for more ancient sweeps, and power is once again greatest for convergent sweeps. Furthermore, trends in power for detecting shared sweeps remained consistent between *K* = 2 and *K* > 2 scenarios, regardless of the choice of approach. However, we found that despite maintaining perfect or near-perfect power for convergent sweeps on samples from *K* ≥ 3 populations, the distribution of SS-H12 includes many replicates with positive values, which are normally associated with ancestral sweeps. The shift toward positive values increases as the convergent sweep becomes more ancient, reflecting a greater fraction of ancestral sweeps between pairs of sampled populations within the overall convergent sweep. Although the conservative approach remains generally more robust to misclassifying shared sweeps within samples from *K* ≥ 3 populations than does the grouped approach, both strategies may fail to identify a convergent sweep as convergent if the sweep time *t* is close enough to *τ*_1_. Additionally, divergent sweeps yield a distribution of SS-H12 values for samples from *K* ≥ 3 populations that may differ from neutrality as *t* approaches *τ*_1_. Despite this observation, we emphasize that divergent sweeps once again do not produce values of SS-H12 that deviate appreciably from values generated under neutrality, leaving shared sweeps as the sole source of prominently outlying sweep signals in practice.

In addition to detecting shared sweeps under a variety of scenarios with high power, we also found that detecting sweeps with SS-H12 provides more power than performing multiple independent analyses across populations with the single-population statistic H12 [Garud et al., 2015]. To demonstrate this, we reanalyzed our simulated CEU-GIH and CEU-YRI replicates (Figures 2 and 3), assessing the ability of H12 to simultaneously detect an outlying sweep signal in both populations. That is, we measured the power of H12 at the 0.5% FPR (Bonferroni-corrected for multiple testing [Neyman and Pearson, 1928], providing the entire experiment with a 1% FPR cutoff) to detect an outlying sweep in the CEU sample *and* in either the GIH (Figure S10) or YRI (Figure S11) samples. For the most recent convergent hard sweeps, joint analysis with H12 has equivalent power to SS-H12 analysis, but the power of H12 never matches that of SS-H12 for ancestral hard sweeps, and for the majority of tested soft sweeps (*v* = 4 and *ν* = 8), regardless of timing. These trends persisted even for SS-H12 computed from half-sized samples (thus, matching the sample sizes of individual H12 analyses), indicating that avoiding multiple testing with SS-H12 analysis is likely to yield a greater return on sampling effort, especially as the number of sampled populations *K* increases.

### Addressing confounding scenarios

Because SS-H12 relies on a signal of elevated expected haplotype homozygosity, it may be confounded by non-adaptive processes that alter levels of population-genetic diversity. For this reason, we first examined the effect of admixture on the power of SS-H12 in the context of ancestral, convergent, and divergent strong (*s* = 0.1) sweeps between population pairs simulated under the aforementioned simplified model, separated by *τ* = 1000 generations, wherein one population (the target) receives gene flow from a diverged, unsampled donor outgroup population (Figures 4 and S12). Admixture occurred as a single pulse 200 generations before sampling, and in the case of the divergent sweep, occurred specifically in the population experiencing the sweep. The donor split from the common ancestor of the two sampled populations (the target and its unadmixed sister) 2 × 10^4^ generations before sampling—within a coalescent unit of the sampled populations, similar to the relationship between Neanderthals and modern humans [Juric et al., 2016, Harris and Nielsen, 2016])—and had an effective size either one-tenth, identical to, or tenfold the size of the target. Although the donor does not experience selection, extensive gene flow from a donor with low genetic diversity may resemble a sweep. Correspondingly, gene flow from a highly diverse donor may obscure sweeps.

**Figure 4:**
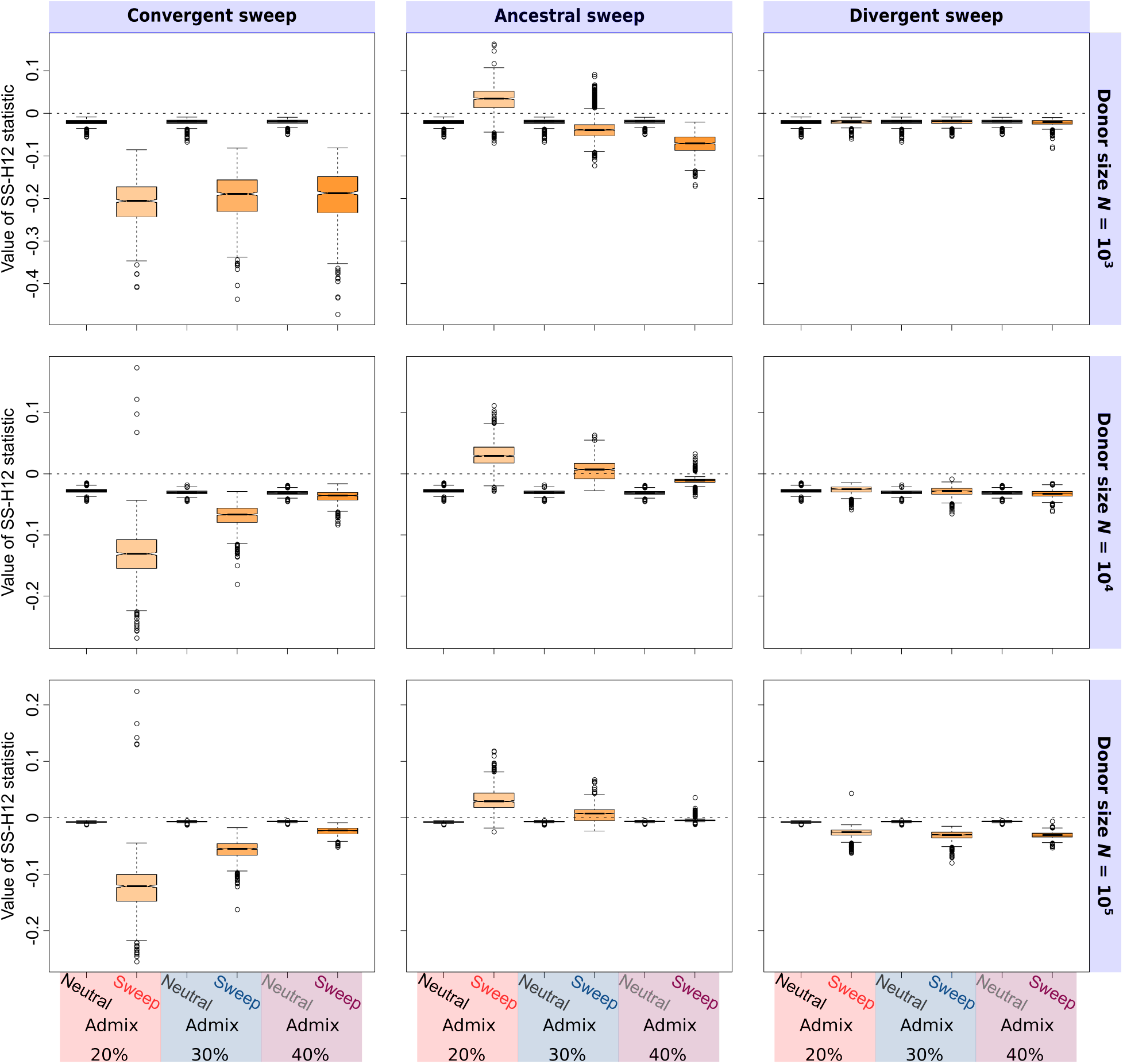
Effect of admixture from a diverged, unsampled donor lineage on distributions of SS-H12 values at peaks of maximum |SS-H12|, in samples consisting of individuals from *K* = 2 populations with *τ* = 1000, under simulated recent ancestral, convergent, and divergent histories (see Figure 1), maintaining a constant effective size in the sampled populations of *N* = 10^4^ diploids. For ancestral sweeps, selection occurred 1400 generations before sampling. For convergent and divergent sweeps, selection occurred 600 generations before sampling. The effective size of the donor population varies from *N* = 10^3^ (an order of magnitude less than the sampled populations), to *N* = 10^5^ (an order of magnitude more), with admixture occurring 200 generations before sampling at rates 0.2 to 0.4, modeled as a single pulse. The donor diverged from the sampled populations 2 × 10^4^ = 2*N* generations before sampling and in the case of divergent sweep scenarios, admixed specifically into the population experiencing a sweep. All sample sizes are of *n* = 100 diploid individuals, with 1000 replicates performed for each scenario.

As expected, gene flow into the target population distorted the SS-H12 distribution of the two-population sample relative to no admixture (compare Figure 4 to S6), and this distortion was proportional to the level of admixture from the donor, as well as the donor’s effective size (Figure 4). Ancestral sweeps were the most likely to be misclassified following admixture from an unsampled donor of small effective size (*N* = 10^3^; Figure 4, first row), increasingly resembling convergent sweeps as the rate of gene flow increased (though ultimately with little change in power to detect the shared sweep; Figure S12). The confounding effect of admixture on ancestral sweep inference emerges because low-diversity gene flow into one population yields a differing signal of elevated expected haplotype homozygosity in each population, spuriously resembling a convergent sweep. In contrast, the distributions of SS-H12 values and the power of SS-H12 for convergent and divergent sweeps remained broadly unchanged relative to no admixture (Figure S6) under low-diversity admixture scenarios (compare panels within the first rows of Figures 4 and S12 to Figure S6). Because two populations subject to convergent or divergent sweeps are already extensively differentiated, further differentiation due to admixture does not impact the accuracy of sweep timing classification using SS-H12.

For intermediate donor effective size (*N* = 10^4^; Figures 4 and S12, second row), the magnitudes of both the ancestral and convergent sweep signals attenuate toward neutral levels, and the power of SS-H12 wanes as the admixture proportion increases. This is because the genetic diversity in the target population increases to levels resembling neutrality, overall yielding a pattern spuriously resembling a divergent sweep that SS-H12 cannot distinguish from neutrality. Accordingly, the magnitude and power of SS-H12 under a divergent sweep scenario following admixture scarcely change under the *N* = 10^4^ scenario. As the effective size of the donor population grows large (*N* = 10^5^; Figures 4 and S12, third row), SS-H12 becomes more robust to the effect of admixture for shared sweeps, accurately identifying ancestral and convergent sweeps with high power at greater admixture proportions relative to the *N* = 10^4^ scenario. However, the power of SS-H12 spuriously rises to 1.0 for divergent sweeps under the *N* = 10^5^ admixture scenario. Both the increased robustness to admixture for the ancestral and convergent sweeps, as well as the elevated power for divergent sweeps, result from a reduction in the magnitude of SS-H12 under neutrality for the *N* = 10^5^ admixture scenario relative to *N* = 10^4^, which does not occur for the sweep scenarios. That is, the magnitude of a sweep signature remains similar across the *N* = 10^5^ and *N* = 10^4^ admixture scenarios, while the magnitude of the neutral background is smaller, meaning that any sweep, even a divergent sweep, is more prominent for larger donor population sizes. We further address this observation in the *Discussion*.

We also observed the effect of long-term background selection on the neutral distribution of SS-H12 values (Figure S13). Background selection may yield signatures of genetic diversity resembling selective sweeps [Charlesworth et al., 1993, 1995, Seger et al., 2010, Nicolaisen and Desai, 2013, Cutter and Payseur, 2013, Huber et al., 2016], though previous work suggests that background selection does not drive particular haplotypes to high frequency [Enard et al., 2014, Harris et al., 2018]. Our two background selection scenarios for samples from *K* = 2 populations, with *τ* = 1100 (CEU-GIH model) and 3740 (CEU-YRI model) generations, were performed as described in the *Materials and Methods*, following the protocol of Cheng et al. [2017]. Briefly, we simulated a 100-kb sequence featuring a centrally-located 11-kb gene consisting of exons, introns, and untranslated regions, across which deleterious variants arose randomly throughout the entire simulation period. In agreement with our expectations, we found that background selection is unlikely to confound inferences from SS-H12, yielding only marginally larger values of |SS-H12| than does neutrality (Figure S13). Accordingly, SS-H12 does not classify background selection appreciably differently from neutrality.

### Classifying shared sweeps as hard or soft from the number of sweeping haplotypes

Because the primary innovation of the single-population approach is its ability to classify sweeps as hard or soft from paired (H12, H2/H1) values, we evaluated the corresponding properties of our current approach for samples consisting of *K* = 2 populations (Figure 5). Here, we color a space of paired (|SS-H12|, H2_Tot_/H1_Tot_) values, each bounded by [0.005, 0.995], according to the inferred most probable number of sweeping haplotypes *ν* for each point in the space. Similarly to the approach of Harris et al. [2018], we inferred the most probable *ν* using an approximate Bayesian computation (ABC) approach in which we determined the posterior distribution of *ν* from 5 × 10^6^ replicates of sweep scenarios with *ν* ∈ {0, 1,…, 16} and *s* ∈ [0.005, 0.5], both drawn uniformly at random for each replicate (the latter drawn from a log-scale), and where *ν* = 0 simulations are neutral replicates. A test point in (|SS-H12|, H2_Tot_/H1_Tot_) space was assigned a value of *ν* based on the most frequently occurring *ν* among simulations whose (|SS-H12|, H2_Tot_/H1_Tot_) coordinates were within a Euclidean distance of 0.1 from that test point (see *Materials and Methods*). We were able to classify recent shared sweeps as hard or soft, but found our current approach to have somewhat different properties to the single-population approach (Figure 5).

**Figure 5:**
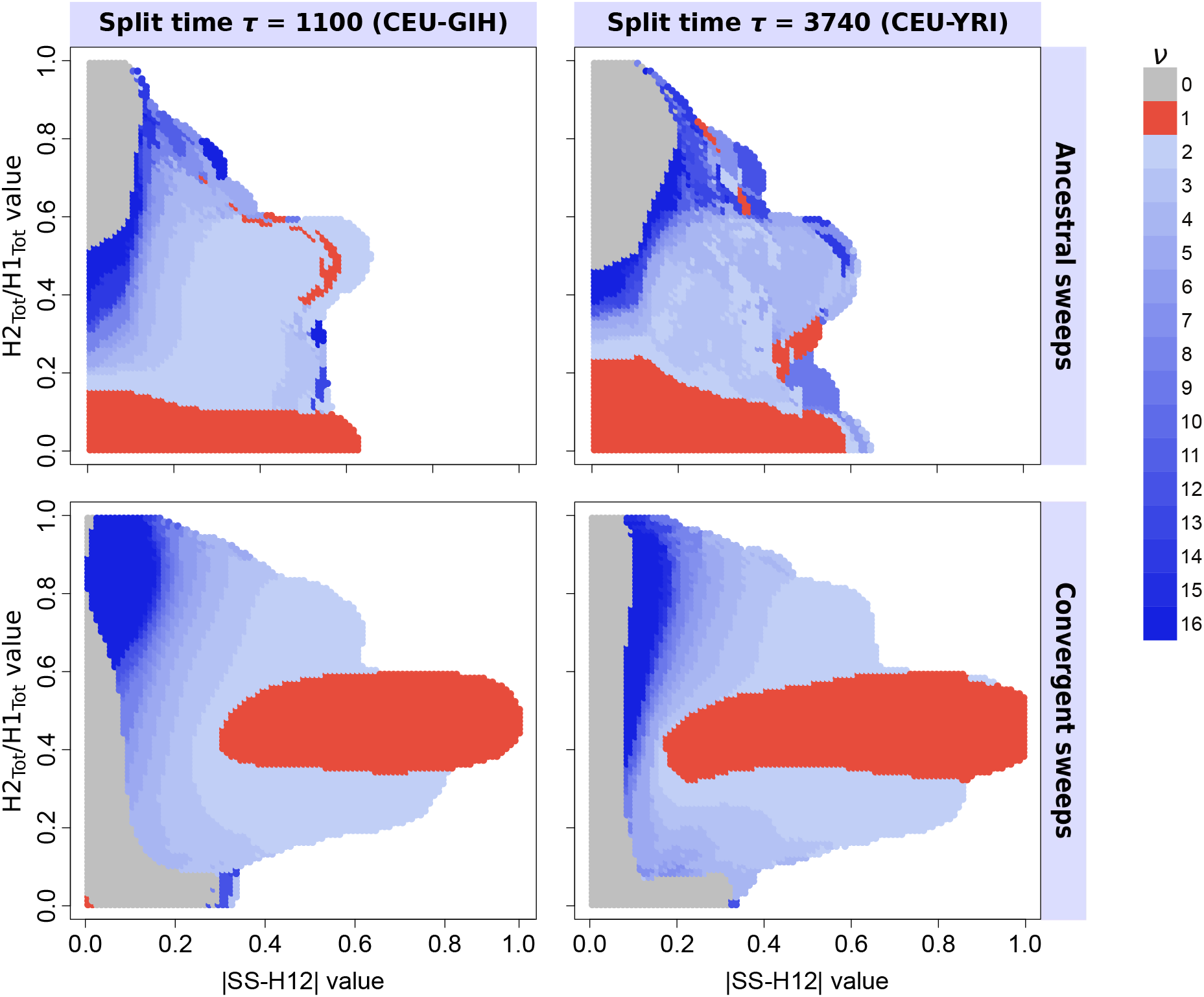
Ability of paired (|SS-H12|, H2_Tot_/H1_Tot_) values to infer the most probable number of sweeping haplotypes *ν* in a shared sweep. Most probable *ν* for each test point was assigned from the posterior distribution of 5 × 10^6^ sweep replicates with *ν* ∈ {0,1,…, 16}, drawn uniformly at random. (Top row) Ancestral sweeps for the CEU-GIH model (*τ* = 1100, left) and the CEU-YRI model (*τ* = 3740, right), with *t* ∈ [1140, 3000] (left) and *t* ∈ [3780, 5000] (right). (Bottom row) Convergent sweeps for the CEU-GIH model (left) and the CEU-YRI model (right), with *t* ∈ [200, 1060] (left) and *t* ∈ [200, 3700] (right). Colored in red are points whose paired (|SS-H12|, H2_Tot_/H1_Tot_) values are more likely to result from hard sweeps, those colored in shades of blue are points more likely to be generated from soft sweeps, and gray indicates a greater probability of neutrality. Regions in white are those for which no observations of sweep replicates within a Euclidean distance of 0.1 exist.

For ancestral sweep scenarios and *τ* = 1100 generations (*t* ∈ [1140, 3000], CEU-GIH model), the pattern of paired (|SS-H12|, H2_Tot_/H1_Tot_) values generally followed that of single-population analyses [Harris et al., 2018], though with irregularities among larger values of |SS-H12| paired with intermediate values of H2_Tot_/H1_Tot_ (Figure 5, top left). For a given value of |SS-H12|, smaller values of H2_Tot_/H1_Tot_ were generally more probable for ancestral sweeps from smaller *v*, and inferred *ν* increased with H2_Tot_/H1_Tot_. This fit our expectations because, as the number of ancestrally sweeping haplotypes in the pooled population increases, the value of H2_Tot_ increases relative to H1_Tot_. Additionally, ancestral sweeps from larger *ν* (softer sweeps) are unlikely to generate large values of |SS-H12| or small values of H2_Tot_/H1_Tot_, and the most elevated values of |SS-H12| were rarely associated with more than four sweeping haplotypes. We note, however, the presence of paired values inferred to derive from *ν* = 1 for some intermediate values of H2_Tot_/H1_Tot_, as well as the presence of points with inferred *ν* ≥ 4 at smaller H2_Tot_/H1_Tot_. This may indicate that among ancestral sweep replicates for the CEU-GIH model, weaker hard sweep signals may occasionally be difficult to resolve from stronger soft sweep signals, as both should yield intermediate levels of haplotypic diversity. Under simulated scenarios of *τ* = 3740 generations (*t* ∈ [3780, 5000], CEU-YRI model; Figure 5, top right), we observed a broadly similar pattern of inferred *ν*, but the increased age of ancestral sweeps under this model relative to the CEU-GIH model resulted in more erratic inferences across intermediate |SS-H12| paired with intermediate H2_Tot_/H1_Tot_. Our approach still maintains a clear tendency to infer sweeps with smaller H2_Tot_/H1_Tot_ as hard, preserving the basic classification ability.

The convergent sweep experiments yielded a distinctly different occupancy of possible paired (|SS-H12|, H2_Tot_/H1_Tot_) values relative to ancestral sweeps, and provided a greater resolution among the tested values of *v*, showing little irregularity in the assignment of *ν* (Figure 5, bottom row). In addition, trends in the occupancy of hard and soft sweeps were generally concordant between replicates for both the CEU-GIH (*τ* = 1100, *t* ∈ [200, 1060]) and CEU-YRI (*τ* = 3740, *t* ∈ [200, 3700]) models. For these experiments, we simulated simultaneous independent sweeps, either both soft or both hard, allowing each population to follow a unique but comparable trajectory. Thus, there were always at least two sweeping haplotypes in the pooled population. Accordingly, convergent hard sweeps, unlike ancestral hard sweeps, are primarily associated with large values of |SS-H12| and intermediate values of H2_Tot_/H1_Tot_. Furthermore, strong convergent sweeps of any sort could not generate small H2_Tot_/H1_Tot_ values unless |SS-H12| was also small. Even so, convergent sweeps from larger *ν* occupy a distinct set of paired (|SS-H12|, H2_Tot_/H1_Tot_) values that is shifted either toward smaller |SS-H12|, larger H2_Tot_/H1_Tot_, or both, demonstrating that the accurate and consistent inference of *ν* is possible for convergent sweeps. Unlike for ancestral sweeps or single-population analyses, we observed that the smallest values of H2_Tot_/H1_Tot_ paired with the smallest values of |SS-H12| were associated with neutrality, representing scenarios in which similar highly diverse haplotype frequency spectra arose in both populations by the time of sampling.

### Application of SS-H12 to human genetic data

We applied SS-H12 to whole-genome sequencing data from global human populations in phase 3 of the 1000 Genomes Project [Auton et al., 2015], which is ideal as input because it contains large sample sizes and no missing genotypes at polymorphic sites. We searched for shared sweep signals within the RNA- and protein-coding genes of geographically proximate and distant human population pairs, performing various comparisons of unadmixed European, South Asian, East Asian, and Sub-Saharan African populations (Tables S2-S10). For the top 40 outlying candidate shared sweeps among population pairs, we assigned *p*-values from a neutral distribution of 10^6^ replicates following human demographic models inferred from `smc++` (see *Materials and Methods*). Our Bonferroni-corrected significance threshold [Neyman and Pearson, 1928] was 2.10659 × 10^−6^ (see Table S1 for critical values associated with each population pair). We additionally inferred the maximum posterior estimates on *ν* ∈ {1, 2,…, 16} for each top candidate from a distribution of 5 × 10^6^ simulated convergent or ancestral sweep replicates, depending on our classification of the candidate from the sign of SS-H12, following the same `smc++`-derived models. We categorized sweeps from *ν* = 1 as hard, and sweeps from *ν* ≥ 2 as soft.

Across all comparisons, we found that ancestral hard sweeps comprised the majority of prominent candidates at RNA- and protein-coding genes, regardless of population pair. Many of these candidate ancestral sweeps were detected with H12 in single populations [Harris et al., 2018], including novel sweeps at *RGS18* in the sub-Saharan African pair of YRI and LWK (*p* < 10^−6^, *ν* =1; previously identified in YRI; see Figure 6) and at *P4HA1* between the European CEU and South Asian GIH populations (*p* < 10^−6^, *ν* =1; previously identified in GIH, though as a soft sweep). We also observed a dearth of high-magnitude negative values in Tables S2-S10, with prominent convergent sweep candidates only occurring between the most diverged population pairs. These consisted of *C2CD5* between CEU and the East Asian JPT population (*ν* = 1), *PAWR* between Indo-European populations CEU and GIH with the sub-Saharan African YRI population (significant for the GIH-YRI comparison, *p* = 2 × 10^−6^, *ν* = 1 for both comparisons; Table S7), and *MPHOSPH9* and *EXOC6B* between JPT and YRI (both significant with p = 10^−6^ and *ν* =1). These observations reflect the broader pattern that negative SS-H12 values are rare between closely-related populations. Indeed, the majority of SS-H12 values at protein-coding genes between populations from the same geographic region are positive, and this distribution shifts toward negative values for more differentiated population pairs, consisting primarily of intermediate-magnitude negative values between the YRI and non-African populations (Figure S14). Our present results are also consistent with the H12-based observations of Harris et al. [2018] in single populations, in that we found a greater proportion of hard sweeps than soft sweeps among outlying sweep candidates in humans, though both were present between all population pairs.

**Figure 6:**
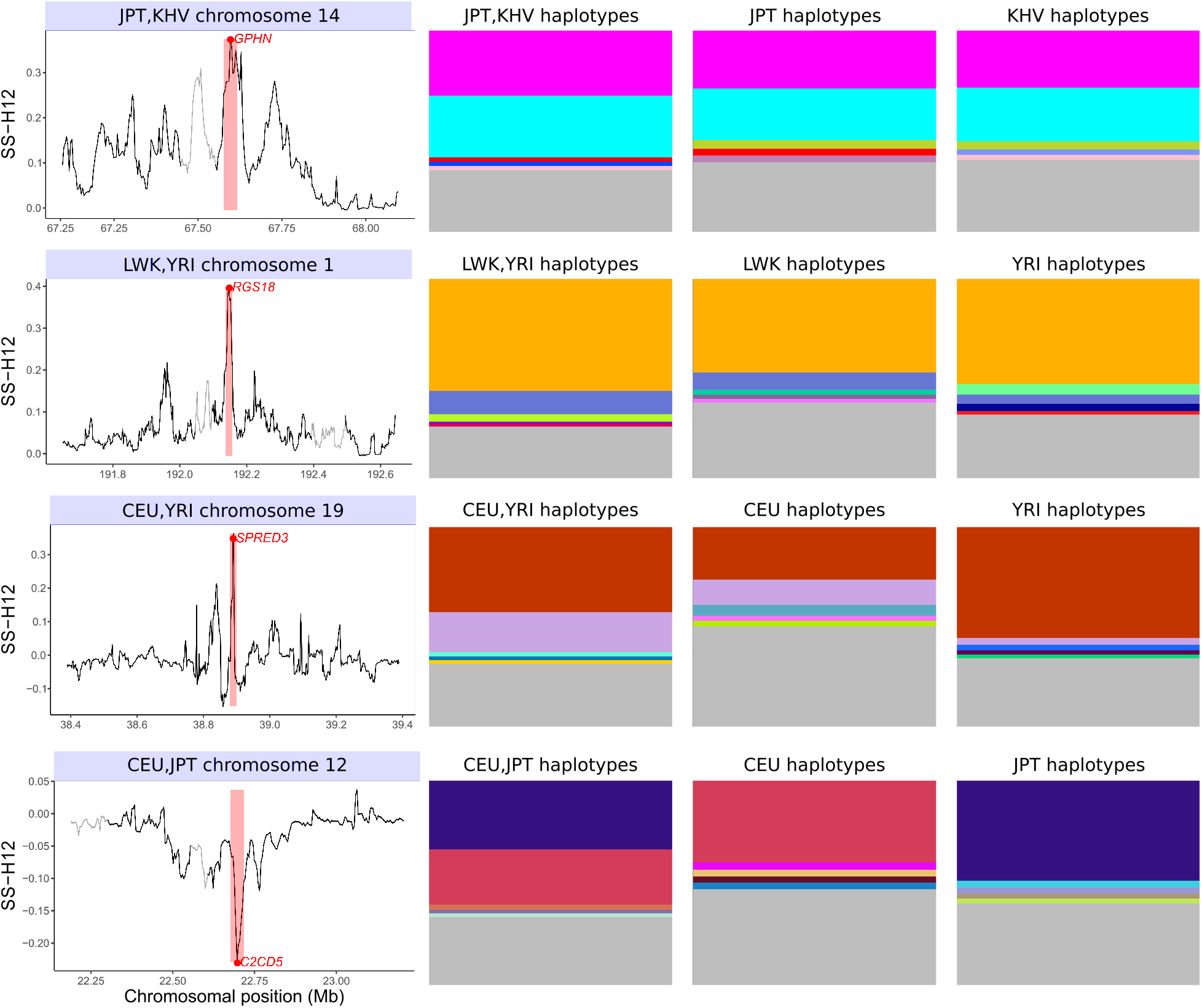
Top outlying shared sweep candidates at RNA- and protein-coding genes in global human populations. The signal peak, including chromosomal position, magnitude, and highlighted window of maximum SS-H12 (left column), as well as the haplotype frequency spectrum (right three columns) are displayed for each candidate. The East Asian JPT and KHV populations experience an ancestral soft sweep at *GPHN* (top row). The sub-Saharan African populations LWK and YRI share an ancestral hard sweep at *RGS18* (second row). The European CEU population experiences a shared sweep with YRI at *SPRED3* (third row). The European CEU and East Asian JPT have a convergent sweep at *C2CD5*, with a different, single high-frequency haplotype present in each population (bottom row). Coloration within the haplotype frequency spectrum plots indicates particular moderate to high-frequency haplotypes, while rows shaded in gray represent the remainder of haplotypes that are each at low frequency.

Our top shared sweep candidates also comprised genes that have been described in greater detail in the literature [Bersaglieri et al., 2004, Sabeti et al., 2007, Gerbault et al., 2009, Liu et al., 2013], including *LCT* and the surrounding cluster of genes on chromosome 2 including *MCM6, DARS*, and *R3HDM1* in the European CEU-GBR pair (*ν* =1 for all; Table S2), reflecting selection for the lactase persistence phenotype. We also recovered the sweep on the light skin pigmentation phenotype in Indo-Europeans [Sabeti et al., 2007, Coop et al., 2009, Mallick et al., 2013, Liu et al., 2013] for comparisons between the CEU population with GBR (Table S2; near-significant with *p* = 4 × 10^−6^ and *ν* =1) and GIH (Table S3; *p* < 10^−6^, *ν* = 1). Although the selected allele for this sweep is thought to lie within the *SLC24A5* gene encoding a solute carrier [Lamason et al., 2005], the mappability and alignability filter that we applied to our data removed *SLC24A5*, but preserved the adjacent *SLC12A1*, which we use as a proxy for the expected signal. Finally, we find *KIAA0825* as a top candidate across comparisons between the CEU and GIH (Table S3; *p* = 2 × 10^−6^, *ν* = 1), YRI and CEU (Table S5; *p* = 3 × 10^−6^, *ν* = 1), YRI and LWK (Table S6; *p* < 10^−6^, *ν* = 1), JPT and YRI (Table S8; *p* < 10^−6^, *ν* =1), and GIH and YRI (Table S7; *p* < 10^−6^, *ν* =1) populations. Although the function of *KIAA0825* has not yet been characterized, it is a previously-reported sweep candidate ancestral to the split of African and non-African human populations [Racimo, 2016].

Across all population comparisons, the top shared sweep candidates at RNA- and protein-coding genes comprised both hard and soft sweeps, yielding a wide range of H2_Tot_/H1_Tot_ values. This emphasizes the multitude of sweep histories that have shaped shared variation among human populations. In Figure 6, we highlight four distinct results that capture the diversity of sweeps we encountered in our analysis. We first examine *GPHN*, which we found as an outlying candidate shared soft sweep in the East Asian JPT and KHV populations (*ν* = 2; Table S10). *GPHN* encodes the scaffold protein gephyrin, which has been the subject of extensive study due to its central role in regulating the function of neurons, among the many other diverse functions of its splice variants [Ramming et al., 2000, Lencz et al., 2007, Tyagarajan and Fritschy, 2014]. *GPHN* has received attention as the candidate of a recent selective sweep ancestral to the human out-of-Africa migration event [Voight et al., 2006, Williamson et al., 2007, Park, 2012], which has resulted in the maintenance of two high-frequency haplotypes worldwide [Climer et al., 2015]. Although not meeting the genome-wide significance threshold, we see that a large signal peak is centered over *GPHN*, and the underlying haplotype structure shows two high-frequency haplotypes at similar frequency in the pooled population and in the individual populations (Figure 6, top row).

Next, we recovered *RGS18* as a significant novel outlying ancestral sweep signal in the sub-Saharan African LWK and YRI populations. *RGS18* occurs as a significant sweep in the YRI population [Harris et al., 2018] and correspondingly displays a single shared high-frequency haplotype between the LWK and YRI populations (Figure 6, second row), matching our assignment of this locus as a hard sweep (*ν* = 1). *RGS18* has been implicated in the development of hypertrophic cardiomyopathy, a leading cause of sudden cardiac death in American athletes of African descent [Maron et al., 2003, Chang et al., 2007]. Between the CEU and YRI populations, we found another novel shared sweep at *SPRED3* (Figure 6, third row; *p* = 2 × 10^−6^, *ν* =1), which encodes a protein that suppresses cell signaling in response to growth factors [Kato et al., 2003]. Although elevated levels of observed homozygosity at this gene have previously been reported in European and sub-Saharan African populations separately [Granka et al., 2012, Ayub et al., 2013], these observations have not previously been tied to one another. While both the CEU and YRI populations share their most frequent haplotype in an ancestral sweep, we found that this haplotype is at much lower frequency in the CEU than in the YRI, potentially indicating differences in the strength and duration of selection between these populations at *SPRED3* after their split in the out-of-Africa event.

Finally, we present the novel convergent hard sweep candidate that we uncovered at *C2CD5* (also known as *CDP138*) between the CEU and JPT populations. As expected of a convergent sweep, the signal peak here is large in magnitude but negative, corresponding to the presence of a different high-frequency haplotype in each population, each of which is also at high frequency in the pooled population (Figure 6, bottom row). The protein product of *C2CD5* is involved in insulin-stimulated glucose transport [Xie et al., 2011, Zhou et al., 2018], and the insulin response is known to differ between European and East Asian populations [Kodama et al., 2013]. Therefore, our discovery of *C2CD5* is in agreement with the results of Kodama et al. [2013], and illustrates the importance of differentiating ancestral and convergent sweeps in understanding the adaptive histories of diverse populations. We also highlight our discovery of *PAWR* as another outlying novel convergent hard sweep candidate with complementary clinical support, for comparisons between GIH and CEU with YRI. The protein product of *PAWR* is involved in promoting cancer cell apoptosis, and is implicated in the development of prostate cancer [Yang et al., 2013]. Because mutations within and adjacent to *PAWR* have been specifically implicated in the development of prostate cancer among individuals of African descent [Bonilla et al., 2011], our identification of a candidate convergent sweep at *PAWR* is consistent with the observation of elevated prostate cancer rates for populations with African ancestry [Kheirandish and Chinegwundoh, 2011, Shenoy et al., 2016].

## Discussion

Characterizing the selective sweeps shared between geographically close and disparate populations can provide insights into the adaptive histories of these populations that may be unavailable when analyzing single populations separately. To this end, we extended the H12 framework of Garud et al. [2015] to identify genomic loci affected by selection in samples composed of individuals from two or more populations. Our approach, SS-H12, has high power to detect recent shared selective sweeps from phased haplotypes, and is sensitive to both hard and soft sweeps. SS-H12 can also distinguish hard and soft sweeps from one another in conjunction with the statistic H2_Tot_/H1_Tot_, thus retaining the most important feature of the single-population approach. Furthermore, SS-H12 has the unique ability to distinguish between sweeps that are shared due to common ancestry (ancestral sweeps), and shared due to independent selective events (convergent sweeps). Analysis with the SS-H12 framework therefore provides a thorough characterization of selection candidates, both previously-described and novel, unlike that of other methods for detecting shared sweeps.

Because the value of SS-H12 fundamentally derives from a measure of expected haplotype homozygosity, it is tailored toward the detection of recent shared selective sweeps. Accordingly, we found that in our simulated shared sweep experiments, the power of SS-H12 to distinguish sweeps from neutrality was greatest for selective events beginning between 400 and 3000 generations prior to the time of sampling, assuming demographic parameters based on human values (Takahata et al. [1995], Nachman and Crowell [2000], Payseur and Nachman [2000], Terhorst et al. [2017], Narasimhan et al. [2017]; Figures 2, 3, and S2-S9). Due to their greater distortion of the haplotype frequency spectrum, stronger sweeps yield larger values of SS-H12 and larger sweep footprints than do weaker sweeps [Gillespie, 2004, Garud et al., 2015, Hermisson and Pennings, 2017]. Correspondingly, we could detect stronger sweeps over a wider range of selection times than weaker sweeps (Figures 2, 3, and S2-S5). However, stronger sweeps reach fixation sooner than do weaker sweeps, and their signal therefore begins to erode sooner than that of weaker sweeps, especially for sweeps from larger *ν* (compare, for example, Figures 2, S2, and S3 for ancestral sweeps). This illustrates the inverse correlation between the strength of sweeps (*s*) that SS-H12 can detect, and the range of selection start times (*t*) for which SS-H12 can detect a sweep. Harris et al. [2018] also observed this effect for the single-population approach.

The timing and strength of a shared sweep were important not only for detecting the sweep, but also for classifying it as ancestral or convergent. Barring the rare occurrence of a convergent sweep on the same haplotype between sister populations (which occurred for a minority of replicates across our tested parameters; Figures 2, 3, and S2-S9), we found that simulated convergent sweeps could reliably be identified from the sign of SS-H12 under scenarios in which SS-H12 has power to detect sweeps shared between population pairs. For ancestral sweeps, however, it was possible for negative values of elevated magnitude to emerge for weaker sweeps, especially if the time of selection *t* was close to the split time *τ* for the CEU-GIH model (bottom row of Figures 2, S2, and S3), or for the CEU-YRI model in general (bottom row of Figures 3, S4, and S5). In the former case, it is likely that the beneficial allele, and the haplotypic background(s) on which it resides, have not risen to high frequency before the ancestral population splits into the modern sampled populations. In the latter case, enough time has passed since *τ* by the time of sampling that extensive population differentiation can arise. Thus, in both cases, copies of the beneficial allele present in each of the two descendant populations may follow distinct trajectories. Using a smaller analysis window may therefore increase power to detect sweeps with less prominent footprints, but at the risk of misinterpreting elevated SS-H12 due to short-range LD as a sweep signal.

More generally, the ability of SS-H12 to identify a shared sweep as ancestral or convergent depends upon the underlying phylogeny of the sampled populations. For both simulated strong and moderate shared sweeps and either the CEU-GIH or CEU-YRI model, the power of SS-H12 was greatest for more recent sweeps. For two sampled populations related as in our CEU-GIH model (*τ* = 1100, shared ancestral bottleneck), SS-H12 can detect a wide range of both ancestral and convergent selective sweeps (Figures 2, S2, and S3). However, SS-H12 has little power to detect ancestral sweeps for *τ* > 3000 because the sweep signal erodes before the time of sampling, and this is compounded by the relative difficulty of detecting sweeps under a bottleneck demographic history. The limitations on *τ* for SS-H12 under the CEU-YRI history are similar, but power extends to older sweeps due to the higher background diversity of samples containing sub-Saharan African individuals (Figures 3, S4, and S5). Whereas the majority of outlying strong sweeps were convergent in either model, detectable moderate sweeps were more likely to be ancestral, as the peak in power for SS-H12 occurred for sweeps older than *τ* under smaller *s*. In practice, we found that most outlying shared sweep candidates between pairs of human populations were ancestral (SS-H12 > 0; Tables S2-S10), indicating that despite the power of our approach to detect convergent sweeps, such events may simply be uncommon because beneficial mutations are rare [Orr, 2010], and so the independent establishment of a beneficial mutation at the same locus across multiple populations should be especially rare for all but the most strongly-selected mutations [Haldane, 1927, Kimura, 1962, Wilson et al., 2014].

Consistent with results from the single-population approach [Garud et al., 2015, Harris et al., 2018], SS-H12 has power to detect shared soft sweeps, and can assign these as ancestral or convergent similarly to shared hard sweeps (Figures S2-S5). Our results for soft sweeps indicate, as expected, that increasing the number of sweeping haplotypes *ν* decreases the power of SS-H12 to detect shared sweeps, particularly strong sweeps. Sweeps from larger *ν* produce smaller distortions in the haplotype frequency spectrum relative to hard sweeps, yielding smaller values of |SS-H12| that are less likely to be distinct from neutrality. Our soft sweep simulations nonetheless consistently maintained similar distributions of SS-H12 to our hard sweep simulations (Figures 2, 3, and S2-S5), indicating that all haplotypes need not be shared between sampled populations, or at similar frequencies, in order to yield outlying SS-H12 signatures. That is, our simulated population split events represented a random sampling of haplotypes from the ancestor, which did not guarantee identical haplotype frequency spectra between descendant sister populations, or between the descendants and the ancestor, and SS-H12 could still identify ancestral shared sweeps with high power. This matches what we observed in the empirical data, wherein the frequencies of shared haplotypes at multiple top candidates differed considerably between populations. Our results for *SPRED3* between CEU and YRI provide a representative example of this (Figure 6, third row). While the H12 signal at *SPRED3* in the CEU population is not strong, there is enough overlap in its shared haplotype with YRI to identify it as a significant ancestral sweep. Thus, our simulated and empirical results suggest that SS-H12 is robust to deviations in haplotype frequency spectra between populations as long as sufficient haplotype sharing still persists.

Our application of SS-H12 to simulated samples composed of individuals from *K* ∈ {3, 4, 5} populations and a simple constant demographic history model with strong selection (*s* = 0.1; Figures S7-S9) demonstrated that our approach maintains consistent power to detect recent shared sweeps regardless of sample structure, and does not deviate from results for *K* = 2 sampled populations (Figure S6). Across all experiments, power curves were nearly identical to one another, with high power for strong hard sweeps starting within *t* ∈ [200, 1500] generations prior to sampling. Furthermore, the conservative and grouped approaches provided comparable power to one another, indicating the validity of either strategy. However, both the conservative and grouped approaches could not consistently classify convergent sweeps, frequently assigning a positive SS-H12 for convergent sweeps initiated before the most recent split time (*i.e*., predating *τ*_*K*−1_). In the case of the conservative approach, this is because we select the smallest-magnitude SS-H12 between pairs of populations as the SS-H12 for the whole sample, regardless of sign. Because the magnitude of SS-H12 is smaller for ancestral sweeps than for convergent sweeps occurring at the same *t*, an ancestral SS-H12 is often assigned if the convergent sweep predates at least one coalescence event between sampled populations, reflecting an internal ancestral sweep within the phylogeny. Accordingly, the distribution of SS-H12 values for *K* = 3 (Figure S7; *τ*_2_ = 750) and *K* = 4 (Figure S8; *τ*_3_ = 500) population samples and *t* = 400 is primarily centered on negative values, but we obtained many positive outliers for *K* = 5 (Figure S9; *τ*_4_ = 250). The grouped approach was overall more sensitive than the conservative to the presence of internal ancestral sweeps. This is likely the effect of the asymmetric topology of the population phylogeny we examined here, as older convergent sweeps result in a greater number of populations sharing the sweep ancestrally. For example, a convergent sweep at *t* = 800 in the *K* = 5 scenario (Figure S9) results in a sweep ancestral to the split of four sampled populations, which comprise 80% of the sample. Thus, a single high-frequency haplotype predominates, allowing for values of H12_Tot_ larger than *f*_Diff_, and SS-H12 > 0. Because all ancestral sweeps are shared identically across all populations, ambiguity in their classification does not occur, and their SS-H12 distributions match those of *K* = 2 scenarios.

Across our tested simulation scenarios, SS-H12 assigned only values of small magnitude to divergent sweeps, eliminating them as potentially-outlying signals in our analyses. This feature is important because sweeps in one population lead to differentiated haplotype frequency spectra between populations, providing values of *f*_Diff_ that may spuriously resemble convergent sweeps. We successfully dampened the signal of divergent sweeps through the application of a correction factor that prevents sweeps unshared among sampled populations from generating outlying values of SS-H12 (Equation 3), regardless of the number of sampled populations *K*. Indeed, the distribution of SS-H12 values generated under divergent sweeps often appears visually no different from neutrality, leaving no appreciable power to the method and no possibility of misidentifying divergent sweeps as shared sweeps (Figures 2, 3, and S2-S5). Although divergent sweeps were never prominent for samples composed of *K* > 2 populations, we note that such sweeps may spuriously show elevated power as more populations share the sweep ancestrally (Figures S7-S9), causing the distribution of SS-H12 values in the sample to become distinct from those under neutrality and shared sweeps alike.

SS-H12 also displayed an extensive robustness to admixture as a confounding factor in shared sweep detection and classification, allowing for the confident application of our approach to a wider set of complex demographic scenarios (Figures 4 and S12). We focused primarily on admixture as a confounding factor because it has been widely documented in the literature across multiple study systems, and is therefore an important consideration in many genome-wide analyses [Chun et al., 2010, Patterson et al., 2012, Pool et al., 2012, Nedić et al., 2014]. For our simulated study scenarios, we sought to model admixture events likely to produce non-negligible distortions in the haplotypic diversity of the sample. We chose as the admixing population a donor that was highly diverged from the sampled populations, splitting one coalescent unit (20,000 generations) before sampling, and allowed for high admixture proportions, up to 40%. In this way, admixture would likely introduce new haplotypes into the sample.

Admixture most impacted the ability of SS-H12 to detect and classify ancestral sweeps, whereas convergent sweeps remained broadly unobscured and distinct from neutrality for all but the most extreme scenarios. This fit our expectations because admixture into one population within a sampled pair sharing an ancestral sweep leads to differing haplotype frequency spectra between the pair, thus spuriously resembling a convergent sweep if donor genetic diversity is low, or neutrality otherwise (Figure 4, middle column). Accordingly, convergent sweeps may still appear convergent in many cases following admixture (Figure 4, left column). In most cases, the effect of admixture is likely to be an overall reduction in the prominence of SS-H12 at sweep loci, which may impact estimates of sweep age and intensity [Malaspinas et al., 2012, Mathieson and McVean, 2013, Smith et al., 2018], but not detection (Figure S12). Once again, the SS-H12 distribution of divergent sweeps showed little departure in prominence from neutrality (Figure 4, right column)—though with a spurious but not impactful rise in power (Figure S12, right column)—but this could be because only the sweeping population received gene flow from the donor population, resulting in little change to the correction factor (Equation 3) relative to no admixture. We caution that admixture from a donor of small size into the non-sweeping population may increase H12 in that population, potentially increasing the value of the correction factor, and SS-H12. Ultimately, only a narrow range of admixture scenarios is likely to affect inferences with SS-H12, and admixture was the only confounding factor we tested that affected SS-H12.

Beyond detecting and classifying recent shared sweeps with high power, accuracy, and specificity, the SS-H12 framework provides the only approach among comparable methods that can classify shared sweeps as hard or soft from the inferred number of sweeping haplotypes (*ν*). We based our shared sweep classification strategy on the approximate Bayesian computation (ABC) approach of Harris et al. [2018], leveraging the observation that shared sweeps yield differing paired (|SS-H12|, H2_Tot_/H1_Tot_) value profiles based on the underlying *ν* parameter. Using an ABC approach, we found that the classification of recent ancestral sweeps broadly followed that of sweeps in single populations, with smaller H2_Tot_/H1_Tot_ corresponding to harder sweeps, and the largest *ν* associated with the largest H2_Tot_/H1_Tot_ (Figure 5, top-left). Because we must constrain ancestral sweeps such that *t* > *τ*, the range of possible SS-H12 values is reduced relative to convergent sweeps, and the boundaries among *ν* classes are somewhat irregular within the posterior distribution. Resolving the most probable *ν* is therefore more challenging for ancestral sweeps (Figure 5). In contrast, the classification of convergent sweeps as hard or soft is both easier and more regular than is the classification of ancestral sweeps, because they will always be more recent and therefore produce a stronger sweep signal on average (Figure 5, bottom). Hard convergent sweeps occupy the largest |SS-H12| values, and pair with intermediate H2_Tot_/H1_Tot_ values, reflecting the presence of two independently-sweeping haplotypes in the pooled population. Sweeps on *ν* ≥ 2 haplotypes can only produce smaller |SS-H12| values, and sweeps from the largest *ν* once again occupy the largest H2_Tot_/H1_Tot_ values. Thus, the SS-H12 framework is powerful for classifying the softness of recent shared sweeps, provided that they are sufficiently recent, losing resolution proportionally to the power trends we observed across Figures 2, 3 and S2-S9.

Because phased haplotype data are often unavailable for non-model organisms, we also evaluated the power of our shared sweep approach applied to diploid unphased multilocus genotypes (MLGs), which we generated by manually merging individuals’ haplotypes in existing simulated samples. Previously, Harris et al. [2018] showed that the single-population approach H12 [Garud et al., 2015] could be applied to MLGs as the analogous statistic G123, which pools the three most frequent MLG frequencies as 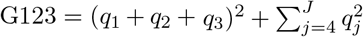, where there are *J* distinct unphased MLGs, with *q_j_* denoting the frequency of the *j*th most frequent MLG, and with *q*_1_ ≥ *q*_2_ ≥ ⋯ ≥ *q_J_*. We construct a MLG analogue of SS-H12, denoted SS-G123, as 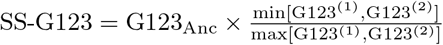, where G123^(1)^ and G123^(2)^ are G123 computed in populations 1 and 2, respectively, where G123_Anc_ = G123_Tot_ − *g*_Diff_, where 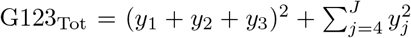 and where 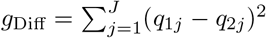, assuming that *y_j_* = *γq*_1*j*_ + (1 − *γ*)*q*_2*j*_, *y*_1_ ≥ *y*_2_ ≥ ⋯ ≥ *y_J_*, is the frequency of the *j*th most frequent MLG in the pooled population, and *q*_1*j*_ and *q*_2*j*_ are the frequencies of this MLG in populations 1 and 2, respectively. We can also classify shared sweeps as hard or soft using MLG data in the same manner as for haplotypes by pairing |SS-G123| and G2_Tot_/G1_Tot_ values, where 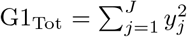 and 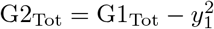, and where larger values of G2_Tot_/G1_Tot_ are once again associated with softer sweeps.

Overall, SS-G123 performed comparably to SS-H12 across identical CEU-GIH and CEU-YRI scenarios (Figures S15-S20), with slight reductions in power at the 1% and more so at the 5% FPRs for MLGs relative to haplotypes. Reductions in power generally occurred for older sweep times, as MLGs are more diverse than haplotypes [Harris et al., 2018], and so the signal of a sweep erodes more rapidly for MLG data as mutation and recombination events accumulate. Under both the CEU-GIH and CEU-YRI models, we found that the magnitude of SS-G123 was generally smaller than the magnitude of SS-H12, matching trends from results with the single-population statistics H12 and G123 [Harris et al., 2018]. Thus, because the power of SS-G123 is on par with that of SS-H12, we expect that the detailed analysis of shared selective sweeps will be possible for a wide variety of organisms for which whole-genome sequence data are available, and consistent with analyses using haplotype data, without introducing the uncertainty associated with phasing. To this end, we have included a parallel analysis of the 1000 Genomes Project dataset [Auton et al., 2015] in which we paired individuals’ haplotypes into their MLGs (see subsequent *Discussion*).

The high power, robustness, and flexibility of SS-H12 allowed us to discover outlying sweep candidates in humans that both corroborated the results of previous investigations, and uncovered previously-unknown shared sweep candidates. Most importantly, our SS-H12 framework provided inferences about the timing and softness of shared sweeps, yielding enhanced levels of detail about candidates that were until now not directly available. As SS-H12 is the only method that distinguishes between recent ancestral and convergent shared sweeps, our investigation was uniquely able to identify loci at which independent convergent sweeps, though rare, may have played a role in shaping modern patterns of genetic diversity. Among these loci was *EXOC6B*, which produces a protein component of the exocyst [Evers et al., 2014] and has been previously highlighted as a characteristic site of selection in East Asian populations [Baye et al., 2009, Durbin and Consortium, 2011, Pybus et al., 2014]. The shared hard sweep (*ν* = 1) at *EXOC6B* appeared as convergent between the East Asian JPT and sub-Saharan African YRI populations (Table S8), but as ancestral between all other population pairs—pairs of non-African populations—in which it appeared (Tables S3, S4 S9, and S10). Thus, we have identified the sweep at *EXOC6B* as a global occurrence that affected African and non-African populations alike, and was not limited to a single region or a single event.

More generally, our investigation into sweep signals shared between disparate population pairs also updates existing notions about when during the peopling of the world particular selective events may have occurred. For example, a sweep at *NNT*, involved in the glucocorticoid response, has been previously reported in sub-Saharan African populations [Voight et al., 2006, Fagny et al., 2014]. As expected, we recovered *NNT* as a significant ancestral hard sweep (*p* < 10^−6^, *ν* = 1) in the comparison between LWK and YRI (Table S6), but additionally in all comparisons between YRI and non-African populations (Tables S5, S7, and S8; significant for all but the CEU-YRI pair). This indicates that selection at *NNT* would have preceded the out-of-Africa event and would not have been exclusive to sub-Saharan African populations. Another candidate that appeared as a top outlier outside of previously-described populations was *SPIDR*, involved in double-stranded DNA break repair [Wan et al., 2013, Smirin-Yosef et al., 2017] and inferred to be a shared candidate among Eurasian populations [Racimo, 2016]. *SPIDR* previously appeared as an outlying H12 signal in the East Asian CHB population [Harris et al., 2018], but in our present analysis was shared ancestrally not only between the East Asian KHV and JPT populations (Table S10; *p* = 10^−6^), but also between JPT and the European CEU (Table S4; *p* < 10^−6^), and the sub-Saharan African LWK and YRI (Table S6; *p* < 10^−6^) populations. Once again, we found evidence of a strong sweep candidate shared among a wider range of populations than previously expected, illustrating the role of shared sweep analysis in amending our understanding of the scope of sweeps in humans worldwide.

In addition to recovering expected and expanded sweep signatures across multiple loci, we also found top outlying ancestral shared sweep candidates whose signals are not especially prominent across single-population analyses, emphasizing that localizing an ancestral sweep depends not only on elevated expected homozygosity generating the signal, but highly on the presence of shared haplotypes between populations to maintain the signal. Foremost among such candidates was *CASC4*, which appeared as an ancestral hard sweep (*ν* = 1) in all comparisons with YRI (Tables S5-S8; significant for all but the CEU-YRI pair). Because this cancer-associated gene [Ly et al., 2014, Anczuków et al., 2015] had been previously described as a shared sweep ancestral to the out-of-Africa event [Racimo, 2016], we expected to see it represented across multiple comparisons. However, *CASC4* does not have a prominent H12 value outside of sub-Saharan African populations, and within YRI is a lower-end outlier [Harris et al., 2018]. Despite this, *CASC4* is within the top 12 outlying candidates across all comparisons with YRI, and appears as the eighth-most outlying signal in the CEU-JPT comparison (Table S4, *p* = 10^−6^, *ν* = 2), even though it does not appear as an outlier in either population individually. Similarly, we found *PHKB*, involved in glycogen storage [Hendrickx and Willems, 1996, Burwinkel et al., 1997, Burwinkel and Kilimann, 1998], as an ancestral hard sweep between the CEU and YRI populations (Table S5; *ν* = 1) that was not prominent in either population alone, though it had been previously inferred to be a sweep candidate ancestral to Eurasian populations once again [Racimo, 2016]. We also identified *MRAP2*, which encodes a melanocortin receptor accessory protein that is implicated in glucocorticoid deficiency [Chan et al., 2009, Asai et al., 2013], similarly to *NNT*, as an ancestral hard sweep between the CEU and JPT populations (Table S4; *p* < 10^−6^, *ν* = 1), and is not prominent in either CEU or JPT. Thus, our empirical results fit well with the expectation deriving from our power comparison between multiple tests of H12 and a single SS-H12 test (Figures S10 and S11), which suggested that SS-H12 can detect shared sweeps that may go unnoticed in separate analyses of populations under selection.

We supplemented our empirical analyses of human data [Auton et al., 2015] by performing parallel scans of sampled individuals’ MLGs (manually merged from their haplotypes) to evaluate the ability of SS-G123 to yield the same insights as SS-H12 for the same data. Overall, we generated congruent lists of top sweep candidate genes between haplotype (Tables S2-S10) and MLG (Tables S11-S19) analyses in terms of both inclusion and significant signals of candidate genes. The primary difference that we encountered between haplotype and MLG analyses was in the inferred softness of candidate sweeps. We found that, as in the single-population analyses of Harris et al. [2018], a greater proportion of top candidate sweeps in the MLG data were classified as soft than in haplotype data, including both candidates classified as hard sweeps in the haplotype data, and candidates absent from the top 40 haplotype candidates. The explanation for both of these discrepancies, which were minor in scope, lies in the greater diversity of MLGs relative to haplotypes. A genomic region with one high-frequency haplotype and one or more intermediate-frequency haplotypes may yield a paired (|SS-H12|, H2_Tot_/H1_Tot_) value that most resembles a hard sweep under the ABC approach using haplotypes, but yield a MLG frequency spectrum featuring multiple intermediate-frequency MLGs that may be inferred as a soft sweep. Meanwhile the greater background diversity of MLG data may allow for the more subtle signatures of soft sweeps to be more readily detectable than in haplotype data. Nonetheless, the rarity of discrepancies between SS-H12 and SS-G123 top candidate lists corroborates the high level of concordance between the two statistics that we found in simulated data (compare Figures 2, 3, and S2-S5 to Figures S15-S20).

The SS-H12 and SS-G123 frameworks represent an important advancement in our ability to contextualize and classify shared sweep events using multilocus sequence data. Whereas previous experiments have identified shared sweeps and can do so with high power, or without the need for MLGs or phased haplotypes, the ability to distinguish both hard and soft shared sweeps from neutrality, as well as differentiate ancestral and convergent sweeps, is invaluable for understanding the manner in which an adaptive event has proceeded. Discerning whether a selective sweep has occurred multiple times or only once can provide novel and updated insights into the relatedness of study populations, and the selective pressures that they endured. Moreover, the sensitivity of our approach to both hard and soft sweeps, and our ability to separate one from the other, add an additional layer of clarity that is otherwise missing from previous analyses, and is especially relevant because uncertainty persists as to the relative contributions of hard and soft sweeps in human history [Jensen, 2014, Schrider and Kern, 2017, Mughal and DeGiorgio, 2018]. Ultimately, we expect inferences deriving from SS-H12 analysis to assist in formulating and guiding more informed questions about discovered candidates across diverse organisms for which sequence data—phased and unphased—exist. After establishing the timing and softness of a shared sweep, appropriate follow-up analyses can include inferring the age of a sweep [Smith et al., 2018], identifying the favored allele or alleles [Akbari et al., 2018], or identifying other populations connected to the shared sweep. We believe that our approach will serve to enhance investigations into a diverse variety of study systems, and facilitate the emergence of new perspectives and paradigms.

Finally, we provide open-source software (titled `SS-X12`) to perform scanning window analyses on haplotype input data using SS-H12 or multilocus genotype input data using SS-G123, as well as results from our empirical scans, at http://personal.psu.edu/mxd60/SS-X12.html. `SS-X12` provides flexible user control, allowing the input of samples drawn from arbitrary populations *K*, and the output of a variety of expected homozygosity summary statistics.

## Materials and Methods

We first tested the power of SS-H12 to detect shared selective sweeps on simulated multilocus sequence data, including both phased haplotypes and unphased multilocus genotypes (MLGs; applied as SS-G123). We generated haplotype data using the forward-time simulator SLiM 2 [version 2.6; Haller and Messer, 2017], which follows a Wright-Fisher model [Fisher, 1930, Wright, 1931] and can reproduce complex demographic and selective scenarios. We generated MLGs from haplotypes by manually merging each simulated individual’s pair of haplotypes into a single MLG. In this way, we were able to directly assess the effects of phasing on identical samples. This series of simulations followed human-inspired parameters [Takahata et al., 1995, Nachman and Crowell, 2000, Payseur and Nachman, 2000, Terhorst et al., 2017, Narasimhan et al., 2017], with mutation rate *μ* = 1.25 × 10^−8^ per site per generation, recombination rate drawn at random from an exponential distribution with mean *r* = 10^−8^ per site per generation and maximum truncated at *r* = 3 × 10^−8^ [Schrider and Kern, 2017, Mughal and DeGiorgio, 2018], and two sampled populations whose sizes we inferred using `smc++` [Terhorst et al., 2017] from whole genome polymorphism data [Auton et al., 2015]. The populations in our model were the CEU of European descent (sample size *n* = 99 diploids) compared with either the GIH (*n* = 103) of South Asian descent (split time between populations *τ* = 1100 generations before sampling), or the YRI (*n* = 108) of sub-Saharan African descent (*τ* = 3740). As is standard for forward-time simulations [Yuan et al., 2012, Ruths and Nakhleh, 2013], we scaled these parameters by a factor λ = 20 to reduce simulation runtime, dividing the population size and duration of the simulation by λ, and multiplying the mutation and recombination rates, as well as the selection coefficient (*s*) by λ. Thus, scaled simulations maintained the same expected levels of genetic variation as would unscaled simulations.

Simulations generated under the aforementioned scheme lasted for an unscaled duration of 20*N* generations. This consisted of a burn-in period of 10*N* generations to produce equilibrium levels of variation in which the ancestor to the sampled modern populations was maintained at size *N* = 10^4^ diploids [Messer, 2013], and another 10*N* generations during which population size was allowed to change. We note that population split events occurred within the latter 10*N* generations of the simulation. Under this approach, we examined three broad classes of sweep scenarios, consisting of ancestral, convergent, and divergent sweeps. For ancestral sweeps, we introduced a selected allele to one or more randomly-drawn haplotypes in the ancestor of both sampled populations (*i.e*., before the population split), which ensured that the same selective event was shared in the histories of both populations. For convergent sweeps, we simultaneously introduced the selected mutation independently in each extant population at the time of selection, after the split had occurred. Finally, divergent sweeps comprised a scenario in which the sweep event occurred in one sampled population only, whereas the other did not experience a sweep by the time of sampling. Across all simulations, we conditioned on the maintenance of the selected allele in any affected population after its introduction. To generate distributions of SS-H12 and SS-G123 for power analysis, we scanned 100 kb of sequence data from simulated individuals using a sliding window approach. We computed both statistics in 20 (CEU-YRI) or 40 kb (CEU-GIH) windows, advancing the window by increments of one kb across the simulated chromosome for a total of 61 (CEU-GIH) or 81 (CEU-YRI) windows. For each replicate, we retained the value of SS-H12 or SS-G123 from the window of maximum absolute value as the score. We selected window sizes sufficiently large to overcome the effect of short-range LD in the sample, which may produce a signature of expected haplotype homozygosity resembling a sweep. This also matched our window sizes for empirical scans. We measured the decay of LD for SNPs in neutral replicates separated by one to 100 kb at one kb intervals using mean *r*^2^. For all parameter sets, we generated 10^3^ sweep replicates and 10^3^ neutral replicates between which population sizes, number of populations, and population split times were identical.

We additionally measured the ability of our approach to detect shared sweep events in simulated samples consisting of individuals from *K* = 2 to 5 populations, related by a rooted tree with *K* leaves, under a simpler demographic model in order to illustrate the effect of sampling multiple populations. SS-H12 and SS-G123 are compatible with an arbitrary number of *K* populations (see subsequent explanation). Under this model, population sizes remained at a constant *N* = 10^4^ diploids throughout the simulation, with mutation rate *μ* = 2.5 × 10^−8^ per site per generation, recombination rate *r* = 10^−8^ per site per generation, sample size of *n* = 100 diploids, and analysis window size of 40 kb. Thus, the scenarios we examined were equivalent except for the number of sampled populations. For multiple-population models, we define ancestral and convergent sweeps as previously, affecting all lineages and beginning either before any population splits, or after all population splits, respectively. Accordingly, a divergent sweep resulted from any positive selection event affecting fewer than all populations, meaning that at least one population did not experience a sweep during the simulation.

For simulated data consisting of individuals from *K* ≥ 3 sampled populations (which we analyze only for haplotype data), we assigned values of SS-H12 in one of two ways. First, we employed a conservative approach in which we computed the SS-H12 score for each possible population pair in the aforementioned manner, but retained as the replicate score only the SS-H12 value with the smallest magnitude. That is, the replicate score had to satisfy the condition |SS-H12_*K*≥3_| = min_*i*≠*j*_{|SS-H12_*ij*_|}, where SS-H12_*ij*_ is SS-H12 computed between populations *i* and *j*. Assigning SS-H12 in this manner ensured that only samples wherein all represented populations shared a sweep were likely to yield outlying values. Second, we explored a grouped approach in which we assigned the SS-H12 statistic between the two branches (denoted *α* and *β*) directly subtending the root of the phylogeny relating the set of *K* populations, treating the two subtrees respectively descending from these branches as single populations. Thus, 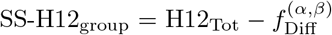, where H12_Tot_ is the expected haplotype homozygosity of the pooled population, and 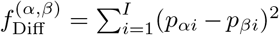, where *p_αi_* and *p_βi_* are the mean frequencies of haplotype *i* on branches *α* and *β*, respectively.

Across all tested population histories, we evaluated the ability of SS-H12 to identify hard selective sweeps from a *de novo* mutation and soft sweeps from selection on standing genetic variation, for both strong (*s* = 0.1) and moderate (*s* = 0.01) strengths of selection. This setting matched the experimental approach of Harris et al. [2018] for single-population statistics, and corresponds to scenarios for which those statistics have power under the specific mutation rate, recombination rate, and effective size we tested here. For all selection scenarios, we placed the beneficial allele at the center of the simulated chromosome, and introduced it at only one time point, constraining the selection start time, but not the selection end time. For hard and soft sweep scenarios, we allowed the selected allele to rise in frequency toward fixation (with no guarantee of reaching fixation). To specify soft sweep scenarios, we conditioned on the selected allele being present in the population on *ν* = 4 or 8 distinct haplotypes at the start of selection (*i.e*., 0.4 to 0.8% of haplotypes in the population would initially carry a beneficial allele), without defining the number of selected haplotypes remaining in the population at the time of sampling, as long as the selected allele was not lost.

For all method performance evaluation experiments, we observed the effect of varying the selection start time, and the time at which populations split from one another. To ensure that we evaluated a sufficiently broad spectrum of sweeps, we initiated selection at *t* ∈ [200, 4000] generations prior to sampling for the CEU-GIH and simplified models, and *t* ∈ [400, 6000] for the CEU-YRI model. This range of selection times was chosen not to represent specific selective events in human history, but to cover the range of time over which hypothesized selective sweeps in recent human history have occurred [Przeworski, 2002, Sabeti et al., 2007, Beleza et al., 2012, Jones et al., 2013, Clemente et al., 2014, Fagny et al., 2014]. We also varied split times (*τ*) across all experiments, with *τ* ∈ [250, 3740] generations before sampling. The combination of *t* and *τ* defined the simulation type. For two-population scenarios, simulations wherein *t* > *τ* produced ancestral sweeps, and simulations with *t* < *τ* yielded convergent or divergent sweeps depending on the number of populations under selection.

The number of population split events specified the number of populations in the simulated sample. For simulations in which the ancestral population split only once, at time *τ*, the sample consisted of *K* = 2 populations. To extend our notation for more than two populations, we index the divergence time as *τ_k_, k* = 1, 2,…, *K* − 1, for *K* populations. Similarly, if we included two population splits, at times *τ*_1_ and *τ*_2_ (where *τ*_1_ > *τ*_2_), then the sample consisted of individuals from *K* = 3 populations. For experiments involving *K* ≥ 3 sampled populations, we split populations at regularly repeating intervals, with each split generating a new population identical to its ancestor. We generated only asymmetric tree topologies, splitting each new population from the same ancestral branch. Furthermore, experiments with *K* ≥ 3 populations allowed for the simulation of more complex selection scenarios featuring nested ancestral sweeps, part of larger convergent or divergent sweep events, wherein the time of selection occurs between *τ*_1_ and *τ*_*K*−1_, with *τ*_1_ > *τ*_2_ > ⋯ > *τ*_*K*−1_ and *τ_k_* = *τ*_1_ − 250(*k* − 1) for *k* =1, 2,…, *K* − 1.

We additionally observed the effects of two confounding factors on SS-H12 to establish the extent to which inferences of shared sweeps in the sampled populations could be misled. For these experiments, we studied only scenarios with *K* = 2 populations. First, we examined the effect of admixture on one of the two sampled populations under the simplified demographic model. Second, we generated samples under longterm background selection, which is known to yield similar patterns of diversity to sweeps [Charlesworth et al., 1993, 1995, Seger et al., 2010, Nicolaisen and Desai, 2013, Cutter and Payseur, 2013, Huber et al., 2016], following the CEU-GIH and CEU-YRI models. Sample sizes *n* for each experiment remained identical to prior experiments for each model. For the admixture experiments, we simulated single pulses of admixture at fractions between 0.05 and 0.4, at intervals of 0.05, from a diverged unsampled donor, 200 generations prior to sampling (thus, following the start of selection in the sample). We simulated three different scenarios of admixture into the sampled target population from the donor population. These consisted of a highly-diverse donor population (*N* = 10^5^, tenfold larger than the sampled population), which may obscure a sweep signature in the sampled target, and from a low-diversity donor population (*N* = 10^3^, 1/10 the size of the sampled population), which may produce a sweep-like signature in the target, in addition to an intermediately-diverse donor population (*N* = 10^4^, equal to the size of the sampled population). For divergent sweep experiments, only the population experiencing the sweep was the target.

Our background selection simulations followed the same protocol as in previous work [Cheng et al., 2017]. At the start of the simulation, we introduced a centrally-located 11-kb gene composed of UTRs (5’ UTR of length 200 nucleotides [nt], 3’ UTR of length 800 nt) flanking a total of 10 exons of length 100 nt separated by introns of length one kb. Strongly deleterious (*s* = −0.1) mutations arose throughout the course of the simulation across all three genomic elements under a gamma distribution of fitness effects with shape parameter 0.2 at rates of 50%, 75%, and 10% for UTRs, exons, and introns, respectively. The sizes of the genic elements follow human mean values [Mignone et al., 2002, Sakharkar et al., 2004]. To enhance the effect of background selection on the simulated chromosome, we also reduced the recombination rate within the simulated gene by two orders of magnitude to *r* = 10^−10^ per site per generation.

As in Harris et al. [2018], we employed an approximate Bayesian computation (ABC) approach to demonstrate the ability of SS-H12, in conjunction with the H2_Tot_/H1_Tot_ statistic, to classify shared sweeps as hard or soft from the inferred number of sweeping haplotypes *ν*. Hard sweeps derive from a single sweeping haplotype, while soft sweeps consist of at least two sweeping haplotypes. Whereas the single-population approach [Garud et al., 2015, Garud and Rosenberg, 2015, Harris et al., 2018] identified hard and soft sweeps from their occupancy of paired (H12, H2/H1) values, we presently use paired (|SS-H12|, H2_Tot_/H1_Tot_) values to classify shared sweeps. We defined a 100 × 100 grid corresponding to paired (|SS-H12|, H2_Tot_/H1_Tot_) values with each axis bounded by [0.005, 0.995] at increments of 0.01, and assigned the most probable value of *ν* to each test point in the grid.

We define the most probable *ν* for a test point as the most frequently-observed value of *ν* from the posterior distribution of 5 × 10^6^ sweep replicates within a Euclidean distance of 0.1 from the test point. For each replicate, we drew *ν* ∈ {0, 2,…, 16} uniformly at random, as well as *s* ∈ [0.005, 0.5] uniformly at random from a log-scale. Across ancestral and convergent sweep scenarios for *K* = 2 sampled sister populations, we generated replicates for the CEU-GIH (*τ* = 1100) and CEU-YRI (*τ* = 3740) models. For ancestral sweeps, we drew *t* ∈ [1140, 3000] for CEU-GIH and *t* ∈ [3780, 5000] for CEU-YRI scenarios, both uniformly at random. Similarly, convergent sweeps were drawn from *t* ∈ [200, 1060] (CEU-GIH) and *t* ∈ [200, 3700] (CEU-YRI). Simulated haplotypes were of length 40 kb (CEU-GIH) or 20 kb (CEU-YRI), corresponding to the window size for method performance evaluations, because in practice a value of *ν* would be assigned to a candidate sweep based on its most prominent associated signal. Mutation and recombination rates, as well as sample size per population, were identical to previous experiments using these demographic models.

We applied SS-H12 and SS-G123 to human empirical data from the 1000 Genomes Project Consortium [Auton et al., 2015]. We scanned all autosomes for signatures of shared sweeps in population pairs using 40 kb windows advancing by increments of four kb for samples of non-African populations, and 20 kb windows advancing by two kb for any samples containing individuals from any African populations. We based these window sizes on the interval over which LD, measured as *r*^2^, decayed beyond less than half its original value relative to pairs of loci separated by one kb (Figure S21). As in Harris et al. [2018], we filtered our output data by removing analysis windows containing fewer than 40 SNPs, the expected number of SNPs corresponding to the extreme case in which a selected allele has swept across all haplotypes except for one, leaving two lineages [Watterson, 1975]. Following Huber et al. [2016], we also divided all chromosomes into non-overlapping bins of length 100 kb and assigned to each bin a mean CRG100 score [Derrien et al., 2012], which measures site mappability and alignability. We removed windows within bins whose mean CRG100 score was below 0.9, with no distinction between genic and non-genic regions. Thus, our overall filtering strategy was identical to that of Harris et al. [2018]. We then intersected remaining candidate selection peaks with the coordinates for protein- and RNA-coding genes from their hg19 coordinates.

Finally, we assigned *p*-values and the most probable inferred *ν* to the top 40 ancestral and convergent RNA- and protein-coding sweep candidates recovered in our genomic scans. We assigned *p*-values by generating 10^6^ neutral replicates in *ms* [Hudson, 2002] under the appropriate two-population demographic history, inferred from `smc++` [Terhorst et al., 2017]. Here, we drew the sequence length for each replicate uniformly at random from the set of all hg19 gene lengths and appended the window size, providing as input at least one full-length analysis window. The *p*-value for a selection candidate is the proportion of |SS-H12| values generated under neutrality whose value exceeds the maximum |SS-H12| assigned to the candidate. Following Bonferroni correction for multiple testing [Neyman and Pearson, 1928], a significant *p*-value was *p* < 0.05/23,735 ≈ 2.10659 × 10^−6^, where 23,735 is the number of protein- and RNA-coding genes for which we assigned a score.

We assigned the most probable *ν* for each sweep candidate following the same protocol as the constant-size demographic history simulation analyses, generating 5 × 10^6^ replicates of sweep scenarios generated in SLiM 2 under `smc++`-inferred demographic histories for ancestral and convergent sweeps, drawing *t* ∈ [200, 5000] uniformly at random, and *s* ∈ [0.005, 0.5] uniformly at random on a log scale. Once again, *t* > *τ* for ancestral sweep scenarios and *t* < *τ* for convergent sweep scenarios, where *τ* is defined by the specific demographic history of the sample. The CEU-GIH and CEU-YRI replicates used here were identical to those in Figure 5. Sequence length for each replicate was identical to analysis window length for equivalent empirical data (20 or 40 kb), because in practice we assign *ν* to windows of this size. For both *p*-value and most probable *ν* assignment, we used a per-site per-generation mutation rate of *μ* = 1.25 × 10^−8^ and a per-site per-generation recombination rate of *r* = 3.125 × 10^−9^ [Narasimhan et al., 2017, Terhorst et al., 2017].

**Table 1:** Summary of SS-H12 signals and their interpretation across various scenarios.

| Scenario | Sign of SS-H12 | Magnitude of SS-H12 | Comments | Reference |
| --- | --- | --- | --- | --- |
| Neutrality | Typically negative | Small | Magnitude becomes positive in bottleneck scenarios where the number of shared haplotypes between populations is higher by chance. | Figures 2-4 and S2-S9 |
| Ancestral sweep | Positive | Large | Magnitude is generally smaller than for convergent sweeps because ancestral sweeps are older; rare negative values may arise for weaker sweep strengths. | Figures 2-4 and S2-S9 |
| Convergent sweep | Predominantly negative | Large | Largest magnitude of SS-H12 across tested scenarios; positive values may arise in the rare event that two independent sweeps on the same haplotype occur between sampled populations. | Figures 2-4 and S2-S9 |
| Divergent sweep | Typically negative | Small | Trends in magnitude of SS-H12 match those of neutrality without exception; large magnitudes are impossible for divergent sweeps due to the correction factor (Equation 3). | Figures 2-4 and S2-S9 |
| Background selection | Typically negative | Small | Background selection has no discernible effect on the distribution of SS-H12 relative to neutrality. | Figure S13 |
| Admixture | Predominantly negative (see comments) | Small or large | Sufficient admixture from a diverse enough donor population will erode the signal of a sweep, yielding negative values of small magnitude; admixture with a low-diversity donor does not affect magnitude or signal of convergent sweeps, but will cause ancestral sweeps to spuriously resemble convergent sweeps. | Figures 4 and S12 |
| Number of sampled populations ($K$) | Negative or positive | Small or large | The number of populations included in the sample does not affect inference with SS-H12. | Figures S6-S9 |

## Acknowledgments

This work was funded by National Institutes of Health grant R35GM128590 and by the Alfred P. Sloan Foundation. Portions of this research were conducted with Advanced CyberInfrastructure computational resources provided by the Institute for CyberScience at Pennsylvania State University.

