## Supplemental Tables S1-S19; Supplemental Figures S1-S21 for "Identifying and classifying shared selective sweeps from multilocus data"

Table S1: Critical values of |SS-H12| and |SS-G123| generated from $10^6$ neutral replicates (identical for either statistic), used to assign $p$-values to empirical top candidates across 1000 Genomes Project [Auton et al., 2015] sampled population pairs.

| Population | SS-H12 | SS-G123 |
|---|---|---|
| CEU-GBR | 0.6151618 | 0.4017583 |
| CEU-GIH | 0.2755997 | 0.0863085 |
| CEU-JPT | 0.2800871 | 0.0717914 |
| CEU-YRI | 0.3924851 | 0.1569262 |
| LWK-YRI | 0.2065271 | 0.0395899 |
| GIH-YRI | 0.2701381 | 0.0582839 |
| JPT-YRI | 0.2408978 | 0.0450692 |
| JPT-GIH | 0.1956585 | 0.0351749 |
| JPT-KHV | 0.4114621 | 0.1926581 |

Table S2: Top 40 SS-H12 sweep candidates at RNA- and protein-coding genes shared between the Central and Western European CEU and GBR populations. Candidates presented are those that remained after application of a mappability and alignability filter to data (see *Materials and Methods*). Target genes that pass the significance threshold are colored in gold in the "$P$-value" column. Genes whose sweeps are assigned as hard ($\nu = 1$) are shaded in red in the "Inferred $\nu$" column, while soft sweeps ($\nu \geq 2$) are colored in blue.

| | Top gene | Chromosome | Maximum SS−H12 | H2/H1 | *P*−value | Inferred $\nu$ |
|---|---|---|---|---|---|---|
| 1 | *ZRANB3* | 2 | 0.6138516 | 0.005253369 | $2.0\times10^{-6}$ | 1 |
| 2 | *SLC12A1* | 15 | 0.5964931 | 0.007102656 | $4.0\times10^{-6}$ | 1 |
| 3 | *R3HDM1* | 2 | 0.5907587 | 0.005776011 | $4.0\times10^{-6}$ | 1 |
| 4 | *DARS* | 2 | 0.5605804 | 0.012398870 | $1.6\times10^{-5}$ | 1 |
| 5 | *MCM6* | 2 | 0.5604343 | 0.010608921 | $1.6\times10^{-5}$ | 1 |
| 6 | *LCT* | 2 | 0.5097962 | 0.009314534 | $8.9\times10^{-5}$ | 1 |
| 7 | *LOC100507600* | 2 | 0.5097962 | 0.009314534 | $8.9\times10^{-5}$ | 1 |
| 8 | *AC093391.2* | 2 | 0.4822492 | 0.008069683 | $3.14\times10^{-4}$ | 1 |
| 9 | *RAB3GAP1* | 2 | 0.4788360 | 0.016863242 | $3.48\times10^{-4}$ | 1 |
| 10 | *UBXN4* | 2 | 0.4709547 | 0.008006012 | $4.45\times10^{-4}$ | 1 |
| 11 | *BCAS3* | 17 | 0.4342505 | 0.024135621 | $1.43\times10^{-3}$ | 1 |
| 12 | *KAT6B* | 10 | 0.4063248 | 0.010954181 | $3.11\times10^{-3}$ | 1 |
| 13 | *PPM1D* | 17 | 0.4043315 | 0.013758902 | $3.27\times10^{-3}$ | 1 |
| 14 | *MYO9A* | 15 | 0.3919436 | 0.014969493 | $4.55\times10^{-3}$ | 1 |
| 15 | *FAM149B1* | 10 | 0.3841613 | 0.013332285 | $5.56\times10^{-3}$ | 1 |
| 16 | *ZNF546* | 19 | 0.3824392 | 0.053683499 | $5.80\times10^{-3}$ | 1 |
| 17 | *MAP3K19* | 2 | 0.3807448 | 0.175652783 | $6.06\times10^{-3}$ | 3 |
| 18 | *PRMT9* | 4 | 0.3782229 | 0.030300365 | $6.48\times10^{-3}$ | 1 |
| 19 | *KITLG* | 12 | 0.3772852 | 0.023892213 | $6.64\times10^{-3}$ | 1 |
| 20 | *MLLT3* | 9 | 0.3756959 | 0.085799301 | $6.92\times10^{-3}$ | 1 |
| 21 | *LAMA3* | 18 | 0.3749289 | 0.327190136 | $7.04\times10^{-3}$ | 3 |
| 22 | *TMEM116* | 12 | 0.3716065 | 0.023778920 | $7.60\times10^{-3}$ | 1 |
| 23 | *KMT2A* | 11 | 0.3684665 | 0.034335416 | $8.22\times10^{-3}$ | 1 |
| 24 | *ADRBK2* | 22 | 0.3648397 | 0.029916410 | $8.96\times10^{-3}$ | 1 |
| 25 | *ACMSD* | 2 | 0.3628647 | 0.058329766 | $9.40\times10^{-3}$ | 1 |
| 26 | *CCNT2−AS1* | 2 | 0.3628647 | 0.058329766 | $9.40\times10^{-3}$ | 1 |
| 27 | *PLAGL2* | 20 | 0.3588937 | 0.315722448 | $1.03\times10^{-2}$ | 3 |
| 28 | *DIRC3* | 2 | 0.3560424 | 0.021817335 | $1.10\times10^{-2}$ | 1 |
| 29 | *POLN* | 4 | 0.3534160 | 0.101220872 | $1.16\times10^{-2}$ | 1 |
| 30 | *AC005592.1* | 5 | 0.3518146 | 0.018718715 | $1.21\times10^{-2}$ | 1 |
| 31 | *PRKDC* | 8 | 0.3491810 | 0.262222086 | $1.28\times10^{-2}$ | 4 |
| 32 | *BVES−AS1* | 6 | 0.3486739 | 0.277810748 | $1.30\times10^{-2}$ | 3 |
| 33 | *PVRL3−AS1* | 3 | 0.3470860 | 0.045461392 | $1.34\times10^{-2}$ | 1 |
| 34 | *ECD* | 10 | 0.3464802 | 0.201522514 | $1.36\times10^{-2}$ | 3 |
| 35 | *CLK3* | 15 | 0.3446577 | 0.022635883 | $1.42\times10^{-2}$ | 1 |
| 36 | *C4orf22* | 4 | 0.3430783 | 0.034845848 | $1.47\times10^{-2}$ | 1 |
| 37 | *EPN2* | 17 | 0.3405261 | 0.099649864 | $1.56\times10^{-2}$ | 1 |
| 38 | *NR6A1* | 9 | 0.3385413 | 0.412935867 | $1.63\times10^{-2}$ | 4 |
| 39 | *CUX2* | 12 | 0.3370432 | 0.466497502 | $1.68\times10^{-2}$ | 4 |
| 40 | *MIR548O2* | 8 | 0.3367965 | 0.120149341 | $1.69\times10^{-2}$ | 1 |

Table S3: Top 40 SS-H12 sweep candidates at RNA- and protein-coding genes shared between the Western European CEU and South Asian GIH populations. Candidates presented are those that remained after application of a mappability and alignability filter to data (see *Materials and Methods*). Target genes that pass the significance threshold are colored in gold in the "$P$-value" column. Genes whose sweeps are assigned as hard ($\nu = 1$) are shaded in red in the "Inferred $\nu$" column, while soft sweeps ($\nu \geq 2$) are colored in blue.

| | Top gene | Chromosome | Maximum SS–H12 | H2/H1 | *P*–value | Inferred $\nu$ |
|---|---|---|---|---|---|---|
| 1 | *SLC12A1* | 15 | 0.4687775 | 0.01928730 | $< 10^{-6}$ | 1 |
| 2 | *PRMT9* | 4 | 0.3592987 | 0.02846361 | $< 10^{-6}$ | 1 |
| 3 | *ZNF546* | 19 | 0.3543484 | 0.01440592 | $< 10^{-6}$ | 1 |
| 4 | *RNU6−28P* | 15 | 0.3328967 | 0.14154431 | $< 10^{-6}$ | 2 |
| 5 | *PPIP5K1* | 15 | 0.3328967 | 0.14154431 | $< 10^{-6}$ | 2 |
| 6 | *CUX2* | 12 | 0.3244117 | 0.01913100 | $< 10^{-6}$ | 1 |
| 7 | *RUNX1T1* | 8 | 0.3239634 | 0.02660100 | $< 10^{-6}$ | 1 |
| 8 | *HDAC1* | 1 | 0.3000066 | 0.02793417 | $< 10^{-6}$ | 1 |
| 9 | *KIAA0947* | 5 | 0.2985386 | 0.09924729 | $< 10^{-6}$ | 1 |
| 10 | *LINC00478* | 21 | 0.2978392 | 0.03430060 | $< 10^{-6}$ | 1 |
| 11 | *P4HA1* | 10 | 0.2940247 | 0.02457430 | $< 10^{-6}$ | 1 |
| 12 | *BCAS3* | 17 | 0.2928297 | 0.01725947 | $< 10^{-6}$ | 1 |
| 13 | *HS2ST1* | 1 | 0.2925146 | 0.02843537 | $< 10^{-6}$ | 1 |
| 14 | *USP37* | 2 | 0.2841170 | 0.26690328 | $1.0 \times 10^{-6}$ | 2 |
| 15 | *EXOC6B* | 2 | 0.2839310 | 0.03022084 | $1.0 \times 10^{-6}$ | 1 |
| 16 | *TRMT11* | 6 | 0.2823063 | 0.09785459 | $1.0 \times 10^{-6}$ | 1 |
| 17 | *MYO9A* | 15 | 0.2807536 | 0.02811068 | $1.0 \times 10^{-6}$ | 1 |
| 18 | *TFAP2E* | 1 | 0.2788397 | 0.19573284 | $1.0 \times 10^{-6}$ | 2 |
| 19 | *PPM1D* | 17 | 0.2780563 | 0.01891077 | $1.0 \times 10^{-6}$ | 1 |
| 20 | *METTL25* | 12 | 0.2774661 | 0.02117459 | $1.0 \times 10^{-6}$ | 1 |
| 21 | *CCDC178* | 18 | 0.2756744 | 0.08725221 | $1.0 \times 10^{-6}$ | 1 |
| 22 | *USP25* | 21 | 0.2754877 | 0.01694058 | $2.0 \times 10^{-6}$ | 1 |
| 23 | *KIAA0825* | 5 | 0.2741257 | 0.04268183 | $2.0 \times 10^{-6}$ | 1 |
| 24 | *CELSR3* | 3 | 0.2722865 | 0.04291467 | $2.0 \times 10^{-6}$ | 1 |
| 25 | *LOC100188947* | 10 | 0.2716363 | 0.03509440 | $2.0 \times 10^{-6}$ | 1 |
| 26 | *HECTD2* | 10 | 0.2716363 | 0.03509440 | $2.0 \times 10^{-6}$ | 1 |
| 27 | *GNA14* | 9 | 0.2715994 | 0.03925893 | $2.0 \times 10^{-6}$ | 1 |
| 28 | *DNAH6* | 2 | 0.2696911 | 0.02045324 | $3.0 \times 10^{-6}$ | 1 |
| 29 | *FBN1* | 15 | 0.2693902 | 0.04027675 | $4.0 \times 10^{-6}$ | 1 |
| 30 | *LYRM7* | 5 | 0.2668650 | 0.04480839 | $5.0 \times 10^{-6}$ | 1 |
| 31 | *KCNQ5* | 6 | 0.2646294 | 0.01399575 | $6.0 \times 10^{-6}$ | 1 |
| 32 | *OSBPL9* | 1 | 0.2636045 | 0.04380993 | $6.0 \times 10^{-6}$ | 1 |
| 33 | *OTUD6B* | 8 | 0.2635197 | 0.16525761 | $6.0 \times 10^{-6}$ | 2 |
| 34 | *C8orf44−SGK3* | 8 | 0.2629975 | 0.02040620 | $6.0 \times 10^{-6}$ | 1 |
| 35 | *SGK3* | 8 | 0.2629975 | 0.02040620 | $6.0 \times 10^{-6}$ | 1 |
| 36 | *FAM69A* | 1 | 0.2622265 | 0.06324476 | $7.0 \times 10^{-6}$ | 1 |
| 37 | *UNC5D* | 8 | 0.2610994 | 0.03623753 | $7.0 \times 10^{-6}$ | 1 |
| 38 | *KITLG* | 12 | 0.2607259 | 0.05006524 | $8.0 \times 10^{-6}$ | 1 |
| 39 | *PSMB2* | 1 | 0.2596058 | 0.02267442 | $1.0 \times 10^{-5}$ | 1 |
| 40 | *MAP2K5* | 15 | 0.2586188 | 0.33523411 | $1.1 \times 10^{-5}$ | 2 |

Table S4: Top 40 SS-H12 sweep candidates at RNA- and protein-coding genes shared between the Western European CEU and East Asian JPT populations. Candidates presented are those that remained after application of a mappability and alignability filter to data (see *Materials and Methods*). Target genes that pass the significance threshold are colored in gold in the "$P$-value" column. Genes whose sweeps are assigned as hard ($\nu = 1$) are shaded in red in the "Inferred $\nu$" column, while soft sweeps ($\nu \geq 2$) are colored in blue.

| | Top gene | Chromosome | Maximum SS−H12 | H2/H1 | *P*−value | Inferred $\nu$ |
|---|---|---|---|---|---|---|
| 1 | *SPIDR* | 8 | 0.4841559 | 0.13789164 | $< 10^{-6}$ | 2 |
| 2 | *MRAP2* | 6 | 0.3577093 | 0.05691596 | $< 10^{-6}$ | 1 |
| 3 | *BVES−AS1* | 6 | 0.3550710 | 0.35553104 | $< 10^{-6}$ | 2 |
| 4 | *BCAS3* | 17 | 0.3003483 | 0.01124741 | $< 10^{-6}$ | 1 |
| 5 | *RUNX1T1* | 8 | 0.2911443 | 0.01286325 | $1.0{\times}10^{-6}$ | 1 |
| 6 | *DIRC3* | 2 | 0.2846624 | 0.02342707 | $1.0{\times}10^{-6}$ | 1 |
| 7 | *LINC00478* | 21 | 0.2845995 | 0.04674953 | $1.0{\times}10^{-6}$ | 1 |
| 8 | *CASC4* | 15 | 0.2817489 | 0.28631390 | $1.0{\times}10^{-6}$ | 2 |
| 9 | *ZNF546* | 19 | 0.2775094 | 0.01187426 | $2.0{\times}10^{-6}$ | 1 |
| 10 | *C16orf70* | 16 | 0.2759704 | 0.15519459 | $2.0{\times}10^{-6}$ | 2 |
| 11 | *CUX2* | 12 | 0.2711322 | 0.07265968 | $3.0{\times}10^{-6}$ | 1 |
| 12 | *C4orf22* | 4 | 0.2676980 | 0.04613398 | $3.0{\times}10^{-6}$ | 1 |
| 13 | *HDAC1* | 1 | 0.2613163 | 0.01923077 | $4.0{\times}10^{-6}$ | 1 |
| 14 | *EXOC6B* | 2 | 0.2550717 | 0.02907524 | $6.0{\times}10^{-6}$ | 1 |
| 15 | *NCAPG* | 4 | 0.2540412 | 0.47979575 | $6.0{\times}10^{-6}$ | 3 |
| 16 | *DENND1A* | 9 | 0.2502850 | 0.03534972 | $8.0{\times}10^{-6}$ | 1 |
| 17 | *LRRC29* | 16 | 0.2498870 | 0.01559488 | $8.0{\times}10^{-6}$ | 1 |
| 18 | *ZNF780B* | 19 | 0.2453862 | 0.01894595 | $1.2{\times}10^{-5}$ | 1 |
| 19 | *C12orf4* | 12 | 0.2413896 | 0.05908229 | $1.2{\times}10^{-5}$ | 1 |
| 20 | *MRPS11* | 15 | 0.2379901 | 0.04262436 | $1.6{\times}10^{-5}$ | 1 |
| 21 | *USP25* | 21 | 0.2354323 | 0.03030037 | $1.7{\times}10^{-5}$ | 1 |
| 22 | *EPB41L1* | 20 | 0.2351457 | 0.05879280 | $1.7{\times}10^{-5}$ | 1 |
| 23 | *LCORL* | 4 | 0.2345143 | 0.49844508 | $1.7{\times}10^{-5}$ | 3 |
| 24 | *NR6A1* | 9 | 0.2344024 | 0.41874136 | $1.7{\times}10^{-5}$ | 3 |
| 25 | *FOXP2* | 7 | 0.2320563 | 0.06177330 | $1.8{\times}10^{-5}$ | 1 |
| 26 | *PRKDC* | 8 | 0.2316219 | 0.02158066 | $1.8{\times}10^{-5}$ | 1 |
| 27 | *CELSR3* | 3 | 0.2282892 | 0.03061224 | $2.0{\times}10^{-5}$ | 1 |
| 28 | *C2CD5* | 12 | −0.2282773 | 0.40252754 | $2.0{\times}10^{-5}$ | 1 |
| 29 | *LOC100130987* | 11 | 0.2266829 | 0.03103294 | $2.1{\times}10^{-5}$ | 1 |
| 30 | *EPS8* | 12 | 0.2262990 | 0.03675885 | $2.1{\times}10^{-5}$ | 1 |
| 31 | *C5orf42* | 5 | 0.2258946 | 0.23871429 | $2.2{\times}10^{-5}$ | 2 |
| 32 | *SGCD* | 5 | 0.2248583 | 0.04520540 | $2.3{\times}10^{-5}$ | 1 |
| 33 | *PHF20* | 20 | 0.2247397 | 0.03888761 | $2.3{\times}10^{-5}$ | 1 |
| 34 | *KIAA1324L* | 7 | 0.2246612 | 0.05278810 | $2.3{\times}10^{-5}$ | 1 |
| 35 | *SPIN1* | 9 | 0.2244122 | 0.02988283 | $2.5{\times}10^{-5}$ | 1 |
| 36 | *KCND2* | 7 | 0.2238942 | 0.03960568 | $2.6{\times}10^{-5}$ | 1 |
| 37 | *DNAH6* | 2 | 0.2228843 | 0.04768519 | $2.6{\times}10^{-5}$ | 1 |
| 38 | *CCBL2* | 1 | 0.2220159 | 0.38969612 | $2.6{\times}10^{-5}$ | 2 |
| 39 | *DBT* | 1 | 0.2211143 | 0.02032617 | $2.8{\times}10^{-5}$ | 1 |
| 40 | *CLCF1* | 11 | 0.2209926 | 0.03304531 | $2.8{\times}10^{-5}$ | 1 |

Table S5: Top 40 SS-H12 sweep candidates at RNA- and protein-coding genes shared between the Central European CEU and West African YRI populations. Candidates presented are those that remained after application of a mappability and alignability filter to data (see *Materials and Methods*). Target genes that pass the significance threshold are colored in gold in the "$P$-value" column. Genes whose sweeps are assigned as hard ($\nu = 1$) are shaded in red in the "Inferred $\nu$" column, while soft sweeps ($\nu \geq 2$) are colored in blue.

| | Top gene | Chromosome | Maximum SS−H12 | H2/H1 | *P*−value | Inferred $\nu$ |
|---|---|---|---|---|---|---|
| 1 | *RP11−554F20.1* | 9 | 0.4040332 | 0.011758418 | $1.0\times10^{-6}$ | 1 |
| 2 | *SPRED3* | 19 | 0.3495697 | 0.008281479 | $2.0\times10^{-6}$ | 1 |
| 3 | *CASC4* | 15 | 0.3448822 | 0.018257129 | $3.0\times10^{-6}$ | 1 |
| 4 | *KIAA0825* | 5 | 0.3445359 | 0.023951998 | $3.0\times10^{-6}$ | 1 |
| 5 | *ERLIN2* | 8 | 0.3313498 | 0.020023226 | $7.0\times10^{-6}$ | 1 |
| 6 | *DEDD* | 1 | 0.3260732 | 0.060157676 | $8.0\times10^{-6}$ | 1 |
| 7 | *PHKB* | 16 | 0.3236909 | 0.034041921 | $1.0\times10^{-5}$ | 1 |
| 8 | *CPNE1* | 20 | 0.3176717 | 0.029957092 | $1.3\times10^{-5}$ | 1 |
| 9 | *NNT* | 5 | 0.3171650 | 0.017917746 | $1.3\times10^{-5}$ | 1 |
| 10 | *ATP6V1A* | 3 | 0.3132683 | 0.142126360 | $1.7\times10^{-5}$ | 1 |
| 11 | *DDHD2* | 8 | 0.3089642 | 0.058367643 | $2.1\times10^{-5}$ | 1 |
| 12 | *GLRX2* | 1 | 0.3053542 | 0.233674193 | $2.7\times10^{-5}$ | 2 |
| 13 | *DANCR* | 4 | 0.3039967 | 0.040780000 | $2.8\times10^{-5}$ | 1 |
| 14 | *PIGV* | 1 | 0.3036567 | 0.025363186 | $2.9\times10^{-5}$ | 1 |
| 15 | *USP46* | 4 | 0.2995560 | 0.079339450 | $3.3\times10^{-5}$ | 1 |
| 16 | *RALGAPA2* | 20 | 0.2991084 | 0.025897884 | $3.4\times10^{-5}$ | 1 |
| 17 | *GRIK5* | 19 | 0.2981553 | 0.029953502 | $3.5\times10^{-5}$ | 1 |
| 18 | *PLEKHA8* | 7 | 0.2975913 | 0.086839818 | $3.6\times10^{-5}$ | 1 |
| 19 | *MIR548H3* | 6 | 0.2961103 | 0.037196110 | $3.7\times10^{-5}$ | 1 |
| 20 | *PAWR* | 12 | −0.2904662 | 0.440909091 | $4.3\times10^{-5}$ | 1 |
| 21 | *ENTHD1* | 22 | 0.2886265 | 0.085943221 | $4.4\times10^{-5}$ | 1 |
| 22 | *GRIA2* | 4 | 0.2846541 | 0.121846119 | $4.8\times10^{-5}$ | 1 |
| 23 | *DKK2* | 4 | 0.2823235 | 0.022479797 | $5.4\times10^{-5}$ | 1 |
| 24 | *C2orf69* | 2 | 0.2793307 | 0.144744191 | $6.1\times10^{-5}$ | 1 |
| 25 | *ATF2* | 2 | 0.2722884 | 0.046899637 | $7.2\times10^{-5}$ | 1 |
| 26 | *PPAPDC1B* | 8 | 0.2713097 | 0.048928393 | $7.5\times10^{-5}$ | 1 |
| 27 | *CNNM2* | 10 | 0.2699270 | 0.042492595 | $8.0\times10^{-5}$ | 1 |
| 28 | *KIAA1244* | 6 | 0.2590398 | 0.023619457 | $1.21\times10^{-4}$ | 1 |
| 29 | *GABRA4* | 4 | 0.2587261 | 0.254620730 | $1.23\times10^{-4}$ | 3 |
| 30 | *SLC4A10* | 2 | 0.2583449 | 0.033468250 | $1.25\times10^{-4}$ | 1 |
| 31 | *TGFBR1* | 9 | 0.2583162 | 0.097024952 | $1.25\times10^{-4}$ | 1 |
| 32 | *CDK6* | 7 | 0.2580913 | 0.019577085 | $1.26\times10^{-4}$ | 1 |
| 33 | *PHF20* | 20 | 0.2560493 | 0.024958371 | $1.29\times10^{-4}$ | 1 |
| 34 | *SEMA3C* | 7 | 0.2490896 | 0.038386148 | $1.65\times10^{-4}$ | 1 |
| 35 | *SPG11* | 15 | 0.2474120 | 0.038898738 | $1.74\times10^{-4}$ | 1 |
| 36 | *EXOC4* | 7 | 0.2460852 | 0.068343293 | $1.84\times10^{-4}$ | 1 |
| 37 | *ABHD17B* | 9 | 0.2443668 | 0.067033633 | $1.99\times10^{-4}$ | 1 |
| 38 | *SYT1* | 12 | 0.2427578 | 0.071155043 | $2.09\times10^{-4}$ | 1 |
| 39 | *ITFG1* | 16 | 0.2415787 | 0.064650360 | $2.18\times10^{-4}$ | 1 |
| 40 | *WHSC1L1* | 8 | 0.2411594 | 0.091198255 | $2.20\times10^{-4}$ | 1 |

Table S6: Top 40 SS-H12 sweep candidates at RNA- and protein-coding genes shared between the East African LWK and West African YRI populations. Candidates presented are those that remained after application of a mappability and alignability filter to data (see *Materials and Methods*). Target genes that pass the significance threshold are colored in gold in the "$P$-value" column. Genes whose sweeps are assigned as hard ($\nu = 1$) are shaded in red in the "Inferred $\nu$" column, while soft sweeps ($\nu \geq 2$) are colored in blue.

| | Top gene | Chromosome | Maximum SS−H12 | H2/H1 | $P$−value | Inferred $\nu$ |
|---|---|---|---|---|---|---|
| 1 | *GRIK5* | 19 | 0.4257102 | 0.04258992 | $< 10^{-6}$ | 1 |
| 2 | *RGS18* | 1 | 0.3987118 | 0.05121985 | $< 10^{-6}$ | 1 |
| 3 | *RP11−554F20.1* | 9 | 0.3888096 | 0.03007582 | $< 10^{-6}$ | 1 |
| 4 | *SPIDR* | 8 | 0.3877237 | 0.47505841 | $< 10^{-6}$ | 2 |
| 5 | *KIAA0825* | 5 | 0.3792208 | 0.06521739 | $< 10^{-6}$ | 1 |
| 6 | *ARID1A* | 1 | 0.3696765 | 0.05678665 | $< 10^{-6}$ | 1 |
| 7 | *PPAPDC1B* | 8 | 0.3677224 | 0.02213772 | $< 10^{-6}$ | 1 |
| 8 | *MIR548AE2* | 16 | 0.3602984 | 0.01837085 | $< 10^{-6}$ | 1 |
| 9 | *LONP2* | 16 | 0.3602984 | 0.01837085 | $< 10^{-6}$ | 1 |
| 10 | *NNT* | 5 | 0.3578539 | 0.01703393 | $< 10^{-6}$ | 1 |
| 11 | *ATF2* | 2 | 0.3566302 | 0.10607443 | $< 10^{-6}$ | 1 |
| 12 | *CASC4* | 15 | 0.3498801 | 0.02336532 | $< 10^{-6}$ | 1 |
| 13 | *PIGV* | 1 | 0.3469070 | 0.01980346 | $< 10^{-6}$ | 1 |
| 14 | *BRIP1* | 17 | 0.3454437 | 0.05989424 | $< 10^{-6}$ | 1 |
| 15 | *FLJ46284* | 8 | 0.3335889 | 0.34581390 | $< 10^{-6}$ | 2 |
| 16 | *FER1L6* | 8 | 0.3319929 | 0.36210558 | $< 10^{-6}$ | 2 |
| 17 | *FER1L6−AS1* | 8 | 0.3319929 | 0.36210558 | $< 10^{-6}$ | 2 |
| 18 | *PSMD14* | 2 | 0.3302383 | 0.21238349 | $< 10^{-6}$ | 2 |
| 19 | *EHBP1* | 2 | 0.3301406 | 0.04231345 | $< 10^{-6}$ | 1 |
| 20 | *SYT1* | 12 | 0.3268402 | 0.02131281 | $< 10^{-6}$ | 1 |
| 21 | *MAGI3* | 1 | 0.3234881 | 0.03887382 | $< 10^{-6}$ | 1 |
| 22 | *DDHD2* | 8 | 0.3230242 | 0.03319609 | $< 10^{-6}$ | 1 |
| 23 | *TCF4* | 18 | 0.3221087 | 0.04095041 | $< 10^{-6}$ | 1 |
| 24 | *FGFR1* | 8 | 0.3219437 | 0.11190389 | $< 10^{-6}$ | 1 |
| 25 | *DANCR* | 4 | 0.3217441 | 0.02757380 | $< 10^{-6}$ | 1 |
| 26 | *DDX19B* | 16 | 0.3201310 | 0.05961249 | $< 10^{-6}$ | 1 |
| 27 | *PHKB* | 16 | 0.3199744 | 0.14290567 | $< 10^{-6}$ | 2 |
| 28 | *RRN3P3* | 16 | 0.3192156 | 0.12601099 | $< 10^{-6}$ | 2 |
| 29 | *ANK3* | 10 | 0.3168877 | 0.41411632 | $< 10^{-6}$ | 2 |
| 30 | *CNNM2* | 10 | 0.3157693 | 0.04354118 | $< 10^{-6}$ | 1 |
| 31 | *DKK2* | 4 | 0.3148797 | 0.02507450 | $< 10^{-6}$ | 1 |
| 32 | *UCHL5* | 1 | 0.3148279 | 0.23817198 | $< 10^{-6}$ | 2 |
| 33 | *GRIA4* | 11 | 0.3093281 | 0.06513791 | $< 10^{-6}$ | 1 |
| 34 | *HEMGN* | 9 | 0.3087151 | 0.03213826 | $< 10^{-6}$ | 1 |
| 35 | *RNLS* | 10 | 0.3086794 | 0.05394666 | $< 10^{-6}$ | 1 |
| 36 | *TMC1* | 9 | 0.3059124 | 0.05989460 | $< 10^{-6}$ | 1 |
| 37 | *FAM172A* | 5 | 0.3046091 | 0.36377588 | $< 10^{-6}$ | 2 |
| 38 | *NANS* | 9 | 0.3022555 | 0.14421537 | $< 10^{-6}$ | 2 |
| 39 | *SPRED3* | 19 | 0.3006544 | 0.01691324 | $< 10^{-6}$ | 1 |
| 40 | *CCDC178* | 18 | 0.3005286 | 0.06074190 | $< 10^{-6}$ | 1 |

Table S7: Top 40 SS-H12 sweep candidates at RNA- and protein-coding genes shared between the South Asian GIH and West African YRI populations. Candidates presented are those that remained after application of a mappability and alignability filter to data (see *Materials and Methods*). Target genes that pass the significance threshold are colored in gold in the "$P$-value" column. Genes whose sweeps are assigned as hard ($\nu = 1$) are shaded in red in the "Inferred $\nu$" column, while soft sweeps ($\nu \geq 2$) are colored in blue.

| | Top gene | Chromosome | Maximum SS−H12 | H2/H1 | *P*−value | Inferred ν |
|---|---|---|---|---|---|---|
| 1 | *RP11−554F20.1* | 9 | 0.3771248 | 0.011770772 | $< 10^{-6}$ | 1 |
| 2 | *ATP6V1A* | 3 | 0.3639640 | 0.099682934 | $< 10^{-6}$ | 1 |
| 3 | *DDHD2* | 8 | 0.3602952 | 0.013720551 | $< 10^{-6}$ | 1 |
| 4 | *CDK6* | 7 | 0.3578316 | 0.071008263 | $< 10^{-6}$ | 1 |
| 5 | *KIAA0825* | 5 | 0.3525741 | 0.027694560 | $< 10^{-6}$ | 1 |
| 6 | *SYT1* | 12 | 0.3492609 | 0.012266522 | $< 10^{-6}$ | 1 |
| 7 | *PSMD14* | 2 | 0.3412286 | 0.135670252 | $< 10^{-6}$ | 1 |
| 8 | *WHSC1L1* | 8 | 0.3283306 | 0.043503219 | $< 10^{-6}$ | 1 |
| 9 | *PHKB* | 16 | 0.3232317 | 0.042223410 | $< 10^{-6}$ | 1 |
| 10 | *NNT* | 5 | 0.3223767 | 0.017984343 | $< 10^{-6}$ | 1 |
| 11 | *PPAPDC1B* | 8 | 0.3196440 | 0.032132443 | $< 10^{-6}$ | 1 |
| 12 | *FBXW4* | 10 | 0.3196397 | 0.031138790 | $< 10^{-6}$ | 1 |
| 13 | *CASC4* | 15 | 0.3150844 | 0.009182066 | $< 10^{-6}$ | 1 |
| 14 | *PLEKHA8* | 7 | 0.3142622 | 0.083226858 | $< 10^{-6}$ | 1 |
| 15 | *PIGV* | 1 | 0.3115851 | 0.031716870 | $< 10^{-6}$ | 1 |
| 16 | *ZNF451* | 6 | 0.3085450 | 0.029623301 | $< 10^{-6}$ | 1 |
| 17 | *MIR548AE2* | 16 | 0.3065668 | 0.025105816 | $< 10^{-6}$ | 1 |
| 18 | *LONP2* | 16 | 0.3065668 | 0.025105816 | $< 10^{-6}$ | 1 |
| 19 | *SLC4A10* | 2 | 0.3062220 | 0.024992628 | $< 10^{-6}$ | 1 |
| 20 | *DEDD* | 1 | 0.2978007 | 0.043610857 | $< 10^{-6}$ | 1 |
| 21 | *USP46* | 4 | 0.2930645 | 0.043401898 | $< 10^{-6}$ | 1 |
| 22 | *ATF2* | 2 | 0.2890980 | 0.042325935 | $< 10^{-6}$ | 1 |
| 23 | *C10orf76* | 10 | 0.2791604 | 0.241890959 | $< 10^{-6}$ | 2 |
| 24 | *GRIK5* | 19 | 0.2789334 | 0.026200519 | $< 10^{-6}$ | 1 |
| 25 | *FRYL* | 4 | 0.2770178 | 0.124549768 | $< 10^{-6}$ | 1 |
| 26 | *ABHD17B* | 9 | 0.2761729 | 0.019974819 | $1.0{\times}10^{-6}$ | 1 |
| 27 | *CPNE1* | 20 | 0.2749730 | 0.053890867 | $1.0{\times}10^{-6}$ | 1 |
| 28 | *RALGAPA2* | 20 | 0.2742919 | 0.031716870 | $1.0{\times}10^{-6}$ | 1 |
| 29 | *COL7A1* | 3 | 0.2710832 | 0.030527862 | $1.0{\times}10^{-6}$ | 1 |
| 30 | *PAWR* | 12 | −0.2695280 | 0.500033047 | $2.0{\times}10^{-6}$ | 1 |
| 31 | *ERLIN2* | 8 | 0.2694367 | 0.032420110 | $2.0{\times}10^{-6}$ | 1 |
| 32 | *TYW5* | 2 | 0.2662606 | 0.079070667 | $2.0{\times}10^{-6}$ | 1 |
| 33 | *KCNIP2* | 10 | 0.2654300 | 0.188650307 | $2.0{\times}10^{-6}$ | 2 |
| 34 | *ENTHD1* | 22 | 0.2648954 | 0.059516605 | $2.0{\times}10^{-6}$ | 1 |
| 35 | *CADM1* | 11 | 0.2608986 | 0.130764109 | $2.0{\times}10^{-6}$ | 1 |
| 36 | *NIT1* | 1 | 0.2589696 | 0.060503570 | $2.0{\times}10^{-6}$ | 1 |
| 37 | *GRIA4* | 11 | 0.2580788 | 0.177060197 | $2.0{\times}10^{-6}$ | 2 |
| 38 | *CAMK2G* | 10 | 0.2552114 | 0.145238523 | $3.0{\times}10^{-6}$ | 1 |
| 39 | *DANCR* | 4 | 0.2530703 | 0.034185402 | $3.0{\times}10^{-6}$ | 1 |
| 40 | *PLBD2* | 12 | 0.2530187 | 0.025798687 | $3.0{\times}10^{-6}$ | 1 |

Table S8: Top 40 SS-H12 sweep candidates at RNA- and protein-coding genes shared between the East Asian JPT and West African YRI populations. Candidates presented are those that remained after application of a mappability and alignability filter to data (see *Materials and Methods*). Target genes that pass the significance threshold are colored in gold in the "$P$-value" column. Genes whose sweeps are assigned as hard ($\nu = 1$) are shaded in red in the "Inferred $\nu$" column, while soft sweeps ($\nu \geq 2$) are colored in blue.

| | Top gene | Chromosome | Maximum SS−H12 | H2/H1 | *P*–value | Inferred $\nu$ |
|---|---|---|---|---|---|---|
| 1 | *RP11−554F20.1* | 9 | 0.3931760 | 0.106986532 | $< 10^{-6}$ | 1 |
| 2 | *HEMGN* | 9 | 0.3775289 | 0.024845674 | $< 10^{-6}$ | 1 |
| 3 | *ATP6V1A* | 3 | 0.3612693 | 0.187013598 | $< 10^{-6}$ | 3 |
| 4 | *PSMD14* | 2 | 0.3478155 | 0.120552739 | $< 10^{-6}$ | 1 |
| 5 | *KIAA0825* | 5 | 0.3462541 | 0.042399755 | $< 10^{-6}$ | 1 |
| 6 | *ENTHD1* | 22 | 0.3457729 | 0.024417586 | $< 10^{-6}$ | 1 |
| 7 | *UCHL5* | 1 | 0.3410178 | 0.296451830 | $< 10^{-6}$ | 2 |
| 8 | *EHBP1* | 2 | 0.3296022 | 0.040707208 | $< 10^{-6}$ | 1 |
| 9 | *DDHD2* | 8 | 0.3232796 | 0.047658115 | $< 10^{-6}$ | 1 |
| 10 | *GRIK5* | 19 | 0.3201569 | 0.022160505 | $< 10^{-6}$ | 1 |
| 11 | *CPNE1* | 20 | 0.3095707 | 0.020447795 | $< 10^{-6}$ | 1 |
| 12 | *CASC4* | 15 | 0.3082424 | 0.008452639 | $< 10^{-6}$ | 1 |
| 13 | *PIGV* | 1 | 0.3080457 | 0.019652483 | $< 10^{-6}$ | 1 |
| 14 | *GRIA4* | 11 | 0.2877642 | 0.076983380 | $< 10^{-6}$ | 1 |
| 15 | *FBXW4* | 10 | 0.2812126 | 0.032817676 | $< 10^{-6}$ | 1 |
| 16 | *USP46* | 4 | 0.2778346 | 0.050033190 | $< 10^{-6}$ | 1 |
| 17 | *PPAPDC1B* | 8 | 0.2770218 | 0.067697788 | $< 10^{-6}$ | 1 |
| 18 | *CSMD3* | 8 | 0.2767410 | 0.025265593 | $< 10^{-6}$ | 1 |
| 19 | *AUTS2* | 7 | 0.2739576 | 0.048349205 | $< 10^{-6}$ | 1 |
| 20 | *WDPCP* | 2 | 0.2732119 | 0.160070411 | $< 10^{-6}$ | 2 |
| 21 | *NNT* | 5 | 0.2719657 | 0.011428056 | $< 10^{-6}$ | 1 |
| 22 | *WDR75* | 2 | 0.2713068 | 0.477669319 | $< 10^{-6}$ | 3 |
| 23 | *MAST2* | 1 | 0.2710749 | 0.113317105 | $< 10^{-6}$ | 1 |
| 24 | *MPHOSPH9* | 12 | −0.2671972 | 0.500226740 | $1.0{\times}10^{-6}$ | 1 |
| 25 | *ATF2* | 2 | 0.2661096 | 0.050953776 | $1.0{\times}10^{-6}$ | 1 |
| 26 | *MIR548H3* | 6 | 0.2657886 | 0.044716955 | $1.0{\times}10^{-6}$ | 1 |
| 27 | *DKK2* | 4 | 0.2654977 | 0.020832637 | $1.0{\times}10^{-6}$ | 1 |
| 28 | *RALGAPA2* | 20 | 0.2634601 | 0.018345166 | $1.0{\times}10^{-6}$ | 1 |
| 29 | *PLEKHA8* | 7 | 0.2629720 | 0.065646654 | $1.0{\times}10^{-6}$ | 1 |
| 30 | *ABCA17P* | 16 | 0.2627850 | 0.029987576 | $1.0{\times}10^{-6}$ | 1 |
| 31 | *ERLIN2* | 8 | 0.2612417 | 0.051206414 | $1.0{\times}10^{-6}$ | 1 |
| 32 | *LOC644554* | 19 | 0.2599085 | 0.046211073 | $1.0{\times}10^{-6}$ | 1 |
| 33 | *EXOC6B* | 2 | −0.2591453 | 0.400289321 | $1.0{\times}10^{-6}$ | 1 |
| 34 | *ARID1A* | 1 | 0.2586148 | 0.012315850 | $1.0{\times}10^{-6}$ | 1 |
| 35 | *AC016582.2* | 19 | 0.2575930 | 0.184115407 | $1.0{\times}10^{-6}$ | 2 |
| 36 | *USP31* | 16 | 0.2542219 | 0.048774295 | $1.0{\times}10^{-6}$ | 1 |
| 37 | *WDR87* | 19 | 0.2539212 | 0.052847194 | $1.0{\times}10^{-6}$ | 1 |
| 38 | *NCK1* | 3 | 0.2533815 | 0.105668649 | $1.0{\times}10^{-6}$ | 1 |
| 39 | *MIR548AE2* | 16 | 0.2530337 | 0.012070399 | $1.0{\times}10^{-6}$ | 1 |
| 40 | *LONP2* | 16 | 0.2530337 | 0.012070399 | $1.0{\times}10^{-6}$ | 1 |

Table S9: Top 40 SS-H12 sweep candidates at RNA- and protein-coding genes shared between the East Asian JPT and South Asian GIH populations. Candidates presented are those that remained after application of a mappability and alignability filter to data (see *Materials and Methods*). Target genes that pass the significance threshold are colored in gold in the "$P$-value" column. Genes whose sweeps are assigned as hard ($\nu = 1$) are shaded in red in the "Inferred $\nu$" column, while soft sweeps ($\nu \geq 2$) are colored in blue.

| | Top gene | Chromosome | Maximum SS−H12 | H2/H1 | *P*−value | Inferred $\nu$ |
|---|---|---|---|---|---|---|
| 1 | *RUNX1T1* | 8 | 0.4474761 | 0.013112088 | $< 10^{-6}$ | 1 |
| 2 | *P4HTM* | 3 | 0.4140126 | 0.004079071 | $< 10^{-6}$ | 1 |
| 3 | *TMTC2* | 12 | 0.3420391 | 0.025676243 | $< 10^{-6}$ | 1 |
| 4 | *EXOC6B* | 2 | 0.3326926 | 0.019515562 | $< 10^{-6}$ | 1 |
| 5 | *RNF121* | 11 | 0.3155150 | 0.030228890 | $< 10^{-6}$ | 1 |
| 6 | *BCAS3* | 17 | 0.3120734 | 0.020570739 | $< 10^{-6}$ | 1 |
| 7 | *HELZ* | 17 | 0.3098360 | 0.111157784 | $< 10^{-6}$ | 1 |
| 8 | *ADAMTS6* | 5 | 0.3076773 | 0.407162839 | $< 10^{-6}$ | 3 |
| 9 | *ORC2* | 2 | 0.3055687 | 0.018545289 | $< 10^{-6}$ | 1 |
| 10 | *CUX2* | 12 | 0.3012879 | 0.114717964 | $< 10^{-6}$ | 1 |
| 11 | *APPBP2* | 17 | 0.2943597 | 0.018755226 | $< 10^{-6}$ | 1 |
| 12 | *ZNF546* | 19 | 0.2903935 | 0.026486969 | $< 10^{-6}$ | 1 |
| 13 | *RP11−682N22.1* | 7 | 0.2866539 | 0.025520555 | $< 10^{-6}$ | 1 |
| 14 | *LINC00478* | 21 | 0.2768036 | 0.051422319 | $< 10^{-6}$ | 1 |
| 15 | *DBT* | 1 | 0.2754919 | 0.019618240 | $< 10^{-6}$ | 1 |
| 16 | *SLC44A1* | 9 | 0.2751889 | 0.027447371 | $< 10^{-6}$ | 1 |
| 17 | *DPH5* | 1 | 0.2734798 | 0.063964153 | $< 10^{-6}$ | 1 |
| 18 | *MBOAT2* | 2 | 0.2732935 | 0.022872389 | $< 10^{-6}$ | 1 |
| 19 | *DR1* | 1 | 0.2719058 | 0.117944872 | $< 10^{-6}$ | 1 |
| 20 | *ZNF780B* | 19 | 0.2710979 | 0.027775374 | $< 10^{-6}$ | 1 |
| 21 | *PRPF40B* | 12 | 0.2704860 | 0.009056569 | $< 10^{-6}$ | 1 |
| 22 | *LOC100188947* | 10 | 0.2693331 | 0.091476681 | $< 10^{-6}$ | 1 |
| 23 | *HECTD2* | 10 | 0.2693331 | 0.091476681 | $< 10^{-6}$ | 1 |
| 24 | *USP32* | 17 | 0.2668435 | 0.030825697 | $< 10^{-6}$ | 1 |
| 25 | *NF1* | 17 | 0.2650127 | 0.446732585 | $< 10^{-6}$ | 2 |
| 26 | *RP11−53O19.1* | 5 | 0.2645393 | 0.266671012 | $< 10^{-6}$ | 2 |
| 27 | *HDAC1* | 1 | 0.2631774 | 0.038364271 | $< 10^{-6}$ | 1 |
| 28 | *ABCB1* | 7 | 0.2624699 | 0.032952008 | $< 10^{-6}$ | 1 |
| 29 | *RUNDC3B* | 7 | 0.2624699 | 0.032952008 | $< 10^{-6}$ | 1 |
| 30 | *ADK* | 10 | 0.2592457 | 0.395106884 | $< 10^{-6}$ | 2 |
| 31 | *CNNM4* | 2 | 0.2583070 | 0.084782739 | $< 10^{-6}$ | 1 |
| 32 | *DDB1* | 11 | 0.2571627 | 0.012680739 | $< 10^{-6}$ | 1 |
| 33 | *XXYLT1* | 3 | 0.2563662 | 0.022777439 | $< 10^{-6}$ | 1 |
| 34 | *COMMD3−BMI1* | 10 | 0.2549803 | 0.029266377 | $< 10^{-6}$ | 1 |
| 35 | *BMI1* | 10 | 0.2549803 | 0.029266377 | $< 10^{-6}$ | 1 |
| 36 | *PRKAR2A* | 3 | 0.2543194 | 0.029957868 | $< 10^{-6}$ | 1 |
| 37 | *USP25* | 21 | 0.2500539 | 0.039729584 | $< 10^{-6}$ | 1 |
| 38 | *ATP6V0D1* | 16 | 0.2497646 | 0.015632539 | $< 10^{-6}$ | 1 |
| 39 | *GPC5* | 13 | 0.2484834 | 0.088347003 | $< 10^{-6}$ | 1 |
| 40 | *SPATA31D3* | 9 | 0.2473412 | 0.031572111 | $< 10^{-6}$ | 1 |

Table S10: Top 40 SS-H12 sweep candidates at RNA- and protein-coding genes shared between the East Asian JPT and Southeast Asian KHV populations. Candidates presented are those that remained after application of a mappability and alignability filter to data (see *Materials and Methods*). Target genes that pass the significance threshold are colored in gold in the "$P$-value" column. Genes whose sweeps are assigned as hard ($\nu = 1$) are shaded in red in the "Inferred $\nu$" column, while soft sweeps ($\nu \geq 2$) are colored in blue.

| | Top gene | Chromosome | Maximum SS−H12 | H2/H1 | *P*−value | Inferred $\nu$ |
|---|---|---|---|---|---|---|
| *1* | *EXOC6B* | 2 | 0.5264721 | 0.006772899 | $< 10^{-6}$ | 1 |
| *2* | *SPAG6* | 10 | 0.4968819 | 0.006607293 | $< 10^{-6}$ | 1 |
| *3* | *P4HTM* | 3 | 0.4842239 | 0.003800968 | $< 10^{-6}$ | 1 |
| *4* | *RUNX1T1* | 8 | 0.4762151 | 0.012305665 | $< 10^{-6}$ | 1 |
| *5* | *C11orf49* | 11 | 0.4727695 | 0.068502981 | $< 10^{-6}$ | 1 |
| *6* | *RP11−696N14.1* | 4 | 0.4622633 | 0.047157537 | $< 10^{-6}$ | 1 |
| *7* | *EXD2* | 14 | 0.4488191 | 0.012119385 | $< 10^{-6}$ | 1 |
| *8* | *BCL2L1* | 20 | 0.4334603 | 0.011505081 | $1.0\times10^{-6}$ | 1 |
| *9* | *SPIDR* | 8 | 0.4323823 | 0.121946971 | $1.0\times10^{-6}$ | 2 |
| *10* | *TMEM33* | 4 | 0.4163834 | 0.013441981 | $1.0\times10^{-6}$ | 1 |
| *11* | *HMCN1* | 1 | 0.4139426 | 0.433746297 | $1.0\times10^{-6}$ | 2 |
| *12* | *TRMT11* | 6 | 0.4132626 | 0.008565410 | $1.0\times10^{-6}$ | 1 |
| *13* | *CSPP1* | 8 | 0.4131015 | 0.370451157 | $1.0\times10^{-6}$ | 2 |
| *14* | *BCL7C* | 16 | 0.4116698 | 0.017374975 | $1.0\times10^{-6}$ | 1 |
| *15* | *AMBRA1* | 11 | 0.4078245 | 0.026064925 | $3.0\times10^{-6}$ | 1 |
| *16* | *ADH1A* | 4 | 0.4068527 | 0.054411554 | $3.0\times10^{-6}$ | 1 |
| *17* | *LINC00536* | 8 | 0.4047586 | 0.319893205 | $4.0\times10^{-6}$ | 2 |
| *18* | *TRUB1* | 10 | 0.3960872 | 0.012888666 | $8.0\times10^{-6}$ | 1 |
| *19* | *SLC25A20* | 3 | 0.3951800 | 0.306872504 | $1.0\times10^{-5}$ | 2 |
| *20* | *NOVA1* | 14 | 0.3937128 | 0.296371985 | $1.1\times10^{-5}$ | 2 |
| *21* | *UBE3A* | 15 | 0.3881960 | 0.421256404 | $1.3\times10^{-5}$ | 2 |
| *22* | *RAD51B* | 14 | 0.3877928 | 0.020655194 | $1.3\times10^{-5}$ | 1 |
| *23* | *C2orf66* | 2 | 0.3847481 | 0.011183304 | $1.4\times10^{-5}$ | 1 |
| *24* | *ARIH2* | 3 | 0.3835818 | 0.011305712 | $1.5\times10^{-5}$ | 1 |
| *25* | *FBXO4* | 5 | 0.3830810 | 0.060543858 | $1.5\times10^{-5}$ | 1 |
| *26* | *PURA* | 5 | 0.3821055 | 0.043743931 | $1.5\times10^{-5}$ | 1 |
| *27* | *LOC100188947* | 10 | 0.3819967 | 0.126665136 | $1.5\times10^{-5}$ | 2 |
| *28* | *HECTD2* | 10 | 0.3819967 | 0.126665136 | $1.5\times10^{-5}$ | 2 |
| *29* | *ZNF282* | 7 | 0.3789665 | 0.013728048 | $1.5\times10^{-5}$ | 1 |
| *30* | *SPATS2* | 12 | 0.3779965 | 0.018103713 | $1.7\times10^{-5}$ | 1 |
| *31* | *DPH6* | 15 | 0.3765334 | 0.016983232 | $1.7\times10^{-5}$ | 1 |
| *32* | *ARIH2OS* | 3 | 0.3750646 | 0.011348289 | $1.9\times10^{-5}$ | 1 |
| *33* | *KCNT2* | 1 | 0.3748550 | 0.093205290 | $1.9\times10^{-5}$ | 1 |
| *34* | *GPHN* | 14 | 0.3744098 | 0.478385232 | $1.9\times10^{-5}$ | 2 |
| *35* | *PHYHIPL* | 10 | 0.3737335 | 0.020357412 | $2.2\times10^{-5}$ | 1 |
| *36* | *CCDC18* | 1 | 0.3732415 | 0.124636275 | $2.2\times10^{-5}$ | 2 |
| *37* | *ABHD17B* | 9 | 0.3730153 | 0.059902525 | $2.2\times10^{-5}$ | 1 |
| *38* | *IFT81* | 12 | 0.3714945 | 0.220865456 | $2.2\times10^{-5}$ | 2 |
| *39* | *FHOD1* | 16 | 0.3711776 | 0.013442943 | $2.2\times10^{-5}$ | 1 |
| *40* | *SYNJ1* | 21 | 0.3678745 | 0.211024411 | $2.5\times10^{-5}$ | 2 |

Table S11: Top 40 SS-G123 sweep candidates at RNA- and protein-coding genes shared between the Central and Western European CEU and GBR populations. Candidates presented are those that remained after application of a mappability and alignability filter to data (see *Materials and Methods*). Target genes that pass the significance threshold are colored in gold in the "$P$-value" column. Genes whose sweeps are assigned as hard ($\nu = 1$) are shaded in red in the "Inferred $\nu$" column, while soft sweeps ($\nu \geq 2$) are colored in blue.

| | Top gene | Chromosome | Maximum SS−G123 | G2/G1 | *P*–value | Inferred $\nu$ |
|---|---|---|---|---|---|---|
| 1 | *ZRANB3* | 2 | 0.4036658 | 0.01992548 | $2.0\times10^{-6}$ | 1 |
| 2 | *R3HDM1* | 2 | 0.4001688 | 0.03074343 | $3.0\times10^{-6}$ | 1 |
| 3 | *DARS* | 2 | 0.3939288 | 0.04113419 | $4.0\times10^{-6}$ | 1 |
| 4 | *SLC12A1* | 15 | 0.3494603 | 0.03411498 | $1.2\times10^{-5}$ | 1 |
| 5 | *MCM6* | 2 | 0.3425388 | 0.04473161 | $1.8\times10^{-5}$ | 1 |
| 6 | *AC093391.2* | 2 | 0.2944935 | 0.04039653 | $8.0\times10^{-5}$ | 1 |
| 7 | *LCT* | 2 | 0.2864912 | 0.02114967 | $1.00\times10^{-4}$ | 1 |
| 8 | *UBXN4* | 2 | 0.2740023 | 0.02600907 | $1.47\times10^{-4}$ | 1 |
| 9 | *LOC100507600* | 2 | 0.2706452 | 0.02798382 | $1.68\times10^{-4}$ | 1 |
| 10 | *RAB3GAP1* | 2 | 0.2506534 | 0.06432749 | $3.48\times10^{-4}$ | 1 |
| 11 | *KMT2A* | 11 | 0.2041324 | 0.10449343 | $1.73\times10^{-3}$ | 1 |
| 12 | *BCAS3* | 17 | 0.1859900 | 0.09758002 | $3.05\times10^{-3}$ | 1 |
| 13 | *PPM1D* | 17 | 0.1800870 | 0.07265290 | $3.63\times10^{-3}$ | 1 |
| 14 | *UNC5D* | 8 | 0.1780280 | 0.39270231 | $3.86\times10^{-3}$ | 3 |
| 15 | *KAT6B* | 10 | 0.1707262 | 0.05624037 | $4.80\times10^{-3}$ | 1 |
| 16 | *TMEM116* | 12 | 0.1651175 | 0.18743961 | $5.66\times10^{-3}$ | 1 |
| 17 | *PRKDC* | 8 | 0.1628838 | 0.14634787 | $6.00\times10^{-3}$ | 1 |
| 18 | *MYO9A* | 15 | 0.1598302 | 0.06094675 | $6.56\times10^{-3}$ | 1 |
| 19 | *ACMSD* | 2 | 0.1572425 | 0.15455562 | $7.09\times10^{-3}$ | 1 |
| 20 | *CCNT2−AS1* | 2 | 0.1572425 | 0.15455562 | $7.09\times10^{-3}$ | 1 |
| 21 | *ZNF546* | 19 | 0.1563353 | 0.28800989 | $7.27\times10^{-3}$ | 2 |
| 22 | *C4orf22* | 4 | 0.1548723 | 0.17300057 | $7.57\times10^{-3}$ | 1 |
| 23 | *COL5A2* | 2 | 0.1531146 | 0.07664563 | $7.97\times10^{-3}$ | 1 |
| 24 | *KITLG* | 12 | 0.1507429 | 0.09976581 | $8.55\times10^{-3}$ | 1 |
| 25 | *MRPS31* | 13 | 0.1404181 | 0.66863672 | $1.17\times10^{-2}$ | 6 |
| 26 | *LAMA3* | 18 | 0.1397230 | 0.41137771 | $1.19\times10^{-2}$ | 3 |
| 27 | *AGO3* | 1 | 0.1389525 | 0.18726491 | $1.22\times10^{-2}$ | 1 |
| 28 | *RALGAPA1* | 14 | 0.1372367 | 0.10397132 | $1.28\times10^{-2}$ | 1 |
| 29 | *RALGAPA1P* | 14 | 0.1372367 | 0.10397132 | $1.28\times10^{-2}$ | 1 |
| 30 | *SYT1* | 12 | 0.1368878 | 0.18617021 | $1.29\times10^{-2}$ | 1 |
| 31 | *GFRA2* | 8 | 0.1354658 | 0.51413190 | $1.35\times10^{-2}$ | 4 |
| 32 | *CNBD2* | 20 | 0.1345655 | 0.10348771 | $1.39\times10^{-2}$ | 1 |
| 33 | *PIK3R4* | 3 | 0.1315715 | 0.31331699 | $1.51\times10^{-2}$ | 2 |
| 34 | *CDK6* | 7 | 0.1302599 | 0.19966667 | $1.57\times10^{-2}$ | 1 |
| 35 | *DIRC3* | 2 | 0.1301000 | 0.13940724 | $1.58\times10^{-2}$ | 1 |
| 36 | *GNA14* | 9 | 0.1297811 | 0.11111111 | $1.59\times10^{-2}$ | 1 |
| 37 | *EXOC5* | 14 | 0.1292055 | 0.27603759 | $1.62\times10^{-2}$ | 1 |
| 38 | *SPATA5L1* | 15 | 0.1291950 | 0.50091659 | $1.62\times10^{-2}$ | 4 |
| 39 | *AC005592.1* | 5 | 0.1283393 | 0.05882353 | $1.66\times10^{-2}$ | 1 |
| 40 | *TMEM163* | 2 | 0.1271473 | 0.27047556 | $1.72\times10^{-2}$ | 1 |

Table S12: Top 40 SS-G123 sweep candidates at RNA- and protein-coding genes shared between the Western European CEU and South Asian GIH populations. Candidates presented are those that remained after application of a mappability and alignability filter to data (see *Materials and Methods*). Target genes that pass the significance threshold are colored in gold in the "$P$-value" column. Genes whose sweeps are assigned as hard ($\nu = 1$) are shaded in red in the "Inferred $\nu$" column, while soft sweeps ($\nu \geq 2$) are colored in blue.

| | Top gene | Chromosome | Maximum SS−G123 | G2/G1 | *P*−value | Inferred $\nu$ |
|---|---|---|---|---|---|---|
| *1* | *SLC12A1* | 15 | 0.24849898 | 0.07039983 | $< 10^{-6}$ | 1 |
| *2* | *RUNX1T1* | 8 | 0.16146633 | 0.14275121 | $< 10^{-6}$ | 1 |
| *3* | *ZNF546* | 19 | 0.13139163 | 0.31590106 | $< 10^{-6}$ | 2 |
| *4* | *RNU6−28P* | 15 | 0.12412153 | 0.49297753 | $< 10^{-6}$ | 2 |
| *5* | *PPIP5K1* | 15 | 0.12412153 | 0.49297753 | $< 10^{-6}$ | 2 |
| *6* | *PRMT9* | 4 | 0.12362840 | 0.12395833 | $< 10^{-6}$ | 1 |
| *7* | *BCAS3* | 17 | 0.12155652 | 0.05834652 | $< 10^{-6}$ | 1 |
| *8* | *HDAC1* | 1 | 0.11263346 | 0.10144553 | $< 10^{-6}$ | 1 |
| *9* | *P4HA1* | 10 | 0.10999698 | 0.16650149 | $1.0\times10^{-6}$ | 1 |
| *10* | *PITPNB* | 22 | 0.10563578 | 0.35822785 | $1.0\times10^{-6}$ | 2 |
| *11* | *CUX2* | 12 | 0.10209057 | 0.06924939 | $2.0\times10^{-6}$ | 1 |
| *12* | *NECAB1* | 8 | 0.10197638 | 0.25569620 | $2.0\times10^{-6}$ | 1 |
| *13* | *HS2ST1* | 1 | 0.10041528 | 0.10199005 | $2.0\times10^{-6}$ | 1 |
| *14* | *METTL25* | 12 | 0.09952687 | 0.08587185 | $2.0\times10^{-6}$ | 1 |
| *15* | *CELSR3* | 3 | 0.09761571 | 0.12500000 | $2.0\times10^{-6}$ | 1 |
| *16* | *KCNQ5* | 6 | 0.09516938 | 0.08284819 | $2.0\times10^{-6}$ | 1 |
| *17* | *USP25* | 21 | 0.09485193 | 0.11653037 | $2.0\times10^{-6}$ | 1 |
| *18* | *OSBPL9* | 1 | 0.09477321 | 0.16461204 | $2.0\times10^{-6}$ | 1 |
| *19* | *PPM1D* | 17 | 0.09457751 | 0.08187773 | $2.0\times10^{-6}$ | 1 |
| *20* | *PSMB2* | 1 | 0.09368669 | 0.10872127 | $2.0\times10^{-6}$ | 1 |
| *21* | *KITLG* | 12 | 0.09321913 | 0.10938491 | $2.0\times10^{-6}$ | 1 |
| *22* | *PTPRK* | 6 | 0.09238986 | 0.15235110 | $2.0\times10^{-6}$ | 1 |
| *23* | *PRPF40B* | 12 | 0.08811411 | 0.10312899 | $2.0\times10^{-6}$ | 1 |
| *24* | *EXOC5* | 14 | 0.08680119 | 0.14383562 | $2.0\times10^{-6}$ | 1 |
| *25* | *NF1* | 17 | 0.08654684 | 0.53987730 | $2.0\times10^{-6}$ | 2 |
| *26* | *KIAA0825* | 5 | 0.08651185 | 0.13956568 | $2.0\times10^{-6}$ | 1 |
| *27* | *KLHL28* | 14 | 0.08623216 | 0.08287895 | $3.0\times10^{-6}$ | 1 |
| *28* | *UNC5D* | 8 | 0.08584013 | 0.23278008 | $4.0\times10^{-6}$ | 1 |
| *29* | *C8orf44−SGK3* | 8 | 0.08500952 | 0.10876730 | $4.0\times10^{-6}$ | 1 |
| *30* | *SGK3* | 8 | 0.08500952 | 0.10876730 | $4.0\times10^{-6}$ | 1 |
| *31* | *GNA14* | 9 | 0.08464131 | 0.20441989 | $4.0\times10^{-6}$ | 1 |
| *32* | *POLN* | 4 | 0.08228096 | 0.25619469 | $4.0\times10^{-6}$ | 1 |
| *33* | *FMNL3* | 12 | 0.08227142 | 0.09967949 | $4.0\times10^{-6}$ | 1 |
| *34* | *IFT80* | 3 | 0.08224157 | 0.65133172 | $4.0\times10^{-6}$ | 4 |
| *35* | *TFAP2E* | 1 | 0.08005606 | 0.45766932 | $6.0\times10^{-6}$ | 2 |
| *36* | *KIAA0947* | 5 | 0.08003881 | 0.33080329 | $6.0\times10^{-6}$ | 1 |
| *37* | *LINC00478* | 21 | 0.07852482 | 0.16499666 | $6.0\times10^{-6}$ | 1 |
| *38* | *EXOC6B* | 2 | 0.07730652 | 0.14328063 | $6.0\times10^{-6}$ | 1 |
| *39* | *FAM149B1* | 10 | 0.07630036 | 0.14383562 | $6.0\times10^{-6}$ | 1 |
| *40* | *CCDC178* | 18 | 0.07628428 | 0.33479106 | $6.0\times10^{-6}$ | 1 |

Table S13: Top 40 SS-G123 sweep candidates at RNA- and protein-coding genes shared between the Western European CEU and East Asian JPT populations. Candidates presented are those that remained after application of a mappability and alignability filter to data (see *Materials and Methods*). Target genes that pass the significance threshold are colored in gold in the "$P$-value" column. Genes whose sweeps are assigned as hard ($\nu = 1$) are shaded in red in the "Inferred $\nu$" column, while soft sweeps ($\nu \geq 2$) are colored in blue.

| | Top gene | Chromosome | Maximum SS−G123 | G2/G1 | $P$−value | Inferred $\nu$ |
|---|---|---|---|---|---|---|
| 1 | *SPIDR* | 8 | 0.19359701 | 0.31787351 | $< 10^{-6}$ | 3 |
| 2 | *RUNX1T1* | 8 | 0.11275519 | 0.05393330 | $< 10^{-6}$ | 1 |
| 3 | *MRAP2* | 6 | 0.11102258 | 0.25897350 | $< 10^{-6}$ | 1 |
| 4 | *BVES−AS1* | 6 | 0.10952935 | 0.51920530 | $< 10^{-6}$ | 2 |
| 5 | *LINC00478* | 21 | 0.10122097 | 0.27387331 | $< 10^{-6}$ | 1 |
| 6 | *USP25* | 21 | 0.09815414 | 0.14385858 | $< 10^{-6}$ | 1 |
| 7 | *BCAS3* | 17 | 0.09127146 | 0.06342583 | $< 10^{-6}$ | 1 |
| 8 | *DNAH6* | 2 | 0.08296677 | 0.20867526 | $< 10^{-6}$ | 1 |
| 9 | *BEND4* | 4 | 0.08124753 | 0.09624478 | $< 10^{-6}$ | 1 |
| 10 | *ZNF546* | 19 | 0.07817327 | 0.06845577 | $2.0{\times}10^{-6}$ | 1 |
| 11 | *HDAC1* | 1 | 0.07768263 | 0.06808180 | $2.0{\times}10^{-6}$ | 1 |
| 12 | *DENND1A* | 9 | 0.07496250 | 0.22050376 | $2.0{\times}10^{-6}$ | 1 |
| 13 | *ZRANB3* | 2 | −0.07278140 | 0.49384359 | $2.0{\times}10^{-6}$ | 2 |
| 14 | *C16orf70* | 16 | 0.07075091 | 0.43935927 | $3.0{\times}10^{-6}$ | 1 |
| 15 | *EXOC6B* | 2 | 0.06885796 | 0.10195948 | $3.0{\times}10^{-6}$ | 1 |
| 16 | *DIRC3* | 2 | 0.06861035 | 0.13584514 | $3.0{\times}10^{-6}$ | 1 |
| 17 | *C4orf22* | 4 | 0.06847775 | 0.22222222 | $3.0{\times}10^{-6}$ | 1 |
| 18 | *NR6A1* | 9 | 0.06812268 | 0.55860716 | $4.0{\times}10^{-6}$ | 2 |
| 19 | *LRRC29* | 16 | 0.06519534 | 0.13352580 | $6.0{\times}10^{-6}$ | 1 |
| 20 | *TMEM116* | 12 | −0.06340056 | 0.44410413 | $6.0{\times}10^{-6}$ | 1 |
| 21 | *C2CD5* | 12 | −0.06292884 | 0.36420361 | $6.0{\times}10^{-6}$ | 1 |
| 22 | *PRKDC* | 8 | 0.06144151 | 0.09442680 | $7.0{\times}10^{-6}$ | 1 |
| 23 | *PRPF40B* | 12 | 0.05952341 | 0.09292184 | $8.0{\times}10^{-6}$ | 1 |
| 24 | *CELSR3* | 3 | 0.05950715 | 0.15054195 | $8.0{\times}10^{-6}$ | 1 |
| 25 | *MTOR* | 1 | 0.05841437 | 0.34873239 | $1.1{\times}10^{-5}$ | 1 |
| 26 | *ANGPTL7* | 1 | 0.05841437 | 0.34873239 | $1.1{\times}10^{-5}$ | 1 |
| 27 | *KIAA1324L* | 7 | 0.05770516 | 0.16563659 | $1.1{\times}10^{-5}$ | 1 |
| 28 | *EPB41L1* | 20 | 0.05760458 | 0.24740228 | $1.1{\times}10^{-5}$ | 1 |
| 29 | *EPS8* | 12 | 0.05647792 | 0.23065250 | $1.4{\times}10^{-5}$ | 1 |
| 30 | *LOC100188947* | 10 | 0.05452677 | 0.26400000 | $1.8{\times}10^{-5}$ | 1 |
| 31 | *HECTD2* | 10 | 0.05452677 | 0.26400000 | $1.8{\times}10^{-5}$ | 1 |
| 32 | *R3HDM1* | 2 | −0.05448265 | 0.31076443 | $1.8{\times}10^{-5}$ | 1 |
| 33 | *ZNF106* | 15 | 0.05420983 | 0.10975158 | $1.9{\times}10^{-5}$ | 1 |
| 34 | *C8orf44−SGK3* | 8 | 0.05260365 | 0.54661301 | $2.2{\times}10^{-5}$ | 2 |
| 35 | *SGK3* | 8 | 0.05260365 | 0.54661301 | $2.2{\times}10^{-5}$ | 2 |
| 36 | *CCBL2* | 1 | 0.05220339 | 0.37208039 | $2.3{\times}10^{-5}$ | 1 |
| 37 | *LYRM7* | 5 | 0.05213381 | 0.12161647 | $2.4{\times}10^{-5}$ | 1 |
| 38 | *C5orf42* | 5 | 0.05108960 | 0.57542225 | $2.7{\times}10^{-5}$ | 2 |
| 39 | *RCBTB2* | 13 | 0.05067259 | 0.52998066 | $2.8{\times}10^{-5}$ | 2 |
| 40 | *ARL13B* | 3 | 0.05000593 | 0.54782609 | $3.3{\times}10^{-5}$ | 2 |

Table S14: Top 40 SS-G123 sweep candidates at RNA- and protein-coding genes shared between the Central European CEU and West African YRI populations. Candidates presented are those that remained after application of a mappability and alignability filter to data (see *Materials and Methods*). Target genes that pass the significance threshold are colored in gold in the "$P$-value" column. Genes whose sweeps are assigned as hard ($\nu = 1$) are shaded in red in the "Inferred $\nu$" column, while soft sweeps ($\nu \geq 2$) are colored in blue.

| | Top gene | Chromosome | Maximum SS−G123 | G2/G1 | *P*−value | Inferred $\nu$ |
|---|---|---|---|---|---|---|
| 1 | *KIAA0825* | 5 | 0.16262192 | 0.09800692 | $2.0\times10^{-6}$ | 1 |
| 2 | *RP11−554F20.1* | 9 | 0.14328641 | 0.07170707 | $3.0\times10^{-6}$ | 1 |
| 3 | *SPRED3* | 19 | 0.13648875 | 0.04008778 | $3.0\times10^{-6}$ | 1 |
| 4 | *CASC4* | 15 | 0.13642990 | 0.06088150 | $3.0\times10^{-6}$ | 1 |
| 5 | *DEDD* | 1 | 0.12323306 | 0.19270458 | $6.0\times10^{-6}$ | 1 |
| 6 | *GSTT1* | 22 | 0.11632223 | 0.07518328 | $7.0\times10^{-6}$ | 1 |
| 7 | *ENTHD1* | 22 | 0.11574899 | 0.29510665 | $7.0\times10^{-6}$ | 1 |
| 8 | *NNT* | 5 | 0.11500267 | 0.09709843 | $8.0\times10^{-6}$ | 1 |
| 9 | *ATP6V1A* | 3 | 0.11487292 | 0.46656145 | $9.0\times10^{-6}$ | 1 |
| 10 | *DANCR* | 4 | 0.10908564 | 0.16342225 | $1.1\times10^{-5}$ | 1 |
| 11 | *MIR548H3* | 6 | 0.10678068 | 0.11806435 | $1.5\times10^{-5}$ | 1 |
| 12 | *GRIA2* | 4 | 0.10578392 | 0.36923077 | $1.7\times10^{-5}$ | 1 |
| 13 | *GRIK5* | 19 | 0.10082463 | 0.14846599 | $1.9\times10^{-5}$ | 1 |
| 14 | *TYW5* | 2 | 0.10050784 | 0.52092831 | $1.9\times10^{-5}$ | 1 |
| 15 | *PIGV* | 1 | 0.09740419 | 0.10112360 | $2.3\times10^{-5}$ | 1 |
| 16 | *RALGAPA2* | 20 | 0.09590598 | 0.09440637 | $2.7\times10^{-5}$ | 1 |
| 17 | *PLEKHA8* | 7 | 0.09390530 | 0.29760666 | $3.3\times10^{-5}$ | 1 |
| 18 | *GLRX2* | 1 | 0.09374561 | 0.53420670 | $3.3\times10^{-5}$ | 1 |
| 19 | *PAWR* | 12 | −0.09282636 | 0.41657077 | $3.4\times10^{-5}$ | 1 |
| 20 | *PHKB* | 16 | 0.09044005 | 0.10755993 | $3.6\times10^{-5}$ | 1 |
| 21 | *ERLIN2* | 8 | 0.08847504 | 0.09065880 | $3.9\times10^{-5}$ | 1 |
| 22 | *SPG11* | 15 | 0.08398508 | 0.14916585 | $5.5\times10^{-5}$ | 1 |
| 23 | *RRN3P3* | 16 | −0.08384446 | 0.67088036 | $5.6\times10^{-5}$ | 8 |
| 24 | *DKK2* | 4 | 0.08252351 | 0.13276618 | $6.1\times10^{-5}$ | 1 |
| 25 | *ABHD17B* | 9 | 0.08249391 | 0.08387291 | $6.1\times10^{-5}$ | 1 |
| 26 | *CNNM2* | 10 | 0.08094510 | 0.15631825 | $6.4\times10^{-5}$ | 1 |
| 27 | *TGFBR1* | 9 | 0.07873090 | 0.42549372 | $7.4\times10^{-5}$ | 1 |
| 28 | *CPNE1* | 20 | 0.07384179 | 0.19304797 | $1.03\times10^{-4}$ | 1 |
| 29 | *PRKAR2A* | 3 | 0.07271481 | 0.59669161 | $1.14\times10^{-4}$ | 2 |
| 30 | *NLK* | 17 | 0.07079549 | 0.24630342 | $1.30\times10^{-4}$ | 1 |
| 31 | *CDK6* | 7 | 0.07054583 | 0.07748561 | $1.32\times10^{-4}$ | 1 |
| 32 | *SYT1* | 12 | 0.06986742 | 0.27157762 | $1.38\times10^{-4}$ | 1 |
| 33 | *GABRA4* | 4 | 0.06971220 | 0.52556668 | $1.38\times10^{-4}$ | 1 |
| 34 | *ZNF451* | 6 | 0.06885744 | 0.19779485 | $1.46\times10^{-4}$ | 1 |
| 35 | *ITFG1* | 16 | 0.06882713 | 0.16926617 | $1.46\times10^{-4}$ | 1 |
| 36 | *FRYL* | 4 | 0.06828603 | 0.56018307 | $1.52\times10^{-4}$ | 2 |
| 37 | *EXOC6B* | 2 | −0.06782534 | 0.46576537 | $1.62\times10^{-4}$ | 1 |
| 38 | *USP46* | 4 | 0.06639854 | 0.25667307 | $1.75\times10^{-4}$ | 1 |
| 39 | *LGSN* | 6 | 0.06638067 | 0.19928826 | $1.76\times10^{-4}$ | 1 |
| 40 | *ATF2* | 2 | 0.06592706 | 0.26808614 | $1.83\times10^{-4}$ | 1 |

Table S15: Top 40 SS-G123 sweep candidates at RNA- and protein-coding genes shared between the East African LWK and West African YRI populations. Candidates presented are those that remained after application of a mappability and alignability filter to data (see *Materials and Methods*). Target genes that pass the significance threshold are colored in gold in the "$P$-value" column. Genes whose sweeps are assigned as hard ($\nu = 1$) are shaded in red in the "Inferred $\nu$" column, while soft sweeps ($\nu \geq 2$) are colored in blue.

| | Top gene | Chromosome | Maximum SS−G123 | G2/G1 | *P*−value | Inferred $\nu$ |
|---|---|---|---|---|---|---|
| 1 | *GRIK5* | 19 | 0.2415148 | 0.12482020 | $< 10^{-6}$ | 1 |
| 2 | *GSTT1* | 22 | 0.2247992 | 0.23472029 | $< 10^{-6}$ | 3 |
| 3 | *RP11−554F20.1* | 9 | 0.2067291 | 0.10210615 | $< 10^{-6}$ | 1 |
| 4 | *KIAA0825* | 5 | 0.2003773 | 0.29785697 | $< 10^{-6}$ | 2 |
| 5 | *RGS18* | 1 | 0.1739911 | 0.21109123 | $< 10^{-6}$ | 2 |
| 6 | *RRN3P3* | 16 | 0.1675603 | 0.40424178 | $< 10^{-6}$ | 2 |
| 7 | *PPAPDC1B* | 8 | 0.1654046 | 0.11850615 | $< 10^{-6}$ | 1 |
| 8 | *SPRED3* | 19 | 0.1609167 | 0.05899437 | $< 10^{-6}$ | 1 |
| 9 | *NNT* | 5 | 0.1564619 | 0.09173050 | $< 10^{-6}$ | 1 |
| 10 | *ARID1A* | 1 | 0.1540840 | 0.14330287 | $< 10^{-6}$ | 1 |
| 11 | *CASC4* | 15 | 0.1429575 | 0.05820969 | $< 10^{-6}$ | 1 |
| 12 | *EHBP1* | 2 | 0.1429198 | 0.17335159 | $< 10^{-6}$ | 1 |
| 13 | *ATF2* | 2 | 0.1396078 | 0.54310604 | $< 10^{-6}$ | 2 |
| 14 | *DKK2* | 4 | 0.1378883 | 0.12261259 | $< 10^{-6}$ | 1 |
| 15 | *MIR548AE2* | 16 | 0.1372927 | 0.09054441 | $< 10^{-6}$ | 1 |
| 16 | *LONP2* | 16 | 0.1372927 | 0.09054441 | $< 10^{-6}$ | 1 |
| 17 | *DDHD2* | 8 | 0.1343500 | 0.11662586 | $< 10^{-6}$ | 1 |
| 18 | *PHKB* | 16 | 0.1315935 | 0.39985719 | $< 10^{-6}$ | 2 |
| 19 | *FER1L6* | 8 | 0.1274631 | 0.42668677 | $< 10^{-6}$ | 2 |
| 20 | *FER1L6−AS1* | 8 | 0.1274631 | 0.42668677 | $< 10^{-6}$ | 2 |
| 21 | *SPIDR* | 8 | 0.1269844 | 0.47826087 | $< 10^{-6}$ | 2 |
| 22 | *GRIA4* | 11 | 0.1268945 | 0.16182811 | $< 10^{-6}$ | 1 |
| 23 | *DANCR* | 4 | 0.1260421 | 0.15609221 | $< 10^{-6}$ | 1 |
| 24 | *TCF4* | 18 | 0.1243683 | 0.20357244 | $< 10^{-6}$ | 1 |
| 25 | *PIGV* | 1 | 0.1237784 | 0.09449782 | $< 10^{-6}$ | 1 |
| 26 | *RALGAPA2* | 20 | 0.1233941 | 0.09403724 | $< 10^{-6}$ | 1 |
| 27 | *TMC1* | 9 | 0.1218435 | 0.28051683 | $< 10^{-6}$ | 1 |
| 28 | *PSMD14* | 2 | 0.1205903 | 0.43648469 | $< 10^{-6}$ | 2 |
| 29 | *CNNM2* | 10 | 0.1205598 | 0.13187097 | $< 10^{-6}$ | 1 |
| 30 | *FAM172A* | 5 | 0.1204877 | 0.46349206 | $< 10^{-6}$ | 2 |
| 31 | *BRIP1* | 17 | 0.1186024 | 0.25193076 | $< 10^{-6}$ | 1 |
| 32 | *GSTTP2* | 22 | 0.1184177 | 0.16994022 | $< 10^{-6}$ | 1 |
| 33 | *MAGI3* | 1 | 0.1173924 | 0.17789681 | $< 10^{-6}$ | 1 |
| 34 | *METTL25* | 12 | 0.1168971 | 0.23853844 | $< 10^{-6}$ | 1 |
| 35 | *CCDC178* | 18 | 0.1152967 | 0.27341227 | $< 10^{-6}$ | 1 |
| 36 | *DDX19B* | 16 | 0.1146779 | 0.23421127 | $< 10^{-6}$ | 1 |
| 37 | *RBBP4* | 1 | 0.1145007 | 0.63782696 | $< 10^{-6}$ | 2 |
| 38 | *C10orf32−ASMT* | 10 | 0.1138325 | 0.38839475 | $< 10^{-6}$ | 2 |
| 39 | *AS3MT* | 10 | 0.1138325 | 0.38839475 | $< 10^{-6}$ | 2 |
| 40 | *ZNF451* | 6 | 0.1137655 | 0.21774436 | $< 10^{-6}$ | 1 |

Table S16: Top 40 SS-G123 sweep candidates at RNA- and protein-coding genes shared between the South Asian GIH and West African YRI populations. Candidates presented are those that remained after application of a mappability and alignability filter to data (see *Materials and Methods*). Target genes that pass the significance threshold are colored in gold in the "$P$-value" column. Genes whose sweeps are assigned as hard ($\nu = 1$) are shaded in red in the "Inferred $\nu$" column, while soft sweeps ($\nu \geq 2$) are colored in blue.

| | Top gene | Chromosome | Maximum SS−G123 | G2/G1 | *P*−value | Inferred $\nu$ |
|---|---|---|---|---|---|---|
| 1 | *CASC4* | 15 | 0.16711386 | 0.03621655 | $< 10^{-6}$ | 1 |
| 2 | *NNT* | 5 | 0.14575420 | 0.04650487 | $< 10^{-6}$ | 1 |
| 3 | *KIAA0825* | 5 | 0.14498089 | 0.07326733 | $< 10^{-6}$ | 1 |
| 4 | *CDK6* | 7 | 0.13787179 | 0.23060483 | $< 10^{-6}$ | 1 |
| 5 | *ATP6V1A* | 3 | 0.12981556 | 0.28721584 | $< 10^{-6}$ | 1 |
| 6 | *DDHD2* | 8 | 0.12639095 | 0.06514875 | $< 10^{-6}$ | 1 |
| 7 | *SYT1* | 12 | 0.12432490 | 0.04995287 | $< 10^{-6}$ | 1 |
| 8 | *DEDD* | 1 | 0.12425622 | 0.15489439 | $< 10^{-6}$ | 1 |
| 9 | *RP11−554F20.1* | 9 | 0.12160852 | 0.05718271 | $< 10^{-6}$ | 1 |
| 10 | *PSMD14* | 2 | 0.11708713 | 0.43952713 | $< 10^{-6}$ | 1 |
| 11 | *FBXW4* | 10 | 0.11552344 | 0.12795875 | $< 10^{-6}$ | 1 |
| 12 | *PIGV* | 1 | 0.11324551 | 0.10666977 | $< 10^{-6}$ | 1 |
| 13 | *MIR548AE2* | 16 | 0.10772443 | 0.11468505 | $< 10^{-6}$ | 1 |
| 14 | *LONP2* | 16 | 0.10772443 | 0.11468505 | $< 10^{-6}$ | 1 |
| 15 | *ABHD17B* | 9 | 0.10372984 | 0.05366309 | $< 10^{-6}$ | 1 |
| 16 | *GRIK5* | 19 | 0.10341289 | 0.11576916 | $< 10^{-6}$ | 1 |
| 17 | *PPAPDC1B* | 8 | 0.10317107 | 0.12980421 | $< 10^{-6}$ | 1 |
| 18 | *C10orf76* | 10 | 0.10178019 | 0.51611553 | $< 10^{-6}$ | 1 |
| 19 | *PLEKHA8* | 7 | 0.10177351 | 0.30315058 | $< 10^{-6}$ | 1 |
| 20 | *PHKB* | 16 | 0.09939696 | 0.15921020 | $< 10^{-6}$ | 1 |
| 21 | *RALGAPA2* | 20 | 0.09639928 | 0.13588492 | $< 10^{-6}$ | 1 |
| 22 | *TULP4* | 6 | 0.09480228 | 0.14873493 | $< 10^{-6}$ | 1 |
| 23 | *ERLIN2* | 8 | 0.09254210 | 0.13007499 | $< 10^{-6}$ | 1 |
| 24 | *EXOC6B* | 2 | −0.09028512 | 0.55109110 | $1.0{\times}10^{-6}$ | 2 |
| 25 | *ZNF451* | 6 | 0.08854748 | 0.09354446 | $1.0{\times}10^{-6}$ | 1 |
| 26 | *CADM1* | 11 | 0.08766952 | 0.35196436 | $1.0{\times}10^{-6}$ | 1 |
| 27 | *COL7A1* | 3 | 0.08603112 | 0.12098249 | $1.0{\times}10^{-6}$ | 1 |
| 28 | *PAWR* | 12 | −0.08580547 | 0.44387569 | $1.0{\times}10^{-6}$ | 1 |
| 29 | *CAMK2G* | 10 | 0.08306775 | 0.43884563 | $2.0{\times}10^{-6}$ | 1 |
| 30 | *FRYL* | 4 | 0.08153959 | 0.50101833 | $2.0{\times}10^{-6}$ | 1 |
| 31 | *SIRPB1* | 20 | 0.07908564 | 0.23271969 | $2.0{\times}10^{-6}$ | 1 |
| 32 | *USP46* | 4 | 0.07871925 | 0.11263519 | $2.0{\times}10^{-6}$ | 1 |
| 33 | *SLC4A10* | 2 | 0.07705620 | 0.12719176 | $2.0{\times}10^{-6}$ | 1 |
| 34 | *CPNE1* | 20 | 0.07523397 | 0.32026984 | $2.0{\times}10^{-6}$ | 1 |
| 35 | *FER1L6* | 8 | 0.07413957 | 0.50163815 | $2.0{\times}10^{-6}$ | 1 |
| 36 | *FER1L6−AS1* | 8 | 0.07413957 | 0.50163815 | $2.0{\times}10^{-6}$ | 1 |
| 37 | *UBE4A* | 11 | 0.07396793 | 0.50684932 | $2.0{\times}10^{-6}$ | 1 |
| 38 | *LOC100131626* | 11 | 0.07396793 | 0.50684932 | $2.0{\times}10^{-6}$ | 1 |
| 39 | *NLK* | 17 | 0.07380719 | 0.33165253 | $2.0{\times}10^{-6}$ | 1 |
| 40 | *ENTHD1* | 22 | 0.07378007 | 0.20033285 | $2.0{\times}10^{-6}$ | 1 |

Table S17: Top 40 SS-G123 sweep candidates at RNA- and protein-coding genes shared between the East Asian JPT and West African YRI populations. Candidates presented are those that remained after application of a mappability and alignability filter to data (see *Materials and Methods*). Target genes that pass the significance threshold are colored in gold in the "$P$-value" column. Genes whose sweeps are assigned as hard ($\nu = 1$) are shaded in red in the "Inferred $\nu$" column, while soft sweeps ($\nu \geq 2$) are colored in blue.

| | Top gene | Chromosome | Maximum SS−G123 | G2/G1 | $P$−value | Inferred $\nu$ |
|---|---|---|---|---|---|---|
| *1* | *UCHL5* | 1 | 0.16261045 | 0.54330176 | $< 10^{-6}$ | 2 |
| *2* | *CASC4* | 15 | 0.15414538 | 0.02859798 | $< 10^{-6}$ | 1 |
| *3* | *GRIK5* | 19 | 0.15077705 | 0.08571429 | $< 10^{-6}$ | 1 |
| *4* | *ATP6V1A* | 3 | 0.14779863 | 0.51696607 | $< 10^{-6}$ | 2 |
| *5* | *SIRPB1* | 20 | 0.14373282 | 0.17088899 | $< 10^{-6}$ | 1 |
| *6* | *HEMGN* | 9 | 0.14078020 | 0.07985481 | $< 10^{-6}$ | 1 |
| *7* | *KIAA0825* | 5 | 0.13892611 | 0.14634146 | $< 10^{-6}$ | 1 |
| *8* | *EHBP1* | 2 | 0.13344376 | 0.15266860 | $< 10^{-6}$ | 1 |
| *9* | *RP11−554F20.1* | 9 | 0.12449166 | 0.28106373 | $< 10^{-6}$ | 1 |
| *10* | *ENTHD1* | 22 | 0.11764075 | 0.12615588 | $< 10^{-6}$ | 1 |
| *11* | *PSMD14* | 2 | 0.10605231 | 0.32749562 | $< 10^{-6}$ | 1 |
| *12* | *NNT* | 5 | 0.10287121 | 0.02879851 | $< 10^{-6}$ | 1 |
| *13* | *EXOC6B* | 2 | −0.09657734 | 0.32384649 | $< 10^{-6}$ | 1 |
| *14* | *DDHD2* | 8 | 0.09268298 | 0.17119048 | $< 10^{-6}$ | 1 |
| *15* | *MIR548AE2* | 16 | 0.09252061 | 0.04543263 | $< 10^{-6}$ | 1 |
| *16* | *LONP2* | 16 | 0.09252061 | 0.04543263 | $< 10^{-6}$ | 1 |
| *17* | *PIGV* | 1 | 0.09246271 | 0.08195141 | $< 10^{-6}$ | 1 |
| *18* | *GRIA4* | 11 | 0.09091467 | 0.24805339 | $< 10^{-6}$ | 1 |
| *19* | *RALGAPA2* | 20 | 0.09018390 | 0.06541614 | $< 10^{-6}$ | 1 |
| *20* | *CSMD3* | 8 | 0.08642643 | 0.10669975 | $< 10^{-6}$ | 1 |
| *21* | *AUTS2* | 7 | 0.08609979 | 0.23589894 | $< 10^{-6}$ | 1 |
| *22* | *MPHOSPH9* | 12 | −0.08463466 | 0.52033195 | $< 10^{-6}$ | 1 |
| *23* | *PPAPDC1B* | 8 | 0.08450565 | 0.28654354 | $< 10^{-6}$ | 1 |
| *24* | *MIR548H3* | 6 | 0.08341730 | 0.14782609 | $< 10^{-6}$ | 1 |
| *25* | *ABCB1* | 7 | 0.08225091 | 0.50471294 | $< 10^{-6}$ | 1 |
| *26* | *RUNDC3B* | 7 | 0.08225091 | 0.50471294 | $< 10^{-6}$ | 1 |
| *27* | *SIPA1L3* | 19 | 0.08082831 | 0.16331995 | $< 10^{-6}$ | 1 |
| *28* | *NLK* | 17 | 0.08023741 | 0.25966716 | $< 10^{-6}$ | 1 |
| *29* | *CPNE1* | 20 | 0.07997871 | 0.11175373 | $< 10^{-6}$ | 1 |
| *30* | *GRIA2* | 4 | 0.07905065 | 0.40514076 | $< 10^{-6}$ | 1 |
| *31* | *CDK6* | 7 | 0.07856601 | 0.11933861 | $< 10^{-6}$ | 1 |
| *32* | *PLEKHA8* | 7 | 0.07799249 | 0.29088472 | $< 10^{-6}$ | 1 |
| *33* | *DKK2* | 4 | 0.07704189 | 0.07856381 | $< 10^{-6}$ | 1 |
| *34* | *SYT1* | 12 | 0.07684115 | 0.12255668 | $< 10^{-6}$ | 1 |
| *35* | *COL7A1* | 3 | 0.07658377 | 0.07582045 | $< 10^{-6}$ | 1 |
| *36* | *GSTT1* | 22 | 0.07574164 | 0.10880196 | $< 10^{-6}$ | 1 |
| *37* | *FBXW4* | 10 | 0.07431220 | 0.15773660 | $1.0 \times 10^{-6}$ | 1 |
| *38* | *USP46* | 4 | 0.07174521 | 0.19367210 | $1.0 \times 10^{-6}$ | 1 |
| *39* | *AC016582.2* | 19 | 0.07004390 | 0.18775372 | $1.0 \times 10^{-6}$ | 1 |
| *40* | *ARID1A* | 1 | 0.06960214 | 0.02739301 | $1.0 \times 10^{-6}$ | 1 |

Table S18: Top 40 SS-G123 sweep candidates at RNA- and protein-coding genes shared between the East Asian JPT and South Asian GIH populations. Candidates presented are those that remained after application of a mappability and alignability filter to data (see *Materials and Methods*). Target genes that pass the significance threshold are colored in gold in the "$P$-value" column. Genes whose sweeps are assigned as hard ($\nu = 1$) are shaded in red in the "Inferred $\nu$" column, while soft sweeps ($\nu \geq 2$) are colored in blue.

| | Top gene | Chromosome | Maximum SS−G123 | G2/G1 | *P*−value | Inferred $\nu$ |
|---|---|---|---|---|---|---|
| 1 | *RUNX1T1* | 8 | 0.19081280 | 0.08034745 | $< 10^{-6}$ | 1 |
| 2 | *P4HTM* | 3 | 0.17561530 | 0.02724586 | $< 10^{-6}$ | 1 |
| 3 | *EXOC6B* | 2 | 0.11912726 | 0.09503897 | $< 10^{-6}$ | 1 |
| 4 | *TMTC2* | 12 | 0.11638215 | 0.18265334 | $< 10^{-6}$ | 1 |
| 5 | *SUGCT* | 7 | 0.11594031 | 0.12736706 | $< 10^{-6}$ | 1 |
| 6 | *HELZ* | 17 | 0.11386077 | 0.35123207 | $< 10^{-6}$ | 2 |
| 7 | *DBT* | 1 | 0.11216524 | 0.06377143 | $< 10^{-6}$ | 1 |
| 8 | *BCAS3* | 17 | 0.10947050 | 0.06812121 | $< 10^{-6}$ | 1 |
| 9 | *RP11−53O19.1* | 5 | 0.10879938 | 0.52299606 | $< 10^{-6}$ | 2 |
| 10 | *ZNF546* | 19 | 0.09867813 | 0.13472000 | $< 10^{-6}$ | 1 |
| 11 | *ADAMTS6* | 5 | 0.09839895 | 0.49891634 | $< 10^{-6}$ | 2 |
| 12 | *APPBP2* | 17 | 0.09796838 | 0.09183407 | $< 10^{-6}$ | 1 |
| 13 | *DR1* | 1 | 0.09702526 | 0.42016095 | $< 10^{-6}$ | 1 |
| 14 | *USP25* | 21 | 0.09547922 | 0.16971106 | $< 10^{-6}$ | 1 |
| 15 | *RNF121* | 11 | 0.09440660 | 0.09574218 | $< 10^{-6}$ | 1 |
| 16 | *NEGR1* | 1 | 0.09420501 | 0.28005997 | $< 10^{-6}$ | 1 |
| 17 | *RNU6−28P* | 15 | 0.09050468 | 0.27059635 | $< 10^{-6}$ | 1 |
| 18 | *PPIP5K1* | 15 | 0.09050468 | 0.27059635 | $< 10^{-6}$ | 1 |
| 19 | *ORC2* | 2 | 0.08524372 | 0.07114440 | $< 10^{-6}$ | 1 |
| 20 | *HDAC1* | 1 | 0.08444452 | 0.12845175 | $< 10^{-6}$ | 1 |
| 21 | *USP32* | 17 | 0.08419480 | 0.14071582 | $< 10^{-6}$ | 1 |
| 22 | *CCDC18* | 1 | 0.08341467 | 0.34291581 | $< 10^{-6}$ | 1 |
| 23 | *LYRM7* | 5 | 0.08066234 | 0.07455296 | $< 10^{-6}$ | 1 |
| 24 | *LOC100188947* | 10 | 0.08039944 | 0.38098427 | $< 10^{-6}$ | 1 |
| 25 | *HECTD2* | 10 | 0.08039944 | 0.38098427 | $< 10^{-6}$ | 1 |
| 26 | *SSH2* | 17 | 0.07918183 | 0.29931234 | $< 10^{-6}$ | 1 |
| 27 | *ZNF780B* | 19 | 0.07884324 | 0.14236707 | $< 10^{-6}$ | 1 |
| 28 | *NLK* | 17 | 0.07866030 | 0.15058966 | $< 10^{-6}$ | 1 |
| 29 | *MTF2* | 1 | 0.07827113 | 0.56458636 | $< 10^{-6}$ | 2 |
| 30 | *ATP6V0D1* | 16 | 0.07404407 | 0.08471815 | $< 10^{-6}$ | 1 |
| 31 | *MBOAT2* | 2 | 0.07358927 | 0.11950505 | $< 10^{-6}$ | 1 |
| 32 | *BCL2L1* | 20 | 0.07326540 | 0.13298940 | $< 10^{-6}$ | 1 |
| 33 | *CENPW* | 6 | 0.07324676 | 0.21777778 | $< 10^{-6}$ | 1 |
| 34 | *BEND4* | 4 | 0.07308935 | 0.08658982 | $< 10^{-6}$ | 1 |
| 35 | *LINC00478* | 21 | 0.07294715 | 0.17406645 | $< 10^{-6}$ | 1 |
| 36 | *XXYLT1* | 3 | 0.07133674 | 0.10906096 | $< 10^{-6}$ | 1 |
| 37 | *ABCB1* | 7 | 0.07111757 | 0.14704879 | $< 10^{-6}$ | 1 |
| 38 | *RUNDC3B* | 7 | 0.07111757 | 0.14704879 | $< 10^{-6}$ | 1 |
| 39 | *NGLY1* | 3 | 0.06960764 | 0.15325248 | $< 10^{-6}$ | 1 |
| 40 | *CELSR3* | 3 | 0.06809979 | 0.09836612 | $< 10^{-6}$ | 1 |

Table S19: Top 40 SS-G123 sweep candidates at RNA- and protein-coding genes shared between the East Asian JPT and Southeast Asian KHV populations. Candidates presented are those that remained after application of a mappability and alignability filter to data (see *Materials and Methods*). Target genes that pass the significance threshold are colored in gold in the "$P$-value" column. Genes whose sweeps are assigned as hard ($\nu = 1$) are shaded in red in the "Inferred $\nu$" column, while soft sweeps ($\nu \geq 2$) are colored in blue.

| | Top gene | Chromosome | Maximum SS−G123 | G2/G1 | *P*−value | Inferred $\nu$ |
|---|---|---|---|---|---|---|
| *1* | *SPIDR* | 8 | 0.2784003 | 0.38334810 | $< 10^{-6}$ | 2 |
| *2* | *RP11−696N14.1* | 4 | 0.2744507 | 0.20046722 | $< 10^{-6}$ | 2 |
| *3* | *EXOC6B* | 2 | 0.2617294 | 0.03631803 | $< 10^{-6}$ | 1 |
| *4* | *RUNX1T1* | 8 | 0.2599384 | 0.06254055 | $< 10^{-6}$ | 1 |
| *5* | *SPAG6* | 10 | 0.2444433 | 0.03124657 | $< 10^{-6}$ | 1 |
| *6* | *EXD2* | 14 | 0.2338336 | 0.06551773 | $< 10^{-6}$ | 1 |
| *7* | *P4HTM* | 3 | 0.2265336 | 0.02184214 | $< 10^{-6}$ | 1 |
| *8* | *AMBRA1* | 11 | 0.2041819 | 0.07887964 | $< 10^{-6}$ | 1 |
| *9* | *BCL7C* | 16 | 0.2019106 | 0.04684318 | $< 10^{-6}$ | 1 |
| *10* | *BCL2L1* | 20 | 0.1971090 | 0.09232554 | $< 10^{-6}$ | 1 |
| *11* | *ADH1A* | 4 | 0.1963534 | 0.27346115 | $< 10^{-6}$ | 2 |
| *12* | *PGAP1* | 2 | 0.1945704 | 0.09764099 | $< 10^{-6}$ | 1 |
| *13* | *C11orf49* | 11 | 0.1898302 | 0.18259828 | $3.0\times10^{-6}$ | 2 |
| *14* | *PRPF40B* | 12 | 0.1861633 | 0.03307622 | $6.0\times10^{-6}$ | 1 |
| *15* | *LOC100188947* | 10 | 0.1858056 | 0.47456664 | $6.0\times10^{-6}$ | 2 |
| *16* | *HECTD2* | 10 | 0.1858056 | 0.47456664 | $6.0\times10^{-6}$ | 2 |
| *17* | *NOVA1* | 14 | 0.1830759 | 0.49632253 | $7.0\times10^{-6}$ | 2 |
| *18* | *ZNF282* | 7 | 0.1813378 | 0.05366309 | $7.0\times10^{-6}$ | 1 |
| *19* | *HMCN1* | 1 | 0.1764870 | 0.47648708 | $8.0\times10^{-6}$ | 2 |
| *20* | *PHYHIPL* | 10 | 0.1743157 | 0.07840000 | $8.0\times10^{-6}$ | 1 |
| *21* | *LINC01024* | 5 | 0.1742617 | 0.13723019 | $8.0\times10^{-6}$ | 1 |
| *22* | *ARIH2* | 3 | 0.1734230 | 0.06296852 | $8.0\times10^{-6}$ | 1 |
| *23* | *PURA* | 5 | 0.1727126 | 0.13155916 | $9.0\times10^{-6}$ | 1 |
| *24* | *ARIH2OS* | 3 | 0.1680339 | 0.05414495 | $1.2\times10^{-5}$ | 1 |
| *25* | *TRUB1* | 10 | 0.1677889 | 0.06171946 | $1.2\times10^{-5}$ | 1 |
| *26* | *CEP112* | 17 | 0.1663139 | 0.06596257 | $1.4\times10^{-5}$ | 1 |
| *27* | *USP38* | 4 | 0.1643823 | 0.14698163 | $1.5\times10^{-5}$ | 1 |
| *28* | *C2orf66* | 2 | 0.1629327 | 0.04013059 | $1.5\times10^{-5}$ | 1 |
| *29* | *LINC01088* | 4 | 0.1622231 | 0.12088799 | $1.5\times10^{-5}$ | 1 |
| *30* | *TRMT11* | 6 | 0.1609308 | 0.04417217 | $1.6\times10^{-5}$ | 1 |
| *31* | *CCDC18* | 1 | 0.1591868 | 0.39498058 | $1.6\times10^{-5}$ | 2 |
| *32* | *LINC00536* | 8 | 0.1575138 | 0.53060485 | $1.6\times10^{-5}$ | 3 |
| *33* | *CSPP1* | 8 | 0.1570753 | 0.46273402 | $1.6\times10^{-5}$ | 2 |
| *34* | *RAD51B* | 14 | 0.1546937 | 0.07764864 | $1.6\times10^{-5}$ | 1 |
| *35* | *SPATS2* | 12 | 0.1543295 | 0.07337854 | $1.6\times10^{-5}$ | 1 |
| *36* | *FBXO4* | 5 | 0.1535929 | 0.03908674 | $1.6\times10^{-5}$ | 1 |
| *37* | *PKIA* | 8 | 0.1529611 | 0.23342175 | $1.7\times10^{-5}$ | 2 |
| *38* | *GPHN* | 14 | 0.1511221 | 0.41336881 | $1.7\times10^{-5}$ | 2 |
| *39* | *ZNF660* | 3 | 0.1509682 | 0.12457299 | $1.7\times10^{-5}$ | 1 |
| *40* | *OXCT1* | 5 | 0.1500597 | 0.04652922 | $1.8\times10^{-5}$ | 1 |

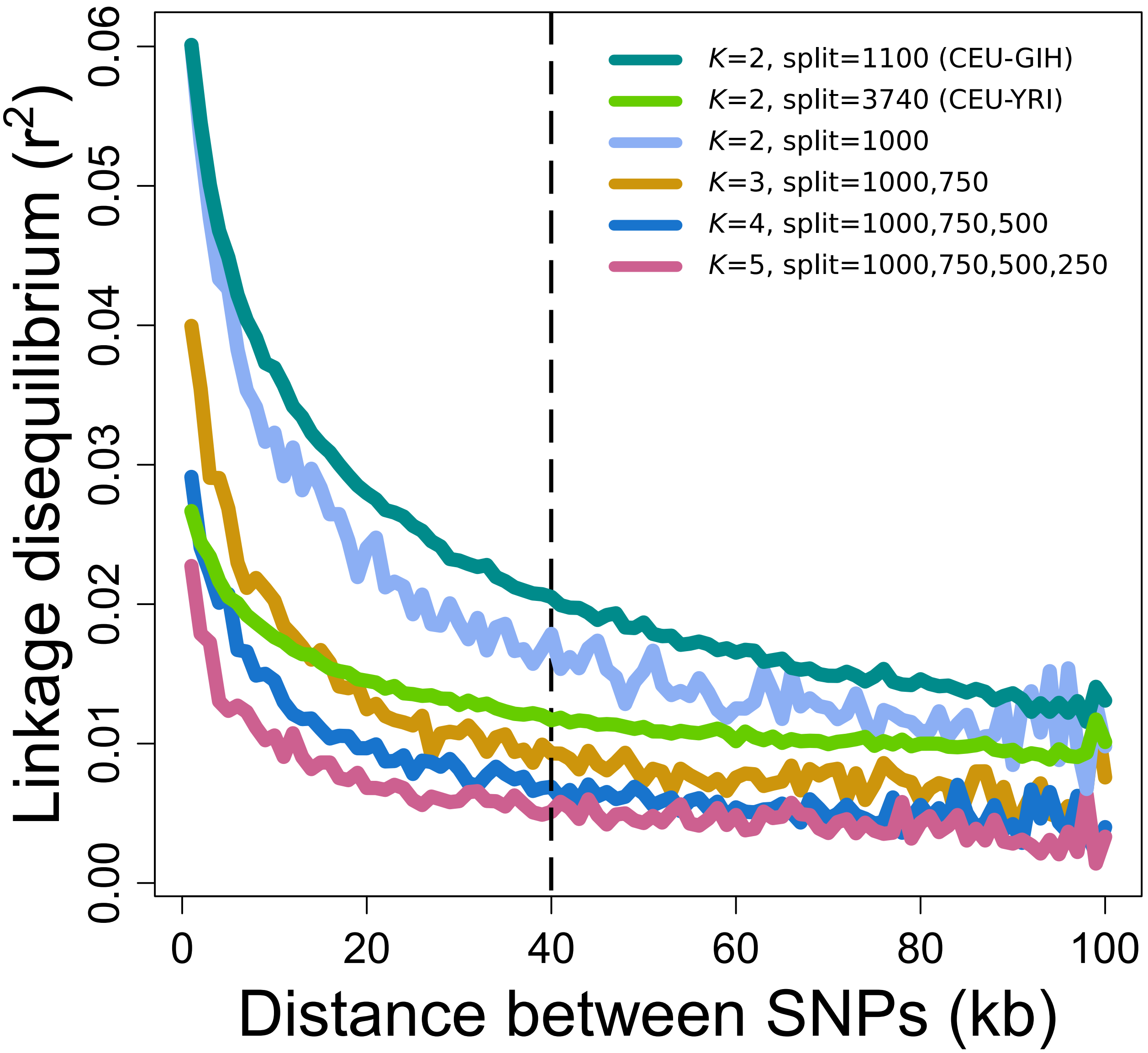


Figure S1: Decay of pairwise LD measured as $r^2$ over 100 kb for pooled samples simulated under neutrality containing two to five populations. For pairs of populations (cyan, chartreuse, and sky blue), we simulated under the CEU-GIH ($\tau = 1100$) or CEU-YRI ($\tau = 3740$) human demographic models, or a simplified constant demographic history model ($\tau = 1000$) with $N = 10^4$, where the ancestral population splits once. For trios of populations (goldenrod), two splits underlie the sample phylogeny, occurring $\tau_1 = 1000$ and $\tau_2 = 750$ generations before sampling. For quartets of populations (dark blue), there are three splits occurring at $\tau_1 = 1000$, $\tau_2 = 750$, and $\tau_3 = 500$ generations before sampling. Finally, $K = 5$ phylogenies result from four splits, at $\tau_1 = 1000$ $\tau_2 = 750$, $\tau_3 = 500$, and $\tau_4 = 250$ generations before sampling. All samples are identical to those analyzed in the main text.

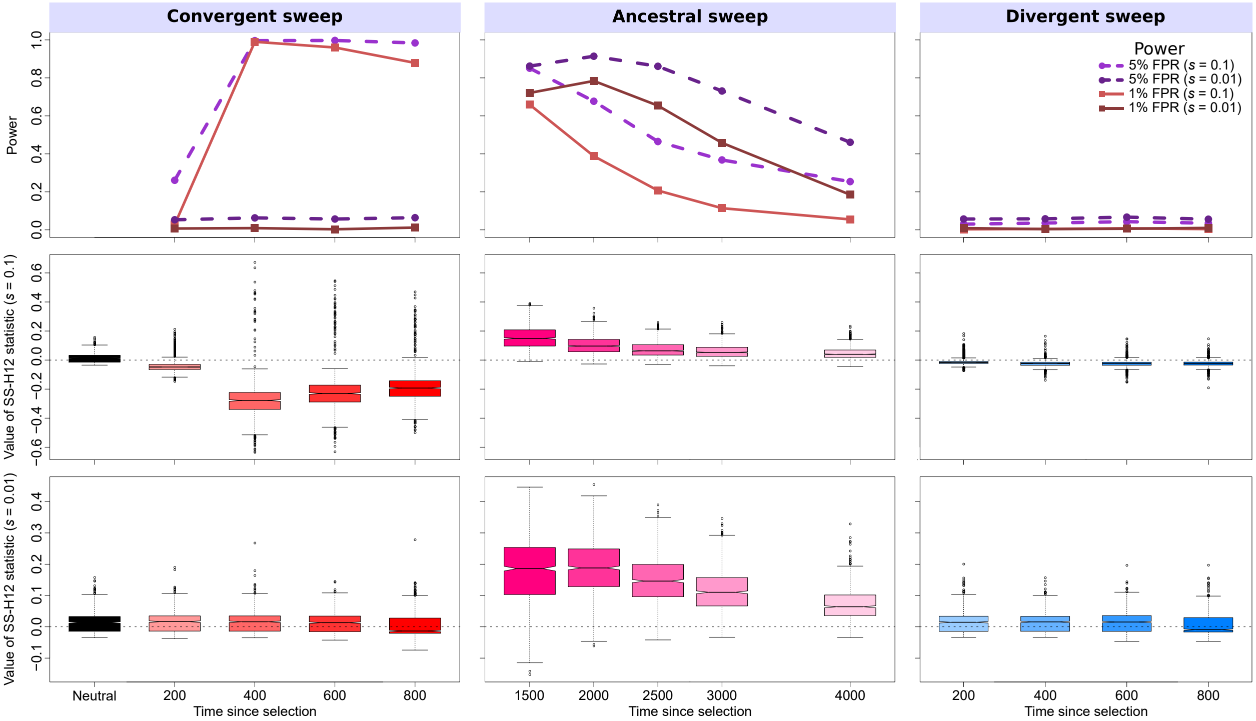


Figure S2: Properties of SS-H12 for simulated strong ($s = 0.1$) and moderate ($s = 0.01$) soft sweep ($\nu = 4$) scenarios under the CEU-GIH model ($\tau = 1100$ generations before sampling). (Top row) Power at 1% (red lines) and 5% (purple lines) false positive rates (FPRs) to detect recent ancestral, convergent, and divergent soft sweeps from selection on standing genetic variation as a function of time at which selection of the favored haplotypes initiated ($t$), with FPR based on the distribution of maximum |SS-H12| across simulated neutral replicates. (Middle row) Box plots summarizing the distribution of SS-H12 values from windows of maximum |SS-H12| across strong sweep replicates, corresponding to each time point in the power curves, with dashed lines in each panel representing SS-H12 = 0. (Bottom row) Box plots summarizing the distribution of SS-H12 values across moderate sweep replicates. For convergent and divergent sweeps, $t < \tau$, while for ancestral sweeps, $t > \tau$. All replicate samples for the CEU-GIH model contain 99 simulated CEU individuals and 103 simulated GIH individuals, as in the 1000 Genomes Project dataset [Auton et al., 2015], and we performed 1000 replicates for each scenario.

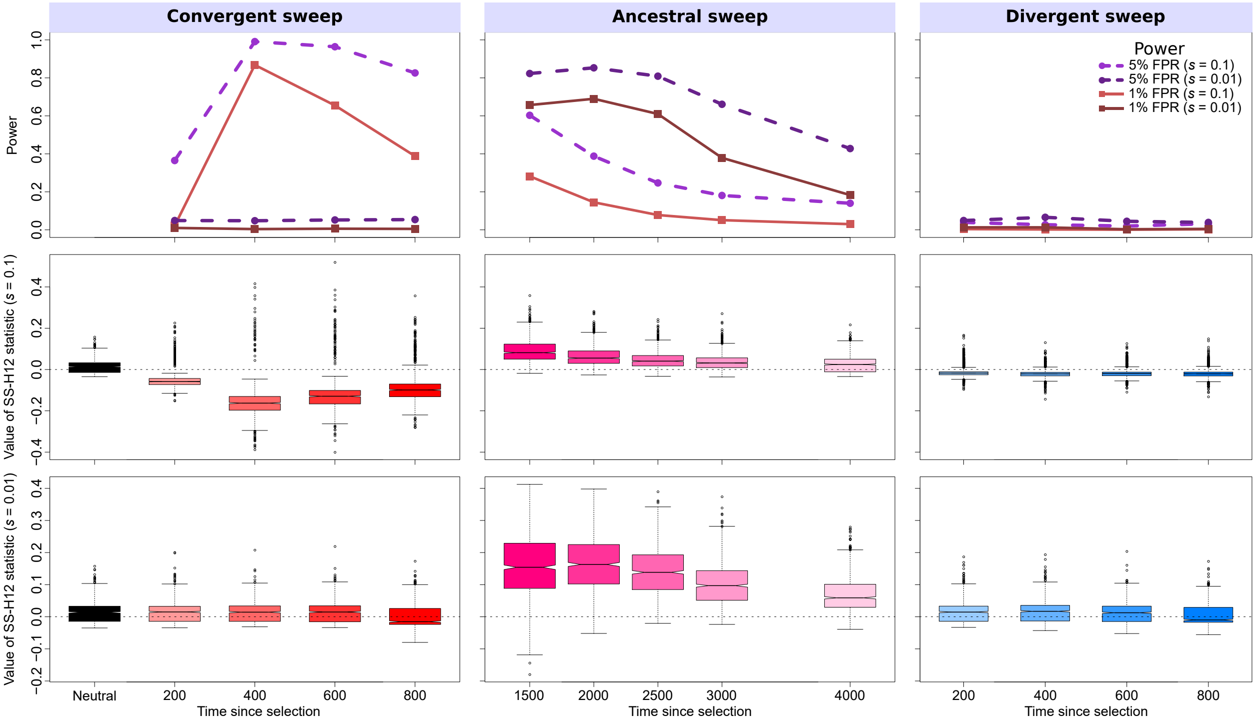


Figure S3: Properties of SS-H12 for simulated strong ($s = 0.1$) and moderate ($s = 0.01$) soft sweep ($\nu = 8$) scenarios under the CEU-GIH model ($\tau = 1100$ generations before sampling). (Top row) Power at 1% (red lines) and 5% (purple lines) false positive rates (FPRs) to detect recent ancestral, convergent, and divergent soft sweeps from selection on standing genetic variation as a function of time at which selection of the favored haplotypes initiated ($t$), with FPR based on the distribution of maximum |SS-H12| across simulated neutral replicates. (Middle row) Box plots summarizing the distribution of SS-H12 values from windows of maximum |SS-H12| across strong sweep replicates, corresponding to each time point in the power curves, with dashed lines in each panel representing SS-H12 = 0. (Bottom row) Box plots summarizing the distribution of SS-H12 values across moderate sweep replicates. For convergent and divergent sweeps, $t < \tau$, while for ancestral sweeps, $t > \tau$. All replicate samples for the CEU-GIH model contain 99 simulated CEU individuals and 103 simulated GIH individuals, as in the 1000 Genomes Project dataset [Auton et al., 2015], and we performed 1000 replicates for each scenario.

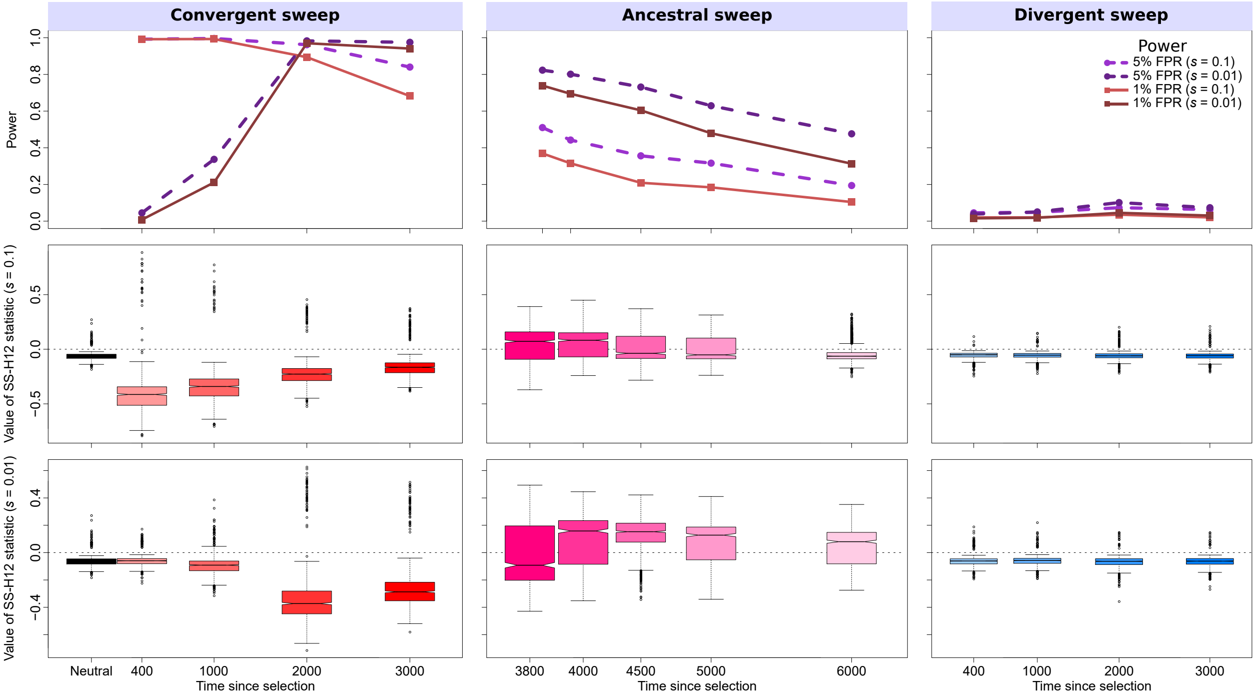


Figure S4: Properties of SS-H12 for simulated strong ($s = 0.1$) and moderate ($s = 0.01$) soft sweep ($\nu = 4$) scenarios under the CEU-YRI model ($\tau = 3740$ generations before sampling). (Top row) Power at 1% (red lines) and 5% (purple lines) false positive rates (FPRs) to detect recent ancestral, convergent, and divergent soft sweeps from selection on standing genetic variation as a function of time at which selection of the favored haplotypes initiated ($t$), with FPR based on the distribution of maximum |SS-H12| across simulated neutral replicates. (Middle row) Box plots summarizing the distribution of SS-H12 values from windows of maximum |SS-H12| across strong sweep replicates, corresponding to each time point in the power curves, with dashed lines in each panel representing SS-H12 = 0. (Bottom row) Box plots summarizing the distribution of SS-H12 values across moderate sweep replicates. For convergent and divergent sweeps, $t < \tau$, while for ancestral sweeps, $t > \tau$. All replicate samples for the CEU-YRI model contain 99 simulated CEU individuals and 108 simulated YRI individuals, as in the 1000 Genomes Project dataset [Auton et al., 2015], and we performed 1000 replicates for each scenario.

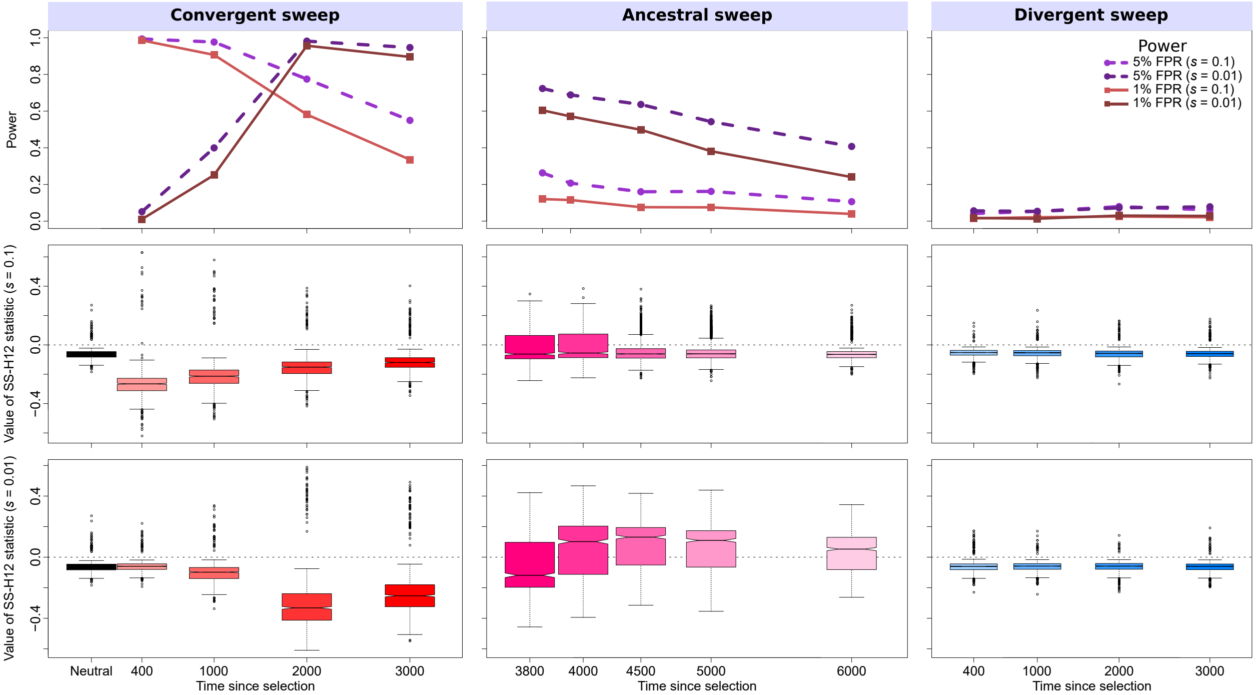


Figure S5: Properties of SS-H12 for simulated strong ($s = 0.1$) and moderate ($s = 0.01$) soft sweep ($\nu = 8$) scenarios under the CEU-YRI model ($\tau = 3740$ generations before sampling). (Top row) Power at 1% (red lines) and 5% (purple lines) false positive rates (FPRs) to detect recent ancestral, convergent, and divergent soft sweeps from selection on standing genetic variation as a function of time at which selection of the favored haplotypes initiated ($t$), with FPR based on the distribution of maximum |SS-H12| across simulated neutral replicates. (Middle row) Box plots summarizing the distribution of SS-H12 values from windows of maximum |SS-H12| across strong sweep replicates, corresponding to each time point in the power curves, with dashed lines in each panel representing SS-H12 = 0. (Bottom row) Box plots summarizing the distribution of SS-H12 values across moderate sweep replicates. For convergent and divergent sweeps, $t < \tau$, while for ancestral sweeps, $t > \tau$. All replicate samples for the CEU-YRI model contain 99 simulated CEU individuals and 108 simulated YRI individuals, as in the 1000 Genomes Project dataset [Auton et al., 2015], and we performed 1000 replicates for each scenario.

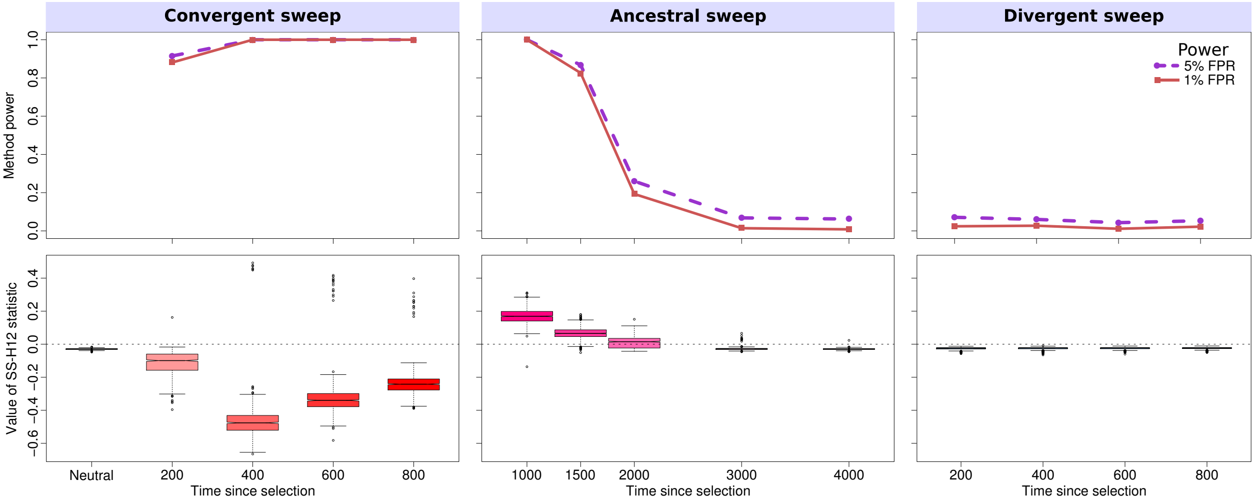


Figure S6: Properties of SS-H12 for simulated hard sweep scenarios for samples drawn from $K = 2$ equally-sized populations in which $\tau = 1000$ generations before sampling, for a constant demographic history in which $N = 10^4$ for the duration of the simulation. (Top) Power at 1 and 5% false positive rates (FPRs) to detect recent ancestral, convergent, and divergent hard sweeps (see Figure 1) as a function of time at which selection initiated, with false positive rate based on the distribution of maximum |SS-H12| across simulated neutral replicates. (Middle row) Box plots summarizing the distribution of SS-H12 values from windows of maximum |SS-H12| for each replicate, corresponding to each point in the power curve, with dashed lines in each panel representing SS-H12 $= 0$. Convergent and divergent sweeps occur after this time (200-800 generations before sampling), while ancestral sweeps occur before this time (1100-4000 generations before sampling). All sweeps are strong ($s = 0.1$) for a sample of $n = 100$ diploid individuals per population, with 1000 replicates performed for each scenario.

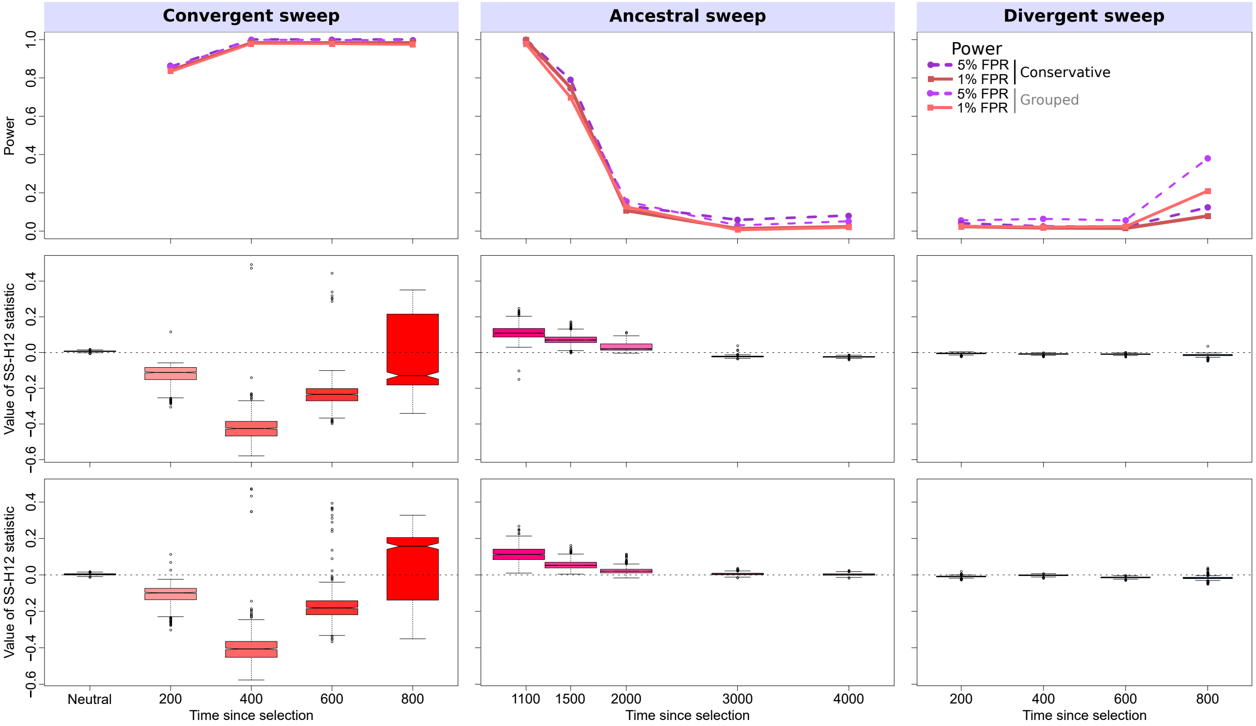


Figure S7: Properties of SS-H12 for simulated hard sweep scenarios for samples drawn from $K = 3$ equally-sized populations in which $\tau_1 = 1000$ and $\tau_2 = 750$ generations before sampling. (Top) Power at 1 and 5% false positive rates (FPRs) to detect recent ancestral, convergent, and divergent hard sweeps (see Figure 1) as a function of time at which selection initiated, with false positive rate based on the distribution of maximum |SS-H12| across simulated neutral replicates. (Middle and bottom rows) Box plots summarizing the distribution of SS-H12 values from windows of maximum |SS-H12| for each replicate, corresponding to each point in the power curve for conservative (middle row) and grouped (bottom row) approaches, with dashed lines in each panel representing SS-H12 = 0. Convergent and divergent sweeps occur after this time (200-800 generations before sampling), while ancestral sweeps occur before this time (1100-4000 generations before sampling). All sweeps are strong ($s = 0.1$) for a sample of $n = 100$ diploid individuals per population, with 1000 replicates performed for each scenario.

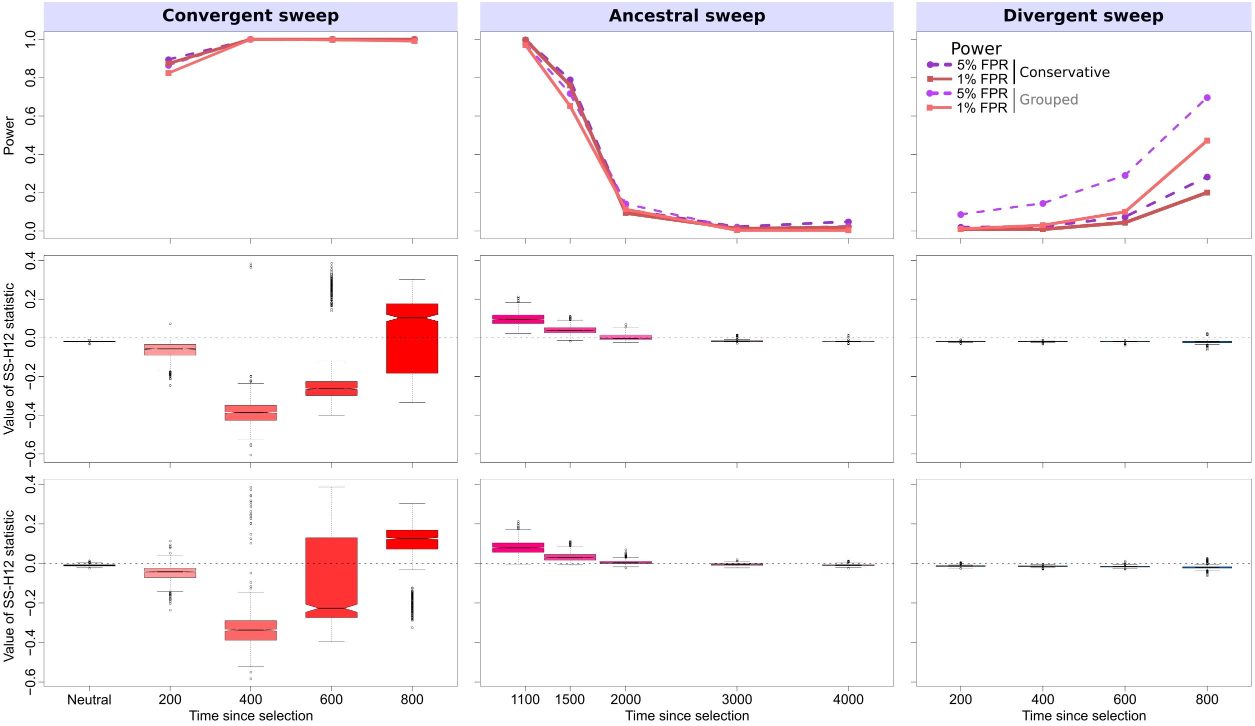


Figure S8: Properties of SS-H12 for simulated hard sweep scenarios for samples drawn from $K = 4$ equally-sized populations in which $\tau_1 = 1000$, $\tau_2 = 750$, and $\tau_3 = 500$ generations before sampling. (Top row) Power at 1 and 5% false positive rates (FPRs) to detect recent ancestral, convergent, and divergent hard sweeps (see Figure 1) as a function of time at which selection initiated, with false positive rate based on the distribution of maximum |SS-H12| across simulated neutral replicates. (Middle and bottom rows) Box plots summarizing the distribution of SS-H12 values from windows of maximum |SS-H12| for each replicate, corresponding to each point in the power curve for conservative (middle row) and grouped (bottom row) approaches, with dashed lines in each panel representing SS-H12 = 0. Convergent and divergent sweeps occur after this time (200-800 generations before sampling), while ancestral sweeps occur before this time (1100-4000 generations before sampling). All sweeps are strong ($s = 0.1$) for a sample of $n = 100$ diploid individuals per population, with 1000 replicates performed for each scenario.

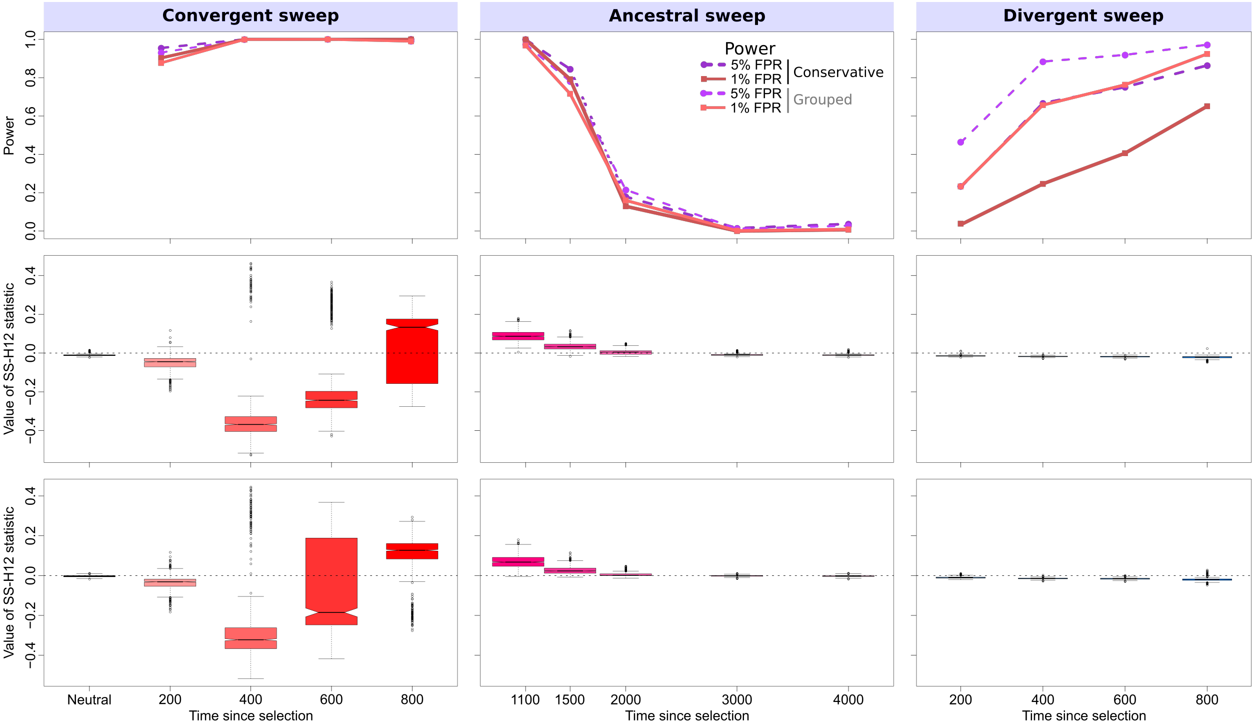


Figure S9: Properties of SS-H12 for simulated hard sweep scenarios for samples drawn from $K = 5$ equally-sized populations in which $\tau_1 = 1000$, $\tau_2 = 750$, $\tau_3 = 500$, and $\tau_4 = 250$ generations before sampling. (Top row) Power at 1 and 5% false positive rates (FPRs) to detect recent ancestral, convergent, and divergent hard sweeps (see Figure 1) as a function of time at which selection initiated, with false positive rate based on the distribution of maximum |SS-H12| across simulated neutral replicates. (Middle and bottom rows) Box plots summarizing the distribution of SS-H12 values from windows of maximum |SS-H12| for each replicate, corresponding to each point in the power curve for conservative (middle row) and grouped (bottom row) approaches, with dashed lines in each panel representing SS-H12 = 0. Convergent and divergent sweeps occur after this time (200-800 generations before sampling), while ancestral sweeps occur before this time (1100-4000 generations before sampling). All sweeps are strong ($s = 0.1$) for a sample of $n = 100$ diploid individuals per population, with 1000 replicates performed for each scenario.

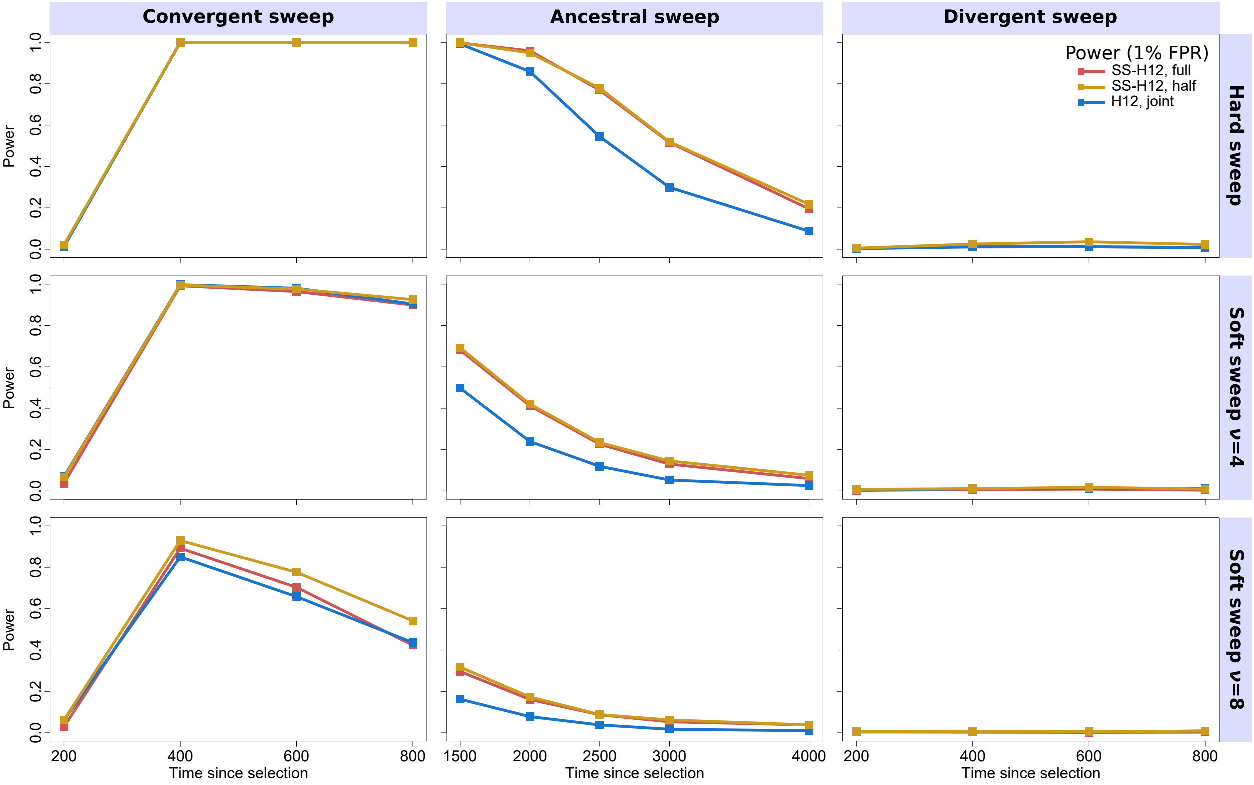


Figure S10: Powers of SS-H12 and H12 at a 1% false positive rate to detect recent ancestral, convergent, and divergent simulated strong ($s = 0.1$) selective sweeps under the CEU-GIH model ($\tau = 1100$ generations before sampling) for hard (top) and soft ($\nu = 4$, middle; $\nu = 8$, bottom) sweeps as a function of time at which positive selection of the favored allele initiated ($t$), with FPR based on the distribution of maximum |SS-H12| or H12 across simulated neutral replicates. "SS-H12, full" refers to analyses in which we generated full-size pooled samples consisting of 99 simulated CEU individuals and 103 simulated GIH individuals, whereas "SS-H12, half" used only half of the alleles in each sample. "H12, joint" analyses used the full sample size for both populations. For convergent and divergent sweeps, $t < \tau$, while for ancestral sweeps, $t > \tau$. We performed 1000 replicates for each scenario.

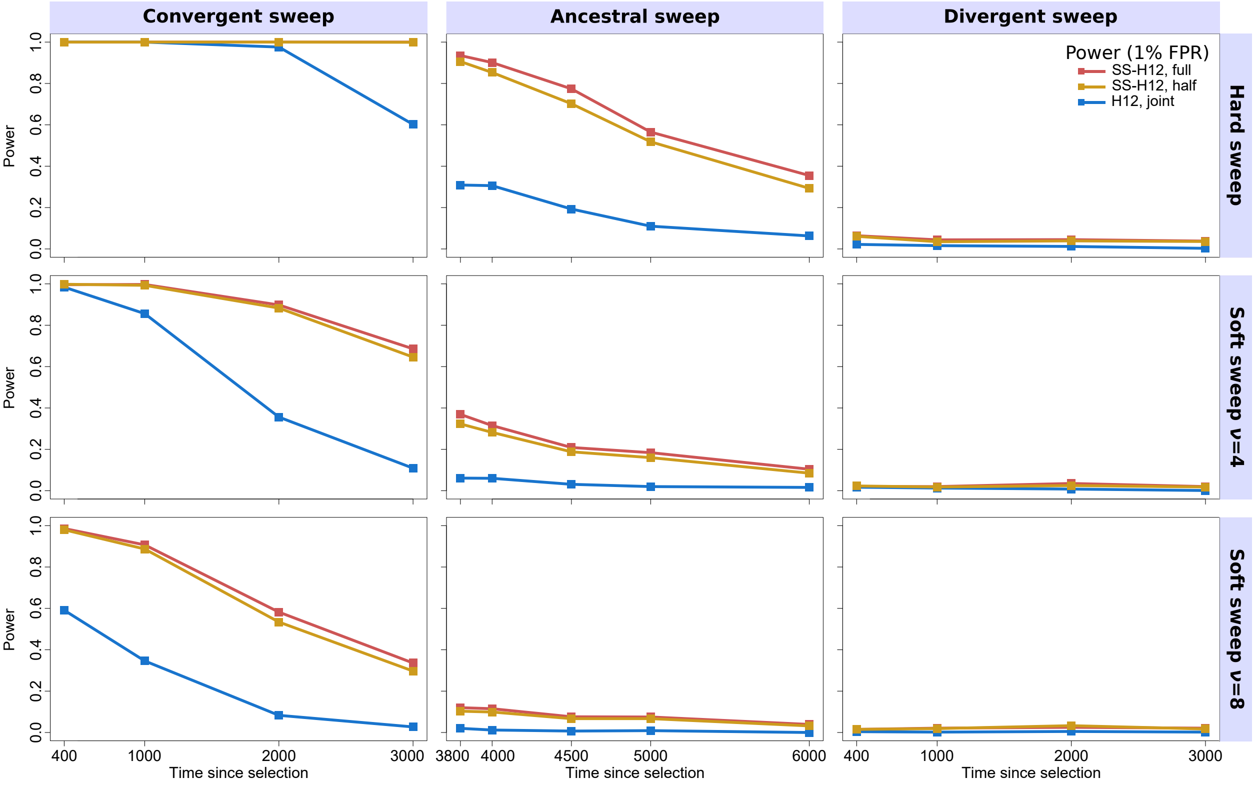


Figure S11: Powers of SS-H12 and H12 at a 1% false positive rate to detect recent ancestral, convergent, and divergent simulated strong ($s = 0.1$) selective sweeps under the CEU-YRI model ($\tau = 3740$ generations before sampling) for hard (top) and soft ($\nu = 4$, middle; $\nu = 8$, bottom) sweeps as a function of time at which positive selection of the favored allele initiated ($t$), with FPR based on the distribution of maximum |SS-H12| or H12 across simulated neutral replicates. "SS-H12, full" refers to analyses in which we generated full-size pooled samples consisting of 99 simulated CEU individuals and 108 simulated GIH individuals, while "SS-H12, half" used only half of the alleles in each sample. "H12, joint" analyses used the full sample size for both populations. For convergent and divergent sweeps, $t < \tau$, while for ancestral sweeps, $t > \tau$. We performed 1000 replicates for each scenario.

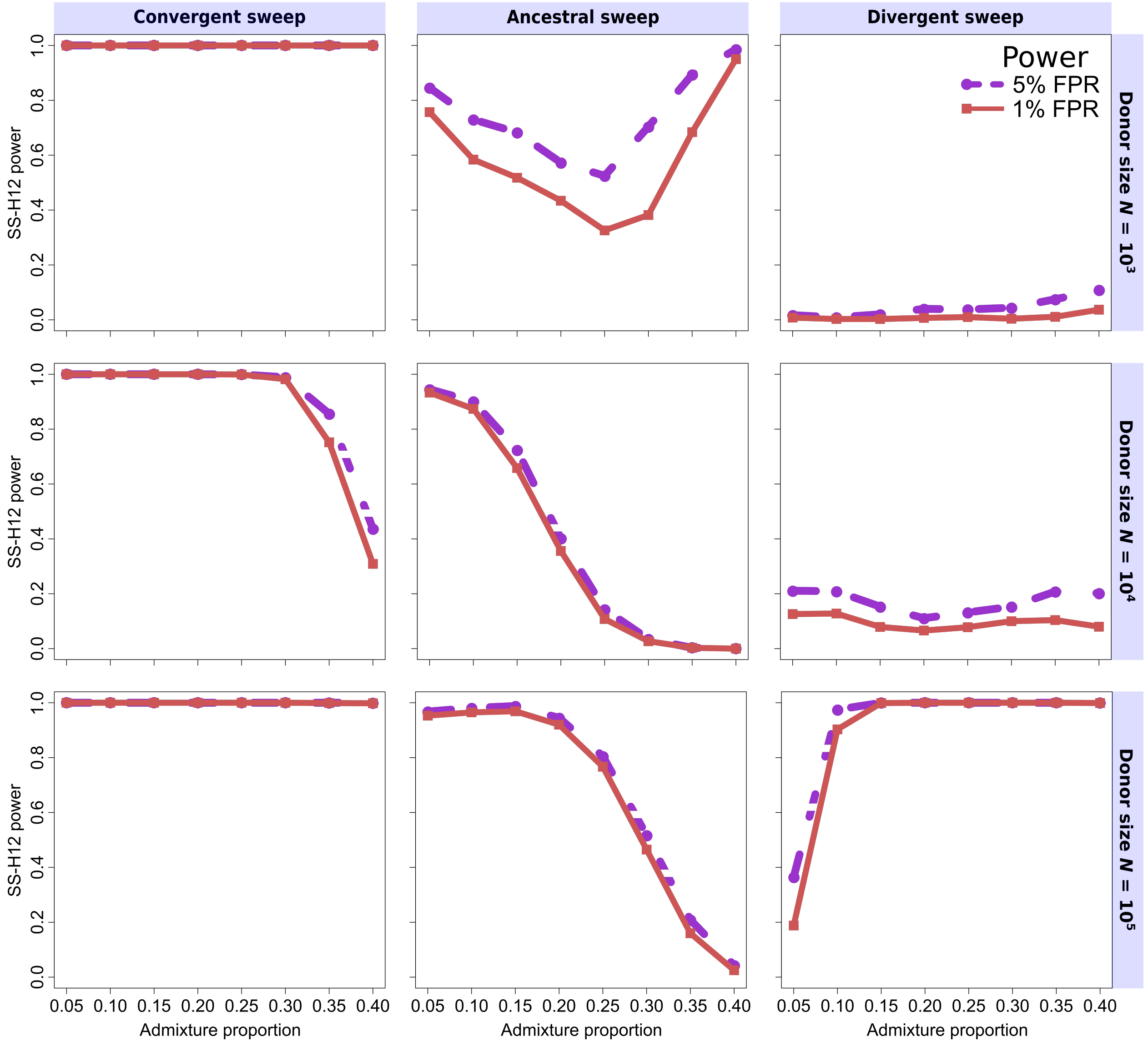


Figure S12: Power at 1 and 5% false positive rates (FPRs) to detect recent strong ($s = 0.1$) ancestral, convergent, and divergent hard sweeps (see Figure 1) as a function of admixture proportion, with false positive rate based on the distribution of maximum |SS-H12| across simulated neutral replicates under a given admixture scenario. Data here are simulated as in Figure 4, with scenarios of admixture proportion 0.2, 0.3, and 0.4 identical to Figure 4.

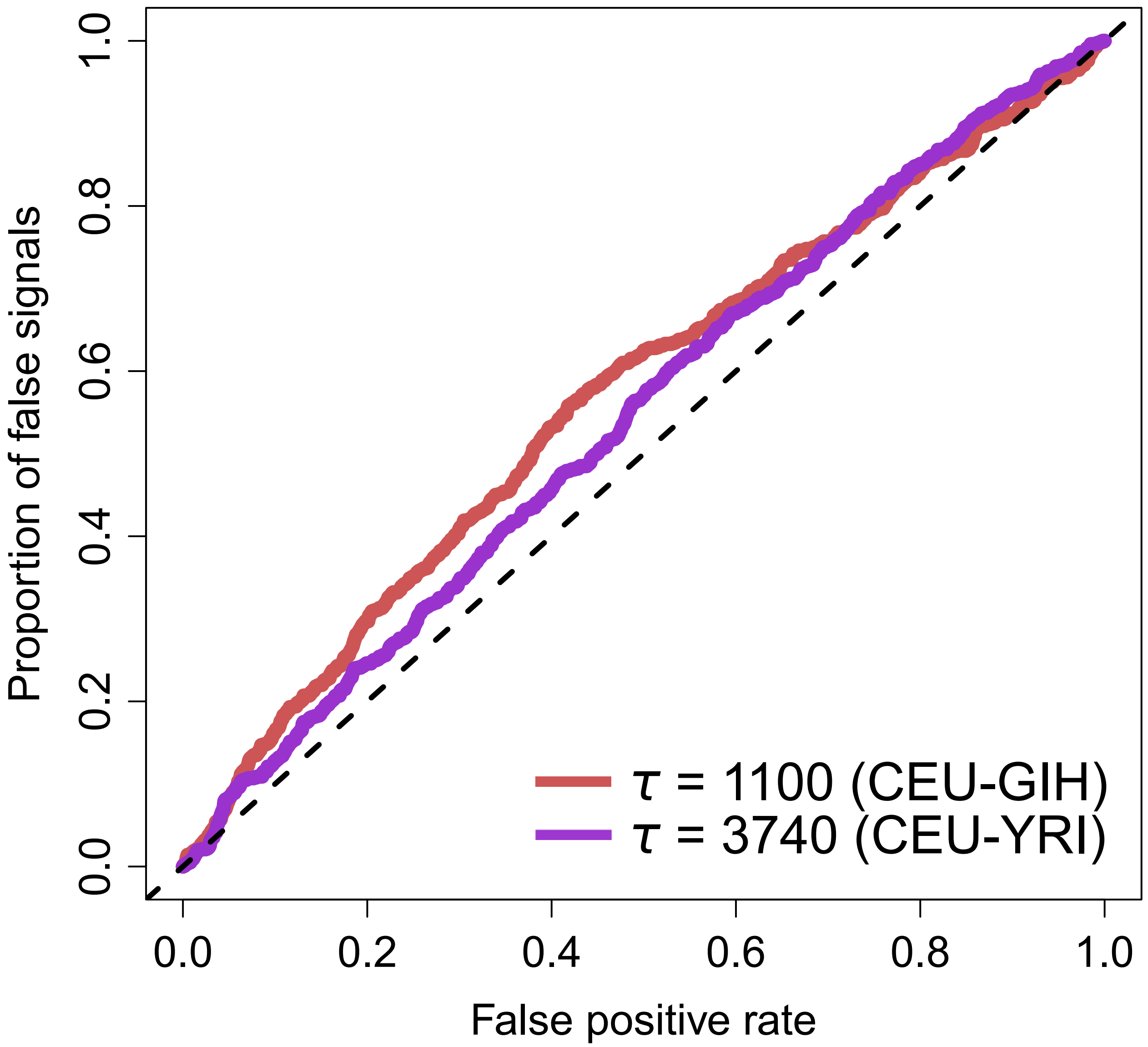


Figure S13: Proportion of false signals generated by background selection as a function of false positive rate based on neutrality for SS-H12 under CEU-GIH (red) or CEU-YRI (purple) demographic models. Neutral replicates used here are identical to those used across Figures 2, 3, and S2-S5. Background selection simulations, totaling 1000 replicates, were generated according to the procedure described in the *Materials and Methods*.

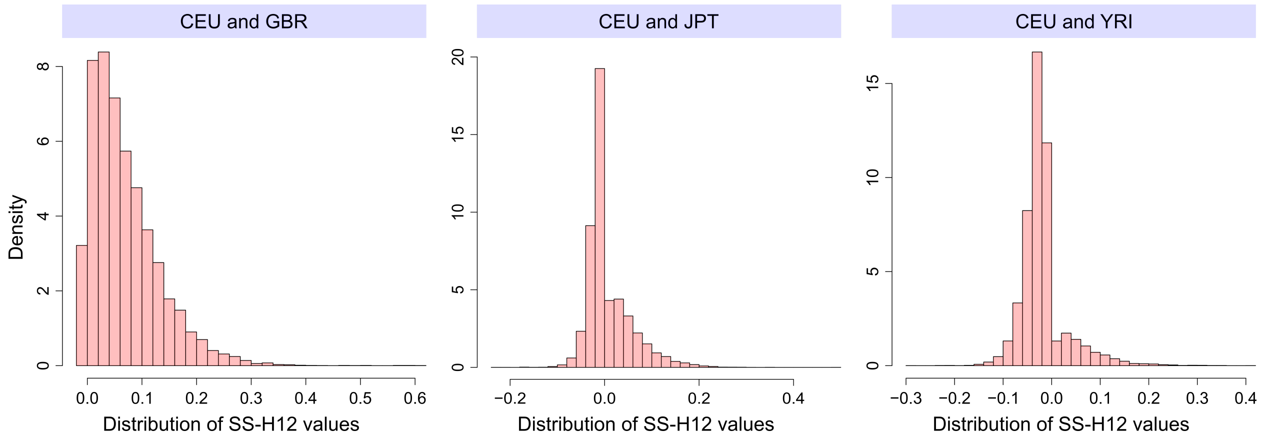


Figure S14: Empirical distributions of SS-H12 values assigned to each protein- and mRNA-coding gene based on the analysis of individuals from the 1000 Genomes Project Consortium [Auton et al., 2015] for three population comparisons with CEU. (Left) Distribution of SS-H12 values between the closely-related CEU and GBR populations of European descent. (Middle) Distribution of SS-H12 values between CEU and the East Asian JPT populations, representing a pair that diverged after the out-of-Africa event. (Right) Distribution of SS-H12 values between CEU and the sub-Saharan African YRI populations, representing a distantly-related population pair. SS-H12 is assigned to each gene from the window of maximum |SS-H12| falling between the transcription start and stop of the gene.

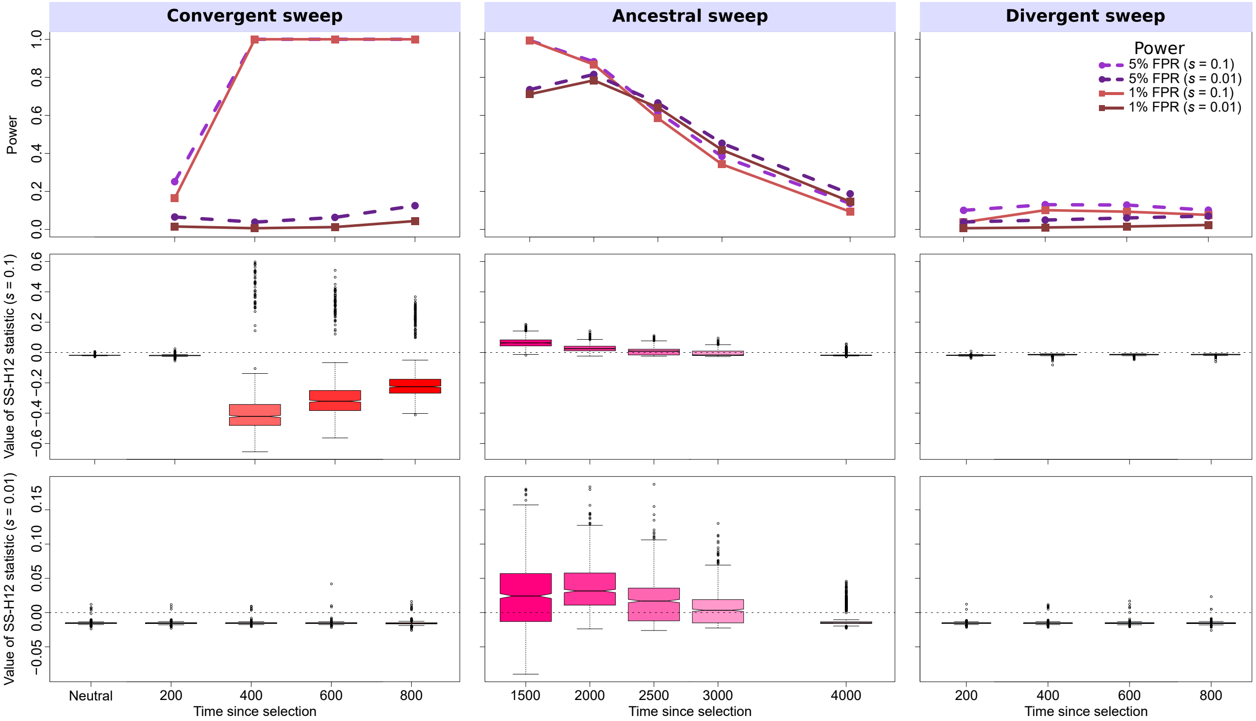


Figure S15: Properties of SS-G123 for simulated strong ($s = 0.1$) and moderate ($s = 0.01$) hard sweep scenarios under the CEU-GIH model ($\tau = 1100$ generations before sampling). (Top row) Power at 1% (red lines) and 5% (purple lines) false positive rates (FPRs) to detect recent ancestral, convergent, and divergent hard sweeps (see Figure 1) as a function of time at which positive selection of the favored allele initiated ($t$), with FPR based on the distribution of maximum |SS-G123| across simulated neutral replicates. (Middle row) Box plots summarizing the distribution of SS-G123 values from windows of maximum |SS-G123| across strong sweep replicates, corresponding to each time point in the power curves, with dashed lines in each panel representing SS-G123 = 0. (Bottom row) Box plots summarizing the distribution of SS-G123 values across moderate sweep replicates. For convergent and divergent sweeps, $t < \tau$, while for ancestral sweeps, $t > \tau$. Simulated replicates are identical to those in Figure 2, but with each individual's two haplotypes merged into their multilocus genotypes.

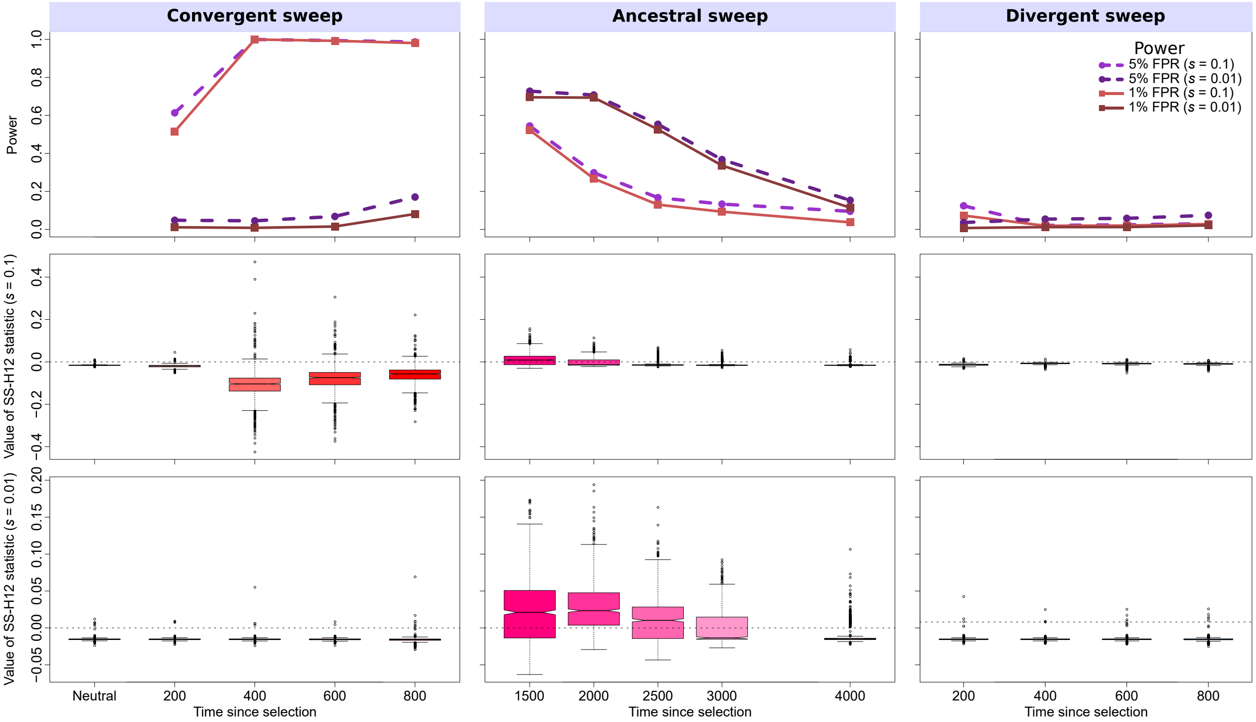


Figure S16: Properties of SS-G123 for simulated strong ($s = 0.1$) and moderate ($s = 0.01$) soft sweep ($\nu = 4$) scenarios under the CEU-GIH model ($\tau = 1100$ generations before sampling). (Top row) Power at 1% (red lines) and 5% (purple lines) false positive rates (FPRs) to detect recent ancestral, convergent, and divergent soft sweeps from selection on standing genetic variation as a function of time at which selection of the favored haplotypes initiated ($t$), with FPR based on the distribution of maximum |SS-G123| across simulated neutral replicates. (Middle row) Box plots summarizing the distribution of SS-G123 values from windows of maximum |SS-G123| across strong sweep replicates, corresponding to each time point in the power curves, with dashed lines in each panel representing SS-G123 = 0. (Bottom row) Box plots summarizing the distribution of SS-G123 values across moderate sweep replicates. For convergent and divergent sweeps, $t < \tau$, while for ancestral sweeps, $t > \tau$. Simulated replicates are identical to those in Figure S2, but with each individual's two haplotypes merged into their multilocus genotypes.

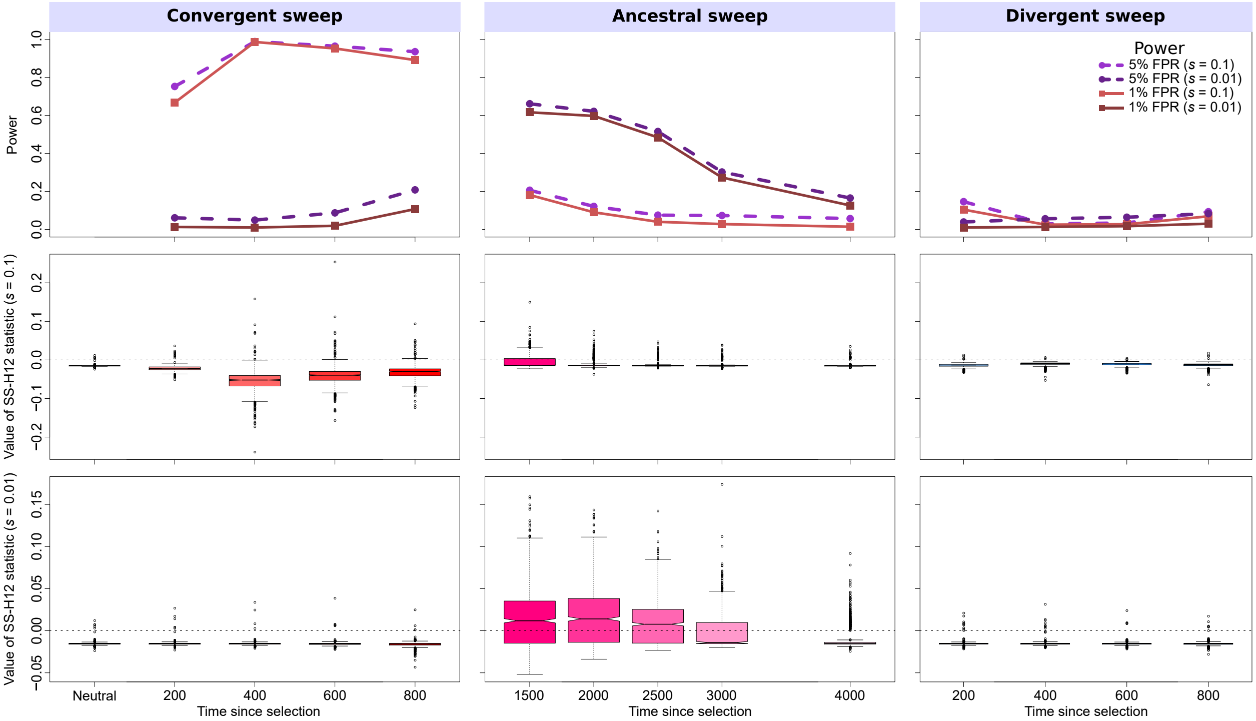


Figure S17: Properties of SS-G123 for simulated strong ($s = 0.1$) and moderate ($s = 0.01$) soft sweep ($\nu = 8$) scenarios under the CEU-GIH model ($\tau = 1100$ generations before sampling). (Top row) Power at 1% (red lines) and 5% (purple lines) false positive rates (FPRs) to detect recent ancestral, convergent, and divergent soft sweeps from selection on standing genetic variation as a function of time at which selection of the favored haplotypes initiated ($t$), with FPR based on the distribution of maximum |SS-G123| across simulated neutral replicates. (Middle row) Box plots summarizing the distribution of SS-G123 values from windows of maximum |SS-G123| across strong sweep replicates, corresponding to each time point in the power curves, with dashed lines in each panel representing SS-G123 = 0. (Bottom row) Box plots summarizing the distribution of SS-G123 values across moderate sweep replicates. For convergent and divergent sweeps, $t < \tau$, while for ancestral sweeps, $t > \tau$. Simulated replicates are identical to those in Figure S3, but with each individual's two haplotypes merged into their multilocus genotypes.

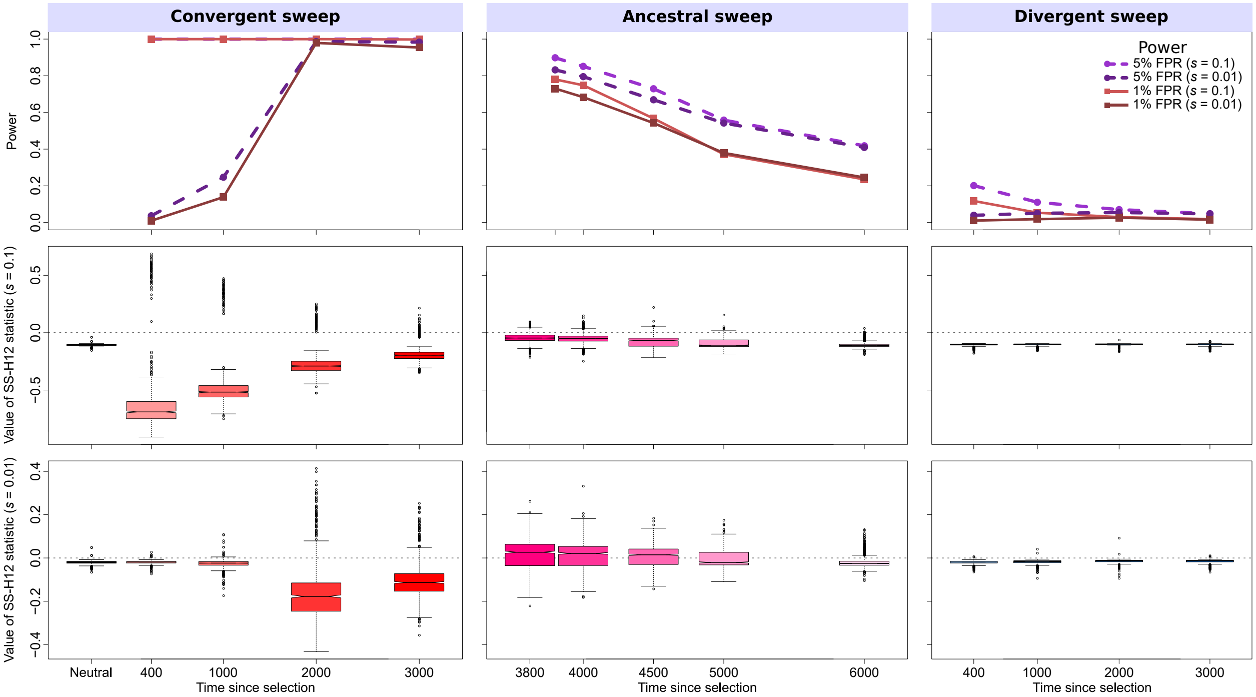


Figure S18: Properties of SS-G123 for simulated strong ($s = 0.1$) and moderate ($s = 0.01$) hard sweep scenarios under the CEU-YRI model ($\tau = 3740$ generations before sampling). (Top row) Power at 1% (red lines) and 5% (purple lines) false positive rates (FPRs) to detect recent ancestral, convergent, and divergent hard sweeps (see Figure 1) as a function of time at which positive selection of the favored allele initiated ($t$), with FPR based on the distribution of maximum |SS-G123| across simulated neutral replicates. (Middle row) Box plots summarizing the distribution of SS-G123 values from windows of maximum |SS-G123| across strong sweep replicates, corresponding to each time point in the power curves, with dashed lines in each panel representing SS-G123 = 0. (Bottom row) Box plots summarizing the distribution of SS-G123 values across moderate sweep replicates. For convergent and divergent sweeps, $t < \tau$, while for ancestral sweeps, $t > \tau$. Simulated replicates are identical to those in Figure 3, but with each individual's two haplotypes merged into their multilocus genotypes.

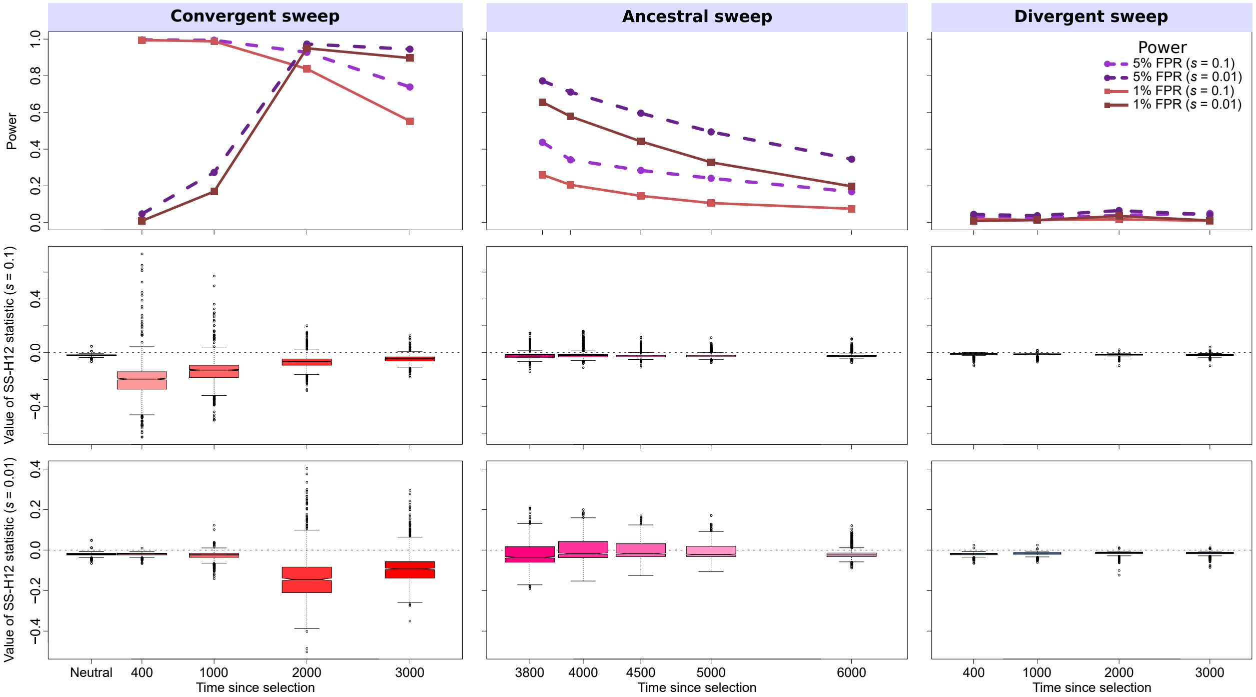


Figure S19: Properties of SS-G123 for simulated strong ($s = 0.1$) and moderate ($s = 0.01$) soft sweep ($\nu = 4$) scenarios under the CEU-YRI model ($\tau = 3740$ generations before sampling). (Top row) Power at 1% (red lines) and 5% (purple lines) false positive rates (FPRs) to detect recent ancestral, convergent, and divergent soft sweeps from selection on standing genetic variation as a function of time at which selection of the favored haplotypes initiated ($t$), with FPR based on the distribution of maximum |SS-G123| across simulated neutral replicates. (Middle row) Box plots summarizing the distribution of SS-G123 values from windows of maximum |SS-G123| across strong sweep replicates, corresponding to each time point in the power curves, with dashed lines in each panel representing SS-G123 = 0. (Bottom row) Box plots summarizing the distribution of SS-G123 values across moderate sweep replicates. For convergent and divergent sweeps, $t < \tau$, while for ancestral sweeps, $t > \tau$. Simulated replicates are identical to those in Figure S4, but with each individual's two haplotypes merged into their multilocus genotypes.

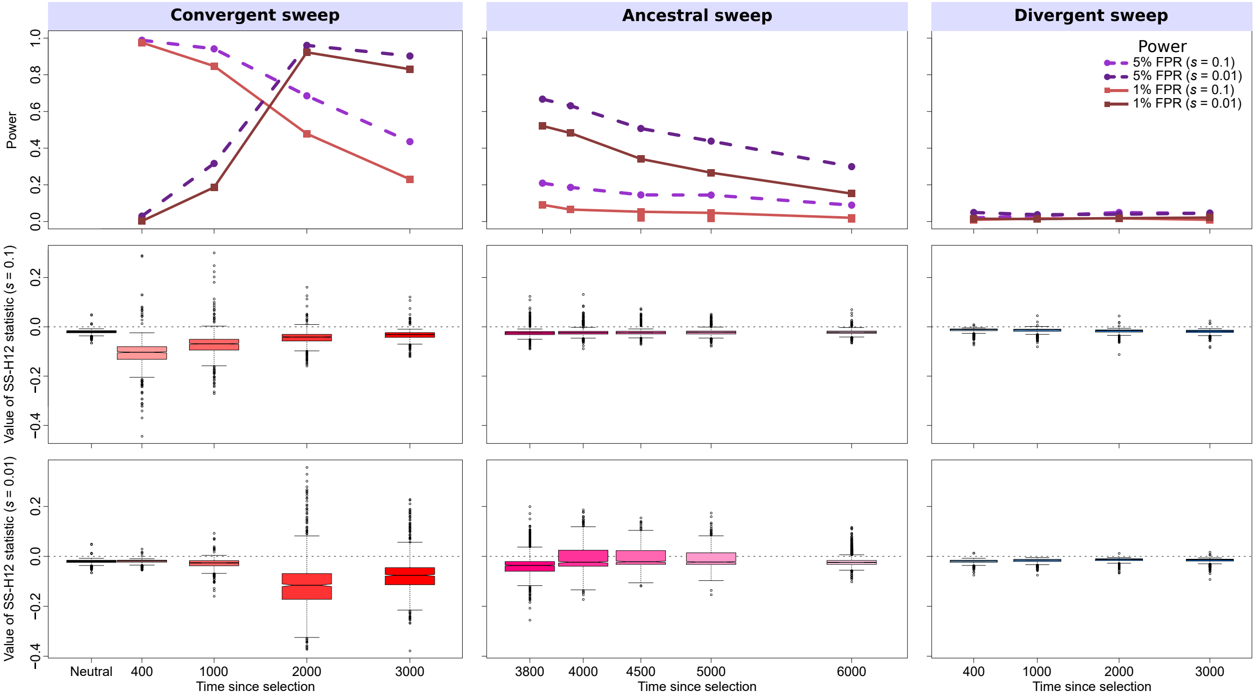


Figure S20: Properties of SS-G123 for simulated strong ($s = 0.1$) and moderate ($s = 0.01$) soft sweep ($\nu = 8$) scenarios under the CEU-YRI model ($\tau = 3740$ generations before sampling). (Top row) Power at 1% (red lines) and 5% (purple lines) false positive rates (FPRs) to detect recent ancestral, convergent, and divergent soft sweeps from selection on standing genetic variation as a function of time at which selection of the favored haplotypes initiated ($t$), with FPR based on the distribution of maximum |SS-G123| across simulated neutral replicates. (Middle row) Box plots summarizing the distribution of SS-G123 values from windows of maximum |SS-G123| across strong sweep replicates, corresponding to each time point in the power curves, with dashed lines in each panel representing SS-G123 = 0. (Bottom row) Box plots summarizing the distribution of SS-G123 values across moderate sweep replicates. For convergent and divergent sweeps, $t < \tau$, while for ancestral sweeps, $t > \tau$. Simulated replicates are identical to those in Figure S5, but with each individual's two haplotypes merged into their multilocus genotypes.

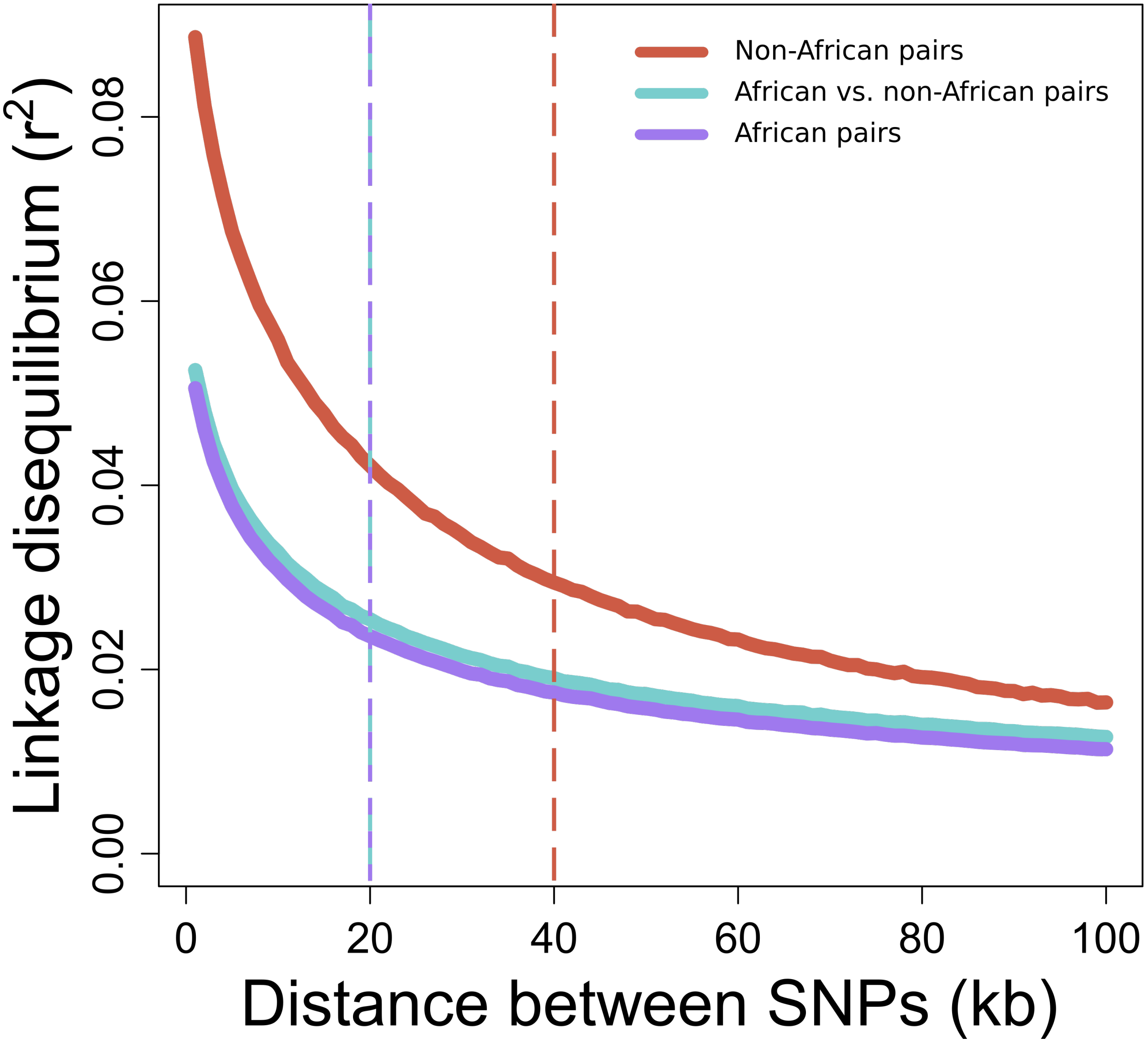


Figure S21: Mean decay of pairwise linkage disequilibrium (LD) measured by $r^2$ over one to 100 kb intervals downstream of each SNP for pooled population pair samples from the 1000 Genomes Project dataset [Auton et al., 2015]. We used the pooled CEU-JPT sample as representative of LD between non-African population pairs (red), the pooled CEU-YRI population as representative of LD between African and non-African population pairs (cyan), and the pooled LWK-YRI population as representative of the LD between African population pairs (purple). All samples are identical to those analyzed in the *Results* section.
